# Strand Orientation Bias Detector (SOBDetector) to remove FFPE sequencing artifacts

**DOI:** 10.1101/386417

**Authors:** Miklos Diossy, Zsofia Sztupinszki, Marcin Krzystanek, Judit Borcsok, Aron C. Eklund, István Csabai, Anders Gorm Pedersen, Zoltan Szallasi

## Abstract

Formalin-fixed paraffin-embedded (FFPE) tissue, the most common tissue specimen stored in clinical practice, presents challenges in the analysis due to formalin-induced artifacts. Here we present SOBDetector, that is designed to remove these artifacts from a list of variants, which were extracted from next-generation sequencing data, using the posterior predictive distribution of a Bayesian logistic regression model trained on TCGA whole exomes. Based on a concordance analysis, SOBDetector outperforms the currently publicly available FFPE-filtration techniques (Accuracy: 0.9 ± 0.015, AUC = 0.96). SOBDetector is implemented in Java 1.8 and is freely available online.

## INTRODUCTION

Because of their wide availability, formalin-fixed paraffin-embedded tissues (FFPE) are valuable resources of translational and clinical research. Next generation sequencing of DNA extracted from such samples is often performed for exploratory or validation purposes. The chemical processes fixating the tissue material, however, introduce a significant amount of DNA damage that manifests as misread bases during sequencing, which makes the analysis of such samples a challenging task. The most prominent of these is the deamination of cytosines, which introduces C→T/G→A mutations. (1–3) Since formalin likely affects only one of the strands (e.g. a C|G pair becomes T|G), a paired-end next generation sequencing approach can be utilized to remove such sequencing artifacts (Supplementary Figure 1). When considering not just the number of reads that support the alternative allele, but also their relative orientation (Forward-Reverse:FR or Reverse-Forward:RF), the FFPE artifacts will likely have a strand/read orientation bias towards one of the directions, while true mutations, provided their loci have sufficient coverage, should have approximately the same amount of FR and RF reads.

## MATERIAL AND METHODS

### Sample collection

The cohort of patients (N = 47) with matched FF and FFPE-derived variant files was determined using the GDC Data Portal (https://portal.gdc.cancer.gov/). Since most of these donors had multiple FF and/or FFPE tumors, and in some cases even multiple germline samples, altogether 235 somatic VCFs were available (Supplementary Table 1). For the initial investigations all the variants (N = 163,000) from these files had been merged into a single data table. Since these mutations were all identified using the default filters of MuTect2 (nightly-2016-02-25-gf39d340 built, which was closest to GATK 3.7(4)), additional common filters were defined over them: Within both the tumor and normal samples the sequencing depths were required to be at least 20, and the minimum of the variant allele-frequencies was set to 0.05, which has reduced the total number of variants to N = 120,948.

In order to create a trainable dataset, the likely artifacts and likely novel mutations were collected into a subset of this table. Variants that were present in at least one FFPE sample, and at least 2 FF samples (derived from different tumor BAMs, but of the same tumor and the same donor) had been marked as likely novel mutations. Altogether 8672 unique variant positions met this criterion, which were present in ⟨*NFF*⟩=4.84 fresh frozen and ⟨*NFFPE*⟩=1.86 FFPE specimens on average, resulting in 56,108 observations. Variants, that were present in only one FFPE sample, and none of the FF counterparts of the same tumor and the same donor had been marked as likely FFPE artifacts. Altogether 7599 variant positions met this condition, resulting in 7599 unique observations (since ⟨*NFF*⟩=0 and ⟨*NFFPE*⟩=1.00). In order to minimalize the chance that genuine mutations are categorized as likely artifacts due to intratumor heterogeneity, other mutations, such as those that were only present in 1 FF and 1 FFPE, or in two FFPE samples, were categorized as unknown variants.

### Training attributes, and separability

SOBDetector was designed to require only the locus (chromosome and position), the reference allele and the alternate allele of the variant, everything else it collects from the alignment files, or it calculates on its own. Originally the following 11 features had been considered as predictors: TUMOR.depth, i.e. the coverage at the given position in the tumor BAM, NORMAL.depth, i.e. the coverage at the given position in the normal BAM, TUMOR.ALT, i.e. the number of reads (out of TUMOR.depth) that support the alternative allele in the tumor, TUMOR.AF, the tumor allelic frequency at the given position, and SOB score, i.e. the strand orientation bias score of the variant, plus the 6 non-redundant mutational directions. These have been collected using a 1 of K encoding strategy. This meant, that instead of a single column containing the direction of the mutation, 6 new columns have been defined, each corresponding to one of the directions. For a variant, only one of these attributes (the one that corresponded the REF>ALT direction) was set to 1, all the others were set to zero.

Since the distribution of these attributes (*x*_*i*_) were skewed and failed the test of normality, they had been log-transformed and standardized prior training, using the following formulas

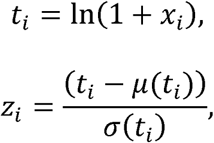

Where *µ*(.) and *σ*(.) are the mean and standard deviation functions respectively. Figure 2B shows the kernel-smoothed density approximations of these *z*_*i*_ transformed attributes.

In order to assess the feasibility of a linear separator, the principal components of the trainable dataset were analyzed. It was found, that the first seven principal components (PCs) were responsible for ~ 90% of the variance within the set, and while the first eigenvector was dominated by the non-directional attributes, from the second, each principal component had major contributions from at least one of the mutational directions. The visual inspection of the two dimensional projections of the data onto its principal components revealed that although the dataset could not be separated perfectly using a linear separator, a logistic regression model with a soft-threshold defined between the groups might work adequately (Supplementary Figure 6).

### The training model

Given the dataset 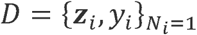, where *y*_*i*_ ∈ {0,1}, encodes whether a variant is a likely genuine mutation (y = 0) or an artifact (y = 1), and *z*_*i*_ ∈ ℝ^*d*^, are the log-transformed, and standardized features defined above. The aim is to calculate the joint posterior distribution of the binary logistic regression model (Supplementary Figure 7):

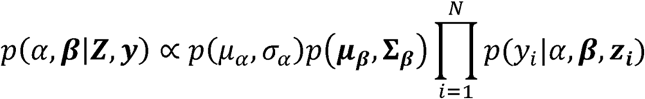

where α is the intercept and **β** is a vector containing the weights of the linear separator. *µ* and *σ* indicate the location and scale parameters of the initial (prior) distributions of the intercept and of the weights. The lhs is the posterior distribution given the set of the standardized and log-transformed features and their assumed classes (**Z** and **y** respectively), while the rhs contains a product of the prior on α and **β**, multiplied by the likelihoods that a certain combination of the bias, the weights and the features result in a particular class y. In case of the logistic regression model, this likelihood is a Bernoulli density:

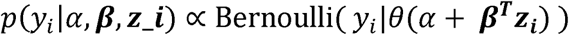

where θ(s) represents the logistic function:

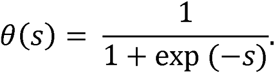

The priors defined over the weights act as regularizers, and they can help in feature selection, and to avoid the overfitting of the data. We have decided to implement a horseshoe prior, which is a heavy tailed mixture distribution, belonging into the family of multivariate scale mixture normals. Its generative process is the following:

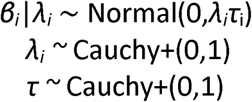

Although this regularizer was originally designed for sparse datasets, it has similar properties to a LASSO regularizer in classical machine learning models, hence it can be effectively used for feature selection in Bayesian models. However, despite of the LASSO’s unit constraint on all of the weights, the horseshoe prior operates with both a global, and a set of feature-specific, so called local shrinkage parameters, giving the model a refined feature selection property. The code of the model was written in standard stan format, and trained from R, using rstan. For training a model, 6 random walkers had been initiated, each set to 2000 iterations, with warm-up lengths equal to five hundred iterations(5). Since in the final trainable set the likely novel mutation group had almost twice as many variants as the likely artifact group, the former was randomly under sampled to the same number of the likely artifacts. 90% of these observations had been collected into a training set (N = 32444), while the remaining (N = 5726) variants had been set aside for evaluation purposes.

The marginal posteriors of each of the attributes had close to 1 Rhat scores, indicating that the walkers have converged to the true posteriors. After training a model using the 11 features (Supplementary Figure 8 and Supplementary Table 2), we have concluded that the mutational directions and the depths in the normal sample had no predictive powers, hence they were removed from the list of predictors. A new, final model consisting only the 4 remaining features (TUMOR.ALT, TUMOR.AF, TUMOR.depth, and SOB scores) was trained on the same training set, and the predicted probabilities of the variants within the test set (evaluation set) were calculated using the following formula:

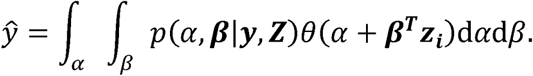

### Evaluating the model

The posterior predictions of the model were estimated using the unbiased evaluation dataset. As a function of a discrete set of considerable thresholds, two curves: the

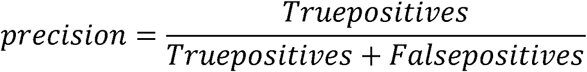

and the

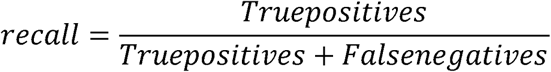

were calculated, along with the harmonic means of the two, called the F1-score (Figure 3B):

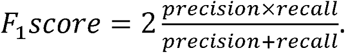

At the maximum of the F1-curve an evaluation set specific threshold had been determined, and the accuracy of the predictions have been measured (=0.904). The credibility interval of the model was estimated using a binomial test.

#### Sample-specific thresholds

Since the SOB scores had the largest predictive power among the training features, not counting some minor fluctuations, the mean of the SOB scores of the variants that were sorted according to their prediction scores, increased monotonically in all of the cases. This allowed us to define a dynamic threshold at the point, where the moving average of the SOB scores remined consistently higher than 0.8. The moving average (s_i_) is defined as:

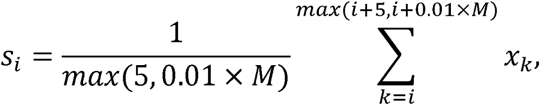

where *M* is the total number of variants in the sample, and *x*_*i*_ is the SOB score of the variant at position *i*. This means, that for the variant at the sorted coordinate *i*, the tool considers the following at least 5, or maximum 1% of the “total number of mutations present in the sample” SOB scores and calculates their average. When these averages reach above 0.8, the threshold gets defined.

### Concordance Analysis

A concordance analysis was conducted using the breast and colon cancer cohorts. These two were selected, because based on our initial investigations, they were affected differently by the fixation process. While the breast cancer formalin-fixed samples had significantly more mutations than their fresh frozen counterparts, indicating that they were enriched by FFPE artifacts, such difference was not observed among the colon cancer VCFs. The colon cancer cohort in general had more shared mutations between the corresponding FF and FFPE pairs, and the FFPE damage was not as severe as it was in the breast cancer case. In this part of the analysis the performances of 10 different mutation calling pipelines had been compared. The major pipelines were the following: GATK 3.7 MuTect2 (The original TCGA VCFs), GATK 4.1.1 Mutect2 with and without a strand orientation bias filtering (FilteryByOrientationBias) step, and GATK 4.1.2 with and without a strand orientation bias (LearnReadOrientationModel) step. While using all of the mutation calling tools, the official GATK documentations, recommendations, and best practices guidelines were followed, including the creation of a panel of normal (PON) using the normal samples of the 47 patients. In addition, all of these pipelines had been complemented with an additional SOBDetector-filtration step, resulting in 10 different sets of variants for each tumor BAMs. The following measures were used: the concordance between variant file1 (f1) and variant file2 (f2) was defined as the ratio of shared mutations by the two sets, divided by the total number of mutations present in file1:

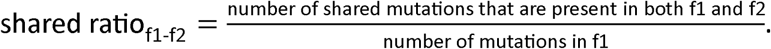

For FFPE samples file1 was the FFPE variant file, file2 was a corresponding FF sample of the same patient. For FF samples both file1 and file2 were fresh frozen specimens of the same donor. An increase in the shared ratio between the pre and post-filtering states indicates, that the concordance between the two samples have increased, and the filtering process was able to remove more false positive mutations than true positive, genuine variants.

The ratio of remaining mutations is the number of remaining mutations that are shared between f1 and f2 after filtering, divided by the number of mutations that that was shared between them prior filtering.

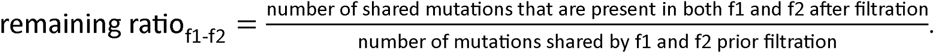

The pre-filtering state of the remaining ratio is always 1 (i.e. 100%). A post filtering state less than 1 informs us, that the filtering process has affected approximately (1-remaining_ratio)×100 percent of the true mutations calls as well.

Ideally, both of these ratios should be high, meaning that a filtration step removes most of the artifacts without removing any of the likely genuine (and hence shared) mutations. In addition, the ratios of the total number of post-filtering and pre-filtering mutations have been evaluated as well, showing what percentage (shared or not) of the original mutations have remained in the variant file after the filtration process.

Using the first two measures, an additional performance score was introduced:

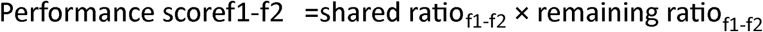

When ratios are measured in percentages, this score lies in the [0; 10,000] range, and the larger it is, the better the performance of the given pipeline. During the evaluations, sample-specific thresholds were used on the estimated probabilities made by the SOBDetector logistic model.

### SOBDetector Tool Usage

The documentation of the tool with examples of its usage, is available in the supplementary notes.

## RESULTS AND DISCUSSION

As a measure of the strand orientation, we defined the strand orientation bias (SOB) score, which is based on the relative orientation of the reads that are supporting the alternate alleles. SOBDetector was created to collect this additional information by reanalyzing the single-nucleotide variants stored in standard variant call format (VCF 4.0 and newer) files, and by using a Bayesian logistic regression classification model, evaluate whether a given mutation is a genuine one, or merely an artifact (Supplementary Figure 2). SOB scores lie in the [0,1] interval, and in theory, the closer they are to 1, the higher the probability, that the corresponding variants are sequencing artifacts.

In order to evaluate how reliable predictor this score is, we have used a set of TCGA (The Cancer Genome Atlas) donors (47 patients, 7 primary sites: Supplementary Figure 3), who had matched FF and FFPE whole exome sequences (WES), that were extracted from the same tumor. All the somatic variants present in the samples (called by GATK MuTect2 v3.7 (4)) were collected into a single dataset. When the two-dimensional distributions in the Tumor LOD (the higher this value, the more confident the mutation caller is about the variant) vs. SOB score space were compared, it became clear, that the FFPE samples were enriched in low-quality SNVs that had high SOB scores, and their distribution was highly dissimilar to the fresh frozen samples’ (Figure 1). After the elimination of the variants that had higher than 0.8 strand orientation bias scores, the two histograms became much more similar (Figure 1). Also, the great majority of the variants that had high SOB scores and more than 6 alternate allele supporting reads (in order to minimize the probability that the strand orientation was due to random chance), were mostly present in the FFPE samples only (Figure 2A, Supplementary Figures 4-5), indicating that the number of alternate allele-supporting reads, and the SOB scores might be an ideal combination for FFPE-artifact filtration. Due to the heterogenous nature of cancer samples(6), however, even true somatic mutations can be supported by only a few reads, therefore by setting a simple threshold on the number of alternate allele supporting reads and their SOB scores, it was clear that we would remove many genuine mutations too.

**Figure 1.:**
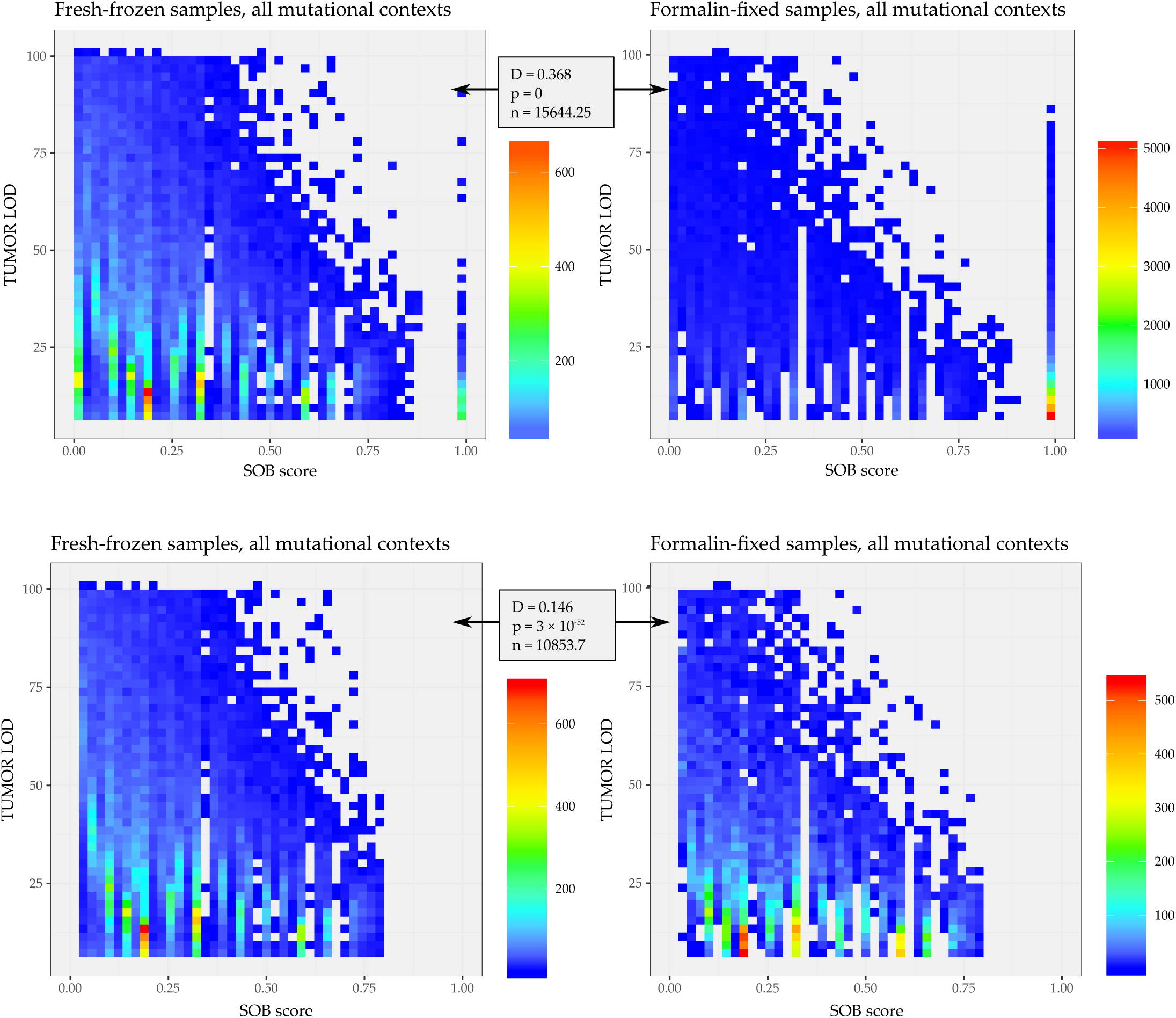
Read orientation bias detection. Collated two dimensional distributions of the variants in the TLOD-SOB score space, before filtering on the SOB scores. The color scale corresponds to the number of somatic mutations identified as genuine variants by the mutation caller. The majority of the mutations present in the FF samples (left side) lie in the [0,0.8] SOB score range, while mutations in the FFPE samples concentrated at low TLOD and high SOB score values. The similarity between the distributions was tested using a two-dimensional Kolmogorov-Smirnov test, the D-statistic and p-value of which being displayed between the corresponding panels. The bottom panels indicate that the similarity between the distributions increases after variants with larger than 0.8 SOB scores are removed.

**Figure 2.:**
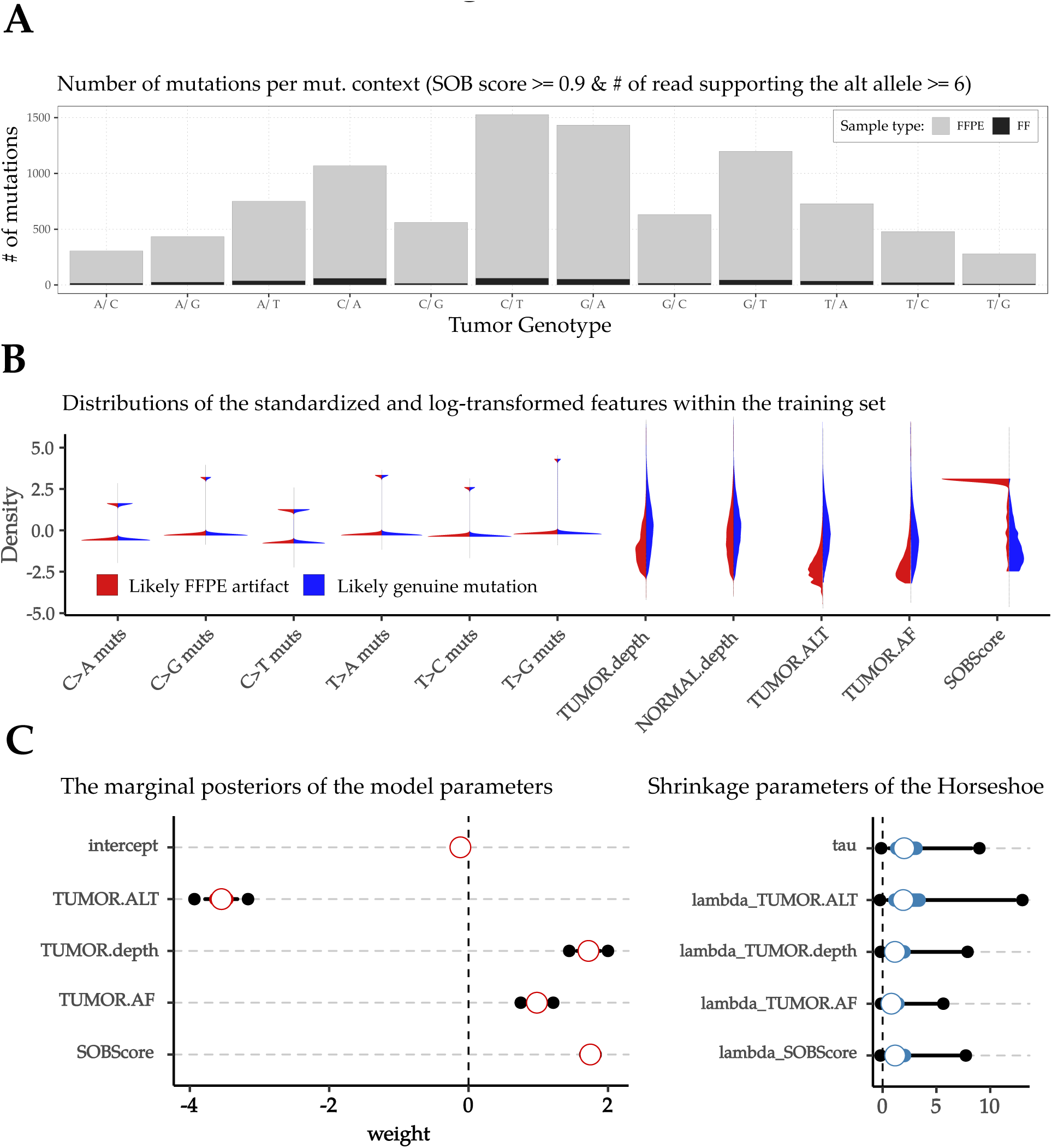
Defining the Bayesian logistic regression classification model. **A:** Summary of the number of mutations in each mutational context that have larger than 0.9 SOB scores and are supported by more than 6 read-pairs. Such variants are almost solely present in FFPE specimens, showing that the combination of these two features can be used as a crude way of artifact filtration. **B:** Distributions of the log-transformed and standardized features within the training dataset, separated into likely genuine mutation and likely artifact categories. The first six features are the 1 of K-encoded substitution subtypes. The remaining five features are the following. TUMOR.depth: number of reads spanning over the locus of a particular variant in the tumor, NORMAL.depth: number of reads spanning over the locus of the variant in the normal BAM, TUMOR.ALT: number of reads that are supporting the variant in the tumor BAM, TUMOR.AF: allele-frequency of the variant in the tumor, SOB score: the strand orientation bias score). Apart of the NORMAL.depth, all of these features distributed differently between the likely genuine mutations and likely FFPE artifacts, with the SOB scores showing the most striking disparity. **C:** On the left: The marginal posterior distributions of the weights of the final implementation of the logistic model. On the right: The marginal posterior distributions of the global (tau)- and local (lambda) shrinkage parameters of the horseshoe prior, that was used to ensure that the estimated linear-separator does not overfit the data.

Therefore, in order to maximize the accuracy of the artifact filtration step, we have collected the most likely genuine mutations (group 0) and the most likely FFPE artifacts (group 1) into a subset of the original collated dataset that could be used as a basis for a classification model. As possible training features we concentrated on attributes that are common to all VCF files or that can be reliably calculated regardless of the variant calling pipeline. Initially, the six possible substitution subtypes (C>A, C>G, C>T, T>A, T>C, T>G), the sequencing depths of the loci both in the tumor and the normal BAM files, the alternate allele frequencies, the number of alternate allele-supporting reads, and the variants’ SOB scores were considered. After investigating the linear separability of the two groups (Supplementary Figure 6), a Bayesian logistic regression model was selected for training, the weights of which were regularized by a horseshoe-prior (7). Surprisingly, the substitution subtypes did not turn out to be reliable predictors, indicating that the likely artifact group did not originate solely as the result of formalin fixation. Similarly, the depth of the normal BAM had no predictive power either. The most substantial difference between the two groups was between their SOB scores (Figure 2B, Supplementary Figure 8, Supplementary Table 2). The marginal posterior distributions of the final weights of the linear separator (Figure 2C, Supplementary Table 3) were determined using 32,444 variants (collected from the somatic VCFs of 44 patients with 7 different primary sites), approximately half of which belonged to group 0, the other half to group 1.

On an unbiased evaluation set of 5726 variants (n0 = 2832, n1=2894), which was set aside for evaluation purposes prior to training, the model performed reasonably well (Figure 3A). To estimate its predictive accuracy, an evaluation set-specific threshold was established at p(artifact|features) = 0.57 (Figure 3B), at which point the accuracy of the model was predicted to be in the [0.9, 0.91] range, with 95% credibility (Figure 3C).

**Figure 3:**
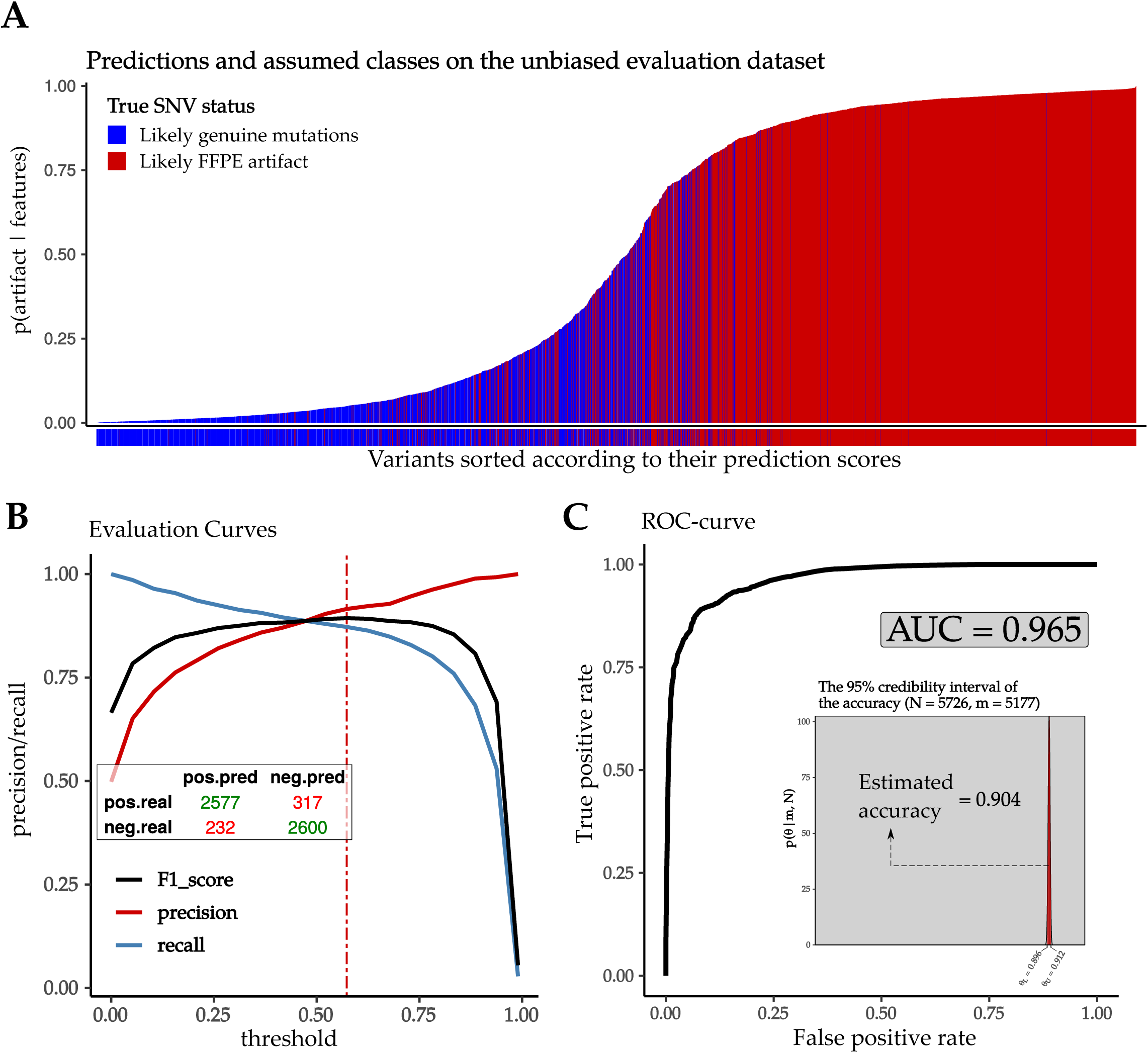
Evaluation of the predictions made by the logistic regression model. **A:** The predictions of the SOBDetector model on the 5726 variants of the unbiased evaluation set. The vertical axis shows the conditional probabilities that given a combination of features, the variant is an artifact. Likely artifacts with low prediction scores are most plausibly true mutations, that are only present in an FFPE sample due to intratumor heterogeneity, and were mistakenly categorized as artifacts, while likely genuine mutations with high prediction scores might be true mutations that have features resembling the FFPE artifacts’ (low number of variant-supporting reads that exhibit a strand orientation bias due to random chance, and hence high SOB scores). As a substitute for the mutation-labels, a small secondary figure is displayed below the bars. It follows the same color code than the main figure, and it is there to ensure, that the mutation/artifact states of the variants with lower predicted scores remain visible even for variants with lower probabilities. **B:** Precision: TP/(TP+FP) and recall: TP/(TP+FN) curves of the SOBDetector model, along with the F-scores: 2*precision*recall/(precision + recall), i.e. the harmonic mean of the precision and the recall. An evaluation set-specific threshold is proposed at the maximum value of the F1-curve (threshold = 0.57). **C:** ROC-curve and AUC value of the model, calculated using the predictions made on the unbiased evaluation set. The estimated accuracy: (TP + TN)/Total was calculated at the 0.57 threshold. The uncertainty was estimated using a binomial test.

A universal threshold could not be determined since the predictions strongly depend on the sequencing-properties (such as the mean sequencing depth, the tumor heterogeneity, and the quality) of the samples. Therefore, we have complemented SOBDetector with an optional sample-specific threshold estimation step. When the variants are sorted according to their predicted probabilities, on average their SOB scores increase. The threshold is set at the probability value, where the moving average of at least 5 (or maximum 1% of the total number of SNVs in the sample) consecutive SOB scores reaches above 0.8 (Supplementary Figure 9). This strategy was designed to maximize the concordance between the matched FFPE and FF samples of the patients. In order to compare the performance of the SOBDetector model to that of the currently available techniques, we have selected the invasive breast carcinoma (N=11), and colon adenocarcinoma cohorts (N=13) for concordance analysis. While the variant files of the breast cancer FFPE tissues had significantly more mutations than their FF counterparts, the difference between the colon cancer FFPE and FF VCFs was more subtle.

SOBDetector was compared to two GATK strand orientation bias filtering methods: FilterByOrientationBias from GATK 4.1.0, and LearnReadOrientationModel from GATK 4.1.2, using several pipeline configurations and three types of ratios. The first of these was the ratio of the shared mutations between matched FF and FFPE samples before and after filtering. This was significantly increased (from 8% to 39.7% on average) in all breast cancer pipelines, (Figure 4A), and in most of the colon cancer pipelines that involved SOBDetector (from 65.9% to 75.9% on average Supplementary Figure 10). This high performance from the part of SOBDetector, however, had an unfavorable side effect. When the ratios of the shared mutations prior and post filtering were measured, SOBDetector removed significantly more true-positives (i.e. likely true variants), than the GATK tools. On average, 14.3%, and 5.4% of the initially shared mutations were lost in the breast and colon cancer cohorts respectively (Supplementary Figures 11-12).

The ratio of the total number of variants after and before filtering was also measured. The numbers have clearly demonstrated, that by using the dynamic threshold, SOBDetector removes significantly more artifacts than true-positive variants, bringing the FF and FFPE mutational spectra much closer to each other than the other filtration techniques did. On average, SOBDetector has removed 74% of the original mutations from the FFPE variant files of the breast, and 18.9% of the colon cancer samples (Supplementary Figures 13-14).

As a benchmarking step a performance measure that accounts for both the ratio of removed likely artifacts and the loss of likely novel mutations was implemented. Ideally, both of these ratios should remain high, and when measured in percentages, their product can reach a maximum of 10,000 points. The pipelines involving SOBDetector had performed significantly better than the ones without it in the breast cancer cohort (Figure 4B), however in the colon cancer cohort such an exceptional performance was not observed (Supplementary Figure 15).

**Figure 4:**
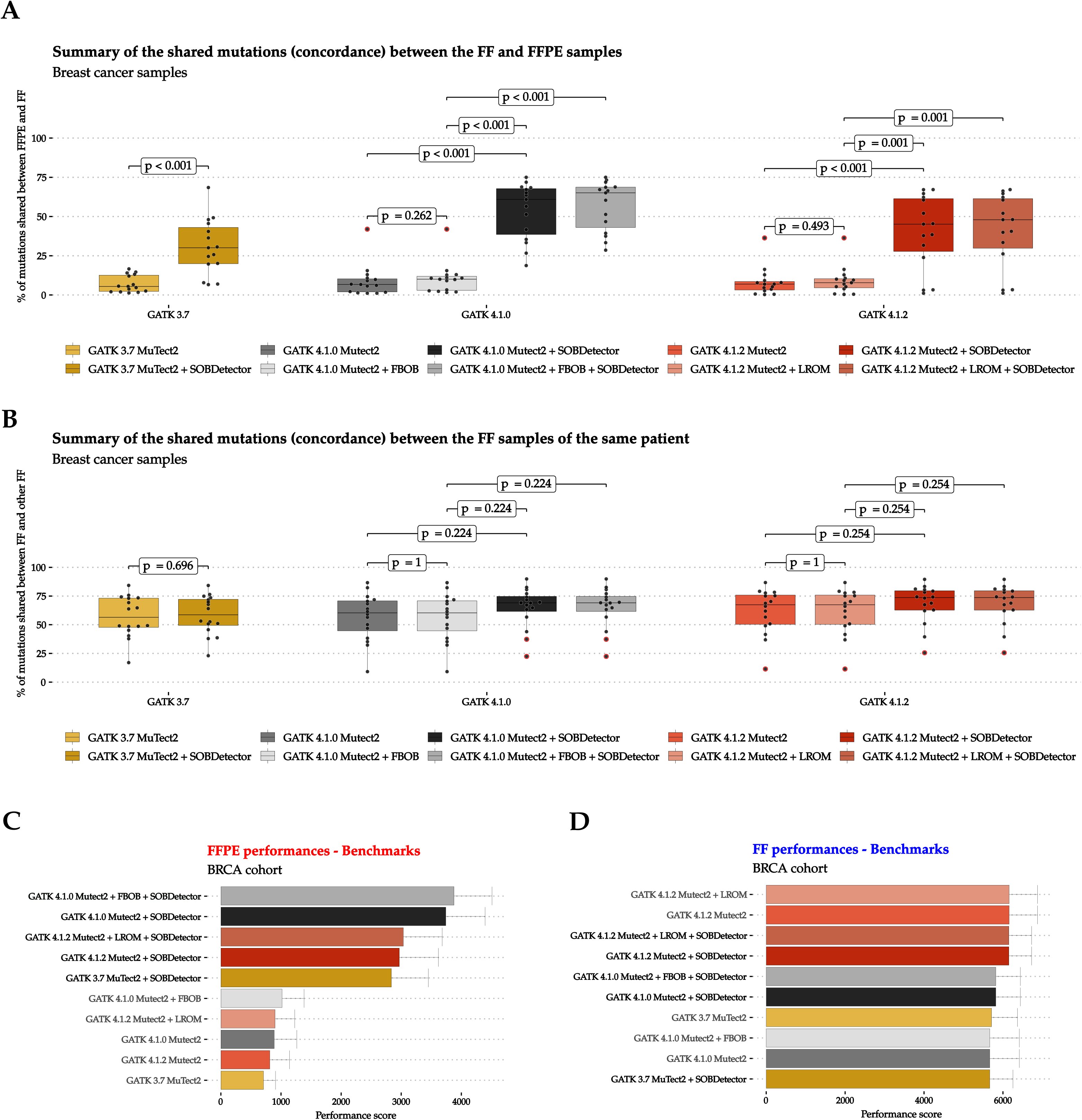
Benchmarking. **A:** Percentages of mutations shared by the matched FFPE and FF VCFs (n_ffpe = 15) derived from the same donors, using 10 different pipelines within the breast invasive carcinoma cohort (11 donors). When a sample had more than one FF-derived VCFs, the average of the shared ratio was calculated. Three major GATK implementations were considered. GATK 3.7, which marks the nightly-2016-02-25-gf39d340 built of the tool, GATK 4.1.1, GATK 4.1.2., GATK 4.1.1 + FilterByOrientationBias: FBOB, GATK 4.1.2 + LearnReadOrientationModel: LROM. All of these pipelines were checked alone, and with an additional SOBDetector filtering step. After an additional filtration using SOBDetector, the ratio of shared mutations between the FFPE and FF variant files has increased significantly in all pipelines. **B:** Percentages of mutations shared between a selected FF and the other FF samples of the same donor. Only those donors were considered who had at least two FF-derived variant files. The pipelines are the same as in panel d). Although an additional filtration using SOBDetector increased the ratio of the shared mutations somewhat, the differences were not significant between any of the comparable pipelines. **C:** The mean performance scores of the pipelines evaluated on the FFPE-derived variant files of the breast invasive carcinoma cohort. The error bars represent the standard errors of the means. The scores were calculated by multiplying the average percentage of shared mutations (between the FFPE and corresponding FF VCFs) by the percentage of the remaining shared mutations after the artifact-filtration step (in order to account for the removal of genuine mutations). The achievable scores vary between 0 and 10,000: the higher the score, the better the performance. Using this measure, the pipelines that were complemented with a SOBDetector filtration step, had performed significantly better than the ones without it. **D:** The mean performance scores of the pipelines on the FF-derived variant files. The error bars represent the standard errors of the means. The scores were calculated as it is described in panel e), however, instead of FFPE samples, FF originating from the same donors were compared. None of the pipelines performed significantly better (or worse) than the others.

We have also investigated whether an additional filtration with SOBDetector can increase the concordance between matched FF samples (same patient, same tumor). The same ratios and performance scores have been calculated as above, and although SOBDetector seemed to increase the ratio of shared mutations a little, this increase could not compensate for the loss of likely genuine mutations, hence the performance scores of the pipeline with and without SOBDetector remained in the same range in both cohorts (Figure 4C,D).

The predictive capabilities of the SOBDetector model were also tested on two non-TCGA cancer cohorts from previous studies (2,8), which were unrelated to the training process. Although the definition of likely artifacts was considerably weaker than in the TCGA case, and the effects of intratumor heterogeneity could not be eliminated, SOBDetector performed adequately well on both cohorts, and it outperformed the GATK strand-orientation bias filtering approach. The complete analysis of these cancer cases is available in the supplementary notes, in the form of separate case studies (Supplementary Figures 16-54).

Due to the possible loss of likely genuine mutations, we recommend to use the dynamic threshold and hard filtering functionality of SOBDetector, when instead of the individual mutations, one is interested in the large scale patterns of true somatic variants, such as the triple-nucleotide substitution signatures (9). In such cases, the loss of a few true somatic mutations is less important than the removal of as many artifacts as possible. However, when the individual genotypes are more important, such as during the identification of therapeutically targetable mutations, no threshold should be used. Instead, the predicted probabilities of the logistic model accompanied by the SOB scores of the variants should only be used as an additional information about the state of the mutation.

## AVAILABILITY

SOBDetector is implemented in Java 1.8, and for academic use, it is freely available at: www.github.com/mikdio/SOBDetector

## ACKNOWLEDGMENTS

The results shown here are based upon data generated by the TCGA Research Network: http://cancergenome.nih.gov/

