## Supplementary Notes for "Strand Orientation Bias Detector (SOBDetector) to remove FFPE sequencing artifacts"

### Strand Orientation Bias Detector (SOBDetector) for FFPE artifact filtration

*Written by:*

*Date of the latest update:*
NOVEMBER 25, 2019

### List of Tables

2 Summary of the marginal posterior probabilities of the first model, estimated by the markov-chain monte carlo walkers. 11
3 Summary of the marginal posterior probabilities of the final model, estimated by the markov-chain monte carlo walkers. 11

### 1 Supplementary tables not included in the Supplementary Text

Supplementary Table 1 is available separately, in xls format.

- **Supplementary Table 1**: The TCGA test cohort and Analysis IDs

#### 1.1 Introduction

There are two options to choose from when collecting and preserving tumor specimens for molecular analysis: fresh-frozen (FF) or formalin-fixed paraffin-embedded (FFPE). While inserting a tissue into a phenol solution and fresh freezing it directly after resection results in an excellent stability, keeping it under -80 °C is indisputably more expensive, than embedding it into paraffin blocks after formalin fixation. This fixation process however, introduces artifactual mutations (mostly in the C>T direction due to the deamination of cytosine bases induced by formalin) into the DNA strands which results in a much lower quality. The major benefit of this ffpe procedure is that the specimen becomes no longer sensitive to heat, so it can be work with at room temperature.

Due to its cost effectiveness and simplicity, ffpe tissue processing remains the most common approach for tissue specimen storage. However, the presence of these artifactual mutations makes mutation calling (especially point mutation calling) a hard task. Over the past years, various amounts of tools have been created that have made the extraction of allele-specific segmentation data and copy number profiles from whole exome sequences (WES) possible (sequenza, ASCAT, . . . ). However, in order to ensure this allele specificity these tools rely on heterogeneous positions, which can be strongly affected by the presence of these ffpe artifacts, making the final copy number estimations unreliable. Also, the well known somatic signatures (Alexandrov et al.) depend solely on the point mutations found in a tumor sample. It is imperative therefore, that before performing these analyses we ensure, that the mutations found by a mutation caller are real, not mere artifacts.

Since formalin very likely affects only one of the strands (e.g. a C|G pair becomes T|G), a paired-end next generation sequencing approach can help in this additional filtering step. By counting not just the number of reads that support the alternative alleles, but the relative orientation of the reads as well (Forward-Reverse:FR or Reverse-Forward:RF), these ffpe artifacts will likely have a strand orientation bias towards one of the directions, while true mutations should have approximately the same amount of FR and RF reads. This article introduces the Strand Orientation Bias Detector (SOBDetector) tool, that reanalyzes the mutations stored in a vcf, and evaluates whether the reads that support the alternate alleles have a strand orientation bias using the original binary alignment (BAM) files. This method works only on Illumina-like sequencing approaches.

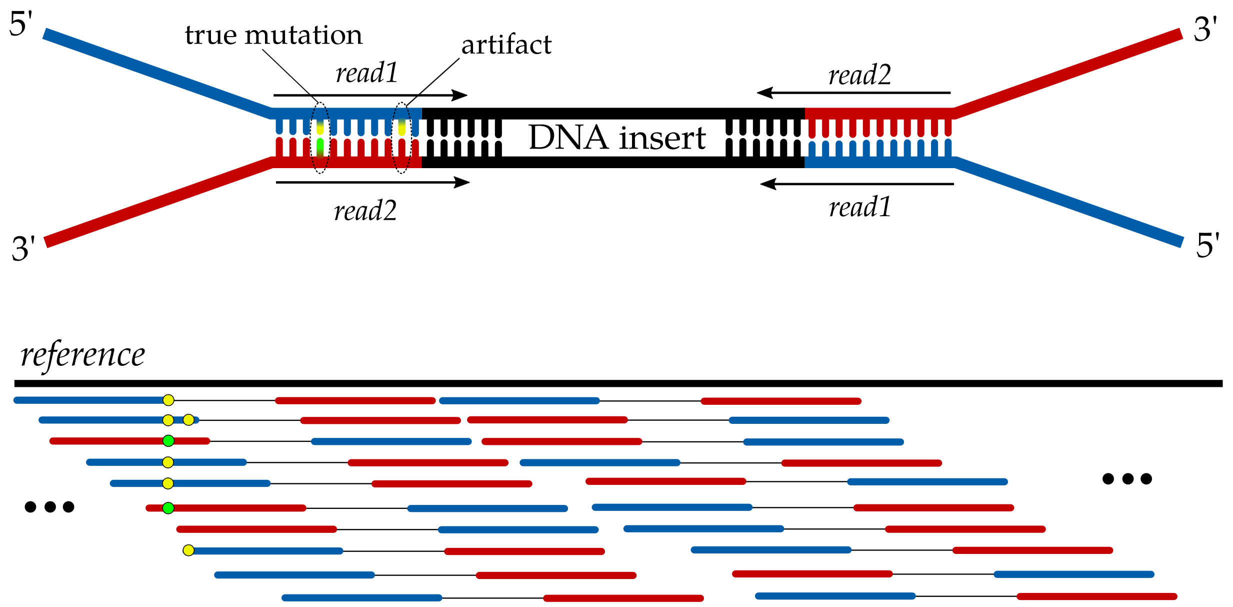

**Suppl.Fig. 1:** Illustrating the manifestation of sequencing artifacts caused by formalin fixation compared to true mutations. Since FFPE artifacts most likely affect only one of the strands, during mutation calling, these artifacts will manifest as mutations with low alternate allele frequencies, and with a strand orientation bias.

#### 1.2 Approach

Although formalin fixation induces multiple types of DNA damage, the most prominent of them is definitely the deamination of cytosines, wich introduces extra C>T/G>A mutations [**?**]. Since these alterations occur before the ligation of Illumina adapters, they are likely to have a read orientation bias. SOBDetector is a simple tool to filter most of them out.

During sequencing, we first fragment the DNA in (ideally) isotermal conditions. Than we ligate the adapter sequences, the primers, indices and sequences that are complementary to the oligose found in the flow cell to the ends of these fragments. Since both the Crick and Watson strands have read1s and read2s on them, after the alignment there should be approximately the same amount of F1R2 and F2R1 read-pairs in the resulting binary alignment files. Since the mutations induced by the storing process can be found only in one of the strands, they are likely to have a bias in the F1R2 and F2R1 orientation as well (Suppl.Fig.1).

As a measure of this orientation bias, we introduce the Strand Orientation Bias (SOB) score:

$$\mathrm{SOB} = \left| \frac{\#\mathrm{ofF1R2}_{alt} - \#\mathrm{ofF2R1}_{alt}}{\#\mathrm{ofF1R2}_{alt} + \#\mathrm{ofF2R1}_{alt}} \right|, \qquad \mathrm{SOB} \in [0,1],$$

where $\#FiRj_{alt}$ $(i,j \in \{1,2\})$ is the number of paired reads, that are in an $FiRj$ orientation, and are supporting the alternate allele. The distribution of the possible values of this score naturally depends on the total number of reads that support the alternative allele, hence it is affected by the sequencing depth of the sample, and the tumor's heterogeneity.

#### 1.3 SOBDetector

Since not all the variant calling tools collect the necessary strand orientation information of the reads that cover the loci of identified SNVs, we have developed SOBDetector, a simple tool, that goes back to the binary alignment (BAM) files, from which the variants were called, and appends the strand orientation of their reads along with the SOB scores to each of them. A simple sketch of the tool is available in Suppl.Fig. 2.

In order to maximize its portability, SOBDetector was implemented in java (v1.8). In theory it works on any operating system on which samtools can operate functionally, however it was only tested on linux and MacOS kernels. The tool is built around two big classes: `SOBDetector`, and `Vcf`. Its pipeline can be summarized through the following key points:

- The app instantiates a SOBDetector and a Vcf object. The processing methods are included in the SOBDetector class, while informations about the variants (chrom, ref, alt, position, filter, info, etc. . . ) are stored in the Vcf object.
- At first, the app checks the input parameters and if it finds that everything is fine, it calls the start() method.
- If an input vcf file is available at the specified location, then the readVcf() method stores its contents in the Vcf object.

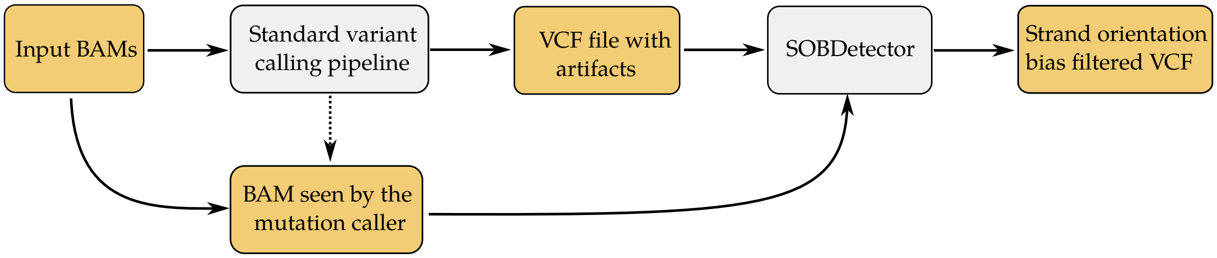

**Suppl.Fig. 2:** Standard SOBDetector usage pipeline. The tool uses a BAM file (germline or somatic) and its corresponding variant calling output: a VCF or a tabulated file. To ensure that SOBDetector analyzes the same reads than the varaint caller, it is important that if the variant caller realigns some of the reads, SOBDetector should get the realigned bam as well.

- The bamProcessor() method then goes through the variants (one by one) of the Vcf object, and with the help of samtools it collects the reads, that cover the location of the variants. The reads are collected into an ArrayList of Strings. If the user is interested only in those variants that have passed the previously determined filters, they can set the onlyPassed input variable to true, in which case variants with non-PASSED FILTER fields will be ignored. This step may decrease the necessary running time substantially.
- If either the REF or ALT attributes of the `vcf` contain more than one characters (i.e. we are dealing with an indel), the location is skipped. (The tool does not consider indels yet.)
- If the user specifies a minimally considerable mapping quality, then it is checked as well.
- For every single read, the tool has to process the cigarStrings as well, since deletions and insertions are encoded there. If a read contains events which increase its length, then the appropriate number of D (deletion), I (insertion), H (hard-clipping) characters are appended to the appropriate locations.
- The user can also provide a minimum for the base quality. In order to check that, the quality strings are converted to Phred33+ quality scores, then if at the analyzed variant position the quality turns out to be smaller then the threshold defined by the user, then the read gets ignored (to ensure that the qualities and sequences are in sync, the quality strings get extended by the same amount of D, H, and I characters than the actual sequential context).
- If a read passes through the (optional) quality thresholds, the tool instantiates a SamFlag object, the methods of which are designed to convert the decimal Sam-flag codes to their actual meaning. Reads that are flagged as "duplicate" or "unmapped" are filtered out here.
- If the read can still be considered, then the analyzed position is checked. The read can support the reference-, the alternate allele, or one of the other two possibilities. The number of ref-supporting, alt-supporting and other-supporting reads are incremented here.
- For all three categories, we also check the strand orientation, which at this point is stored in the SamFlag object, and increment the appropriate variables by one.
- All the strand orientation info corresponding to the analyzed variants are stored in the Vcf object.
- After the tool has finished with the last variant position in the vcf, the Bayesian logistic model method gets called. The collected attributes are log-transformed and standardized, by default using the sample's own means and standard deviations, but the user can also supply alternative values (for example if the sample is part of a greater cohort, and the cohort's statistics would make more sense to use on each sample).
- The standardized features then are feeded to the prediction model, which utilizes the posterior distributions of the horseshoe logistic weights to calculate the predicton scores, which are collected into a separate feature, called pArtifact.
- After this prediction step, the writeVcf() method is called. It writes the contents (now extended with the strand orientation info and the predictions) back to a vcf file, the location of which should be specified by the user (-outputVcf).

###### 1.3.1 Running SOBDetector

The current version (v1.1) of SOBDetector can be used in the following two ways.

###### By using a vcf file

The first is the regular one, when the user provides the location of a vcf file and its corresponding bam's. This can be implemented in the following way:

```
1 java -jar SOBDetector_v1.0.jar \
2     --input-type VCF \
3     --input-variants ./input-variants.vcf \
4     --input-bam ./input.bam \
5     --output-variants ./output.vcf \
6     --only-passed true
```

In line 1 we provide SOBDetector for the java virtual machine. Optional java arguments such as the maximum amount of allowed memory or number of threads can be specified here, though multi-threading is not supported yet and the only thing that is stored in the heap is the vcf file itself, which is generally less than 100 Mb in size. From line two, we provide the mandatory arguments. These are the following:

- −−**input-type** The type of the provided variant file. It can either be `VCF` or `Table` (it will be discussed the latter later).
- −−**input-variants** Absolute or relative path that leads to the variant file (in this case, to a vcf).
- −−**inpupt-bam** Absolute or relative path of the binary alignment file, from which the variants were determined. It is important that in case the variant caller alters the original orientation of the reads while its running, for example by performing local de-novo assembly, we use the altered BAM file here.
- −−**output-variants** Absolute or relative path to the desired output file. If the input is a vcf file, then this will be a vcf too. Note: all the folders must exist, the tool won't create them automatically.

A semi-mandatory argument:

- −−**sample-name** It is possbile that a BAM file contains reads from mutliple samples. If this is the case, the header of the bam must contain the name of the samples. When SOBDetector encounters multiple sample names among the read groups, it produces an error. For example:

```
ERROR: /home/user/Path/samples.bam contains reads from multiple samples:
SAMPLE1
SAMPLE2
SAMPLE3
Use the --sample-name argument to specify which one you are interested in!
```

  It asks the user to specify the sample of interest with the −−**sample-name** argument, for example:

```
java -jar SOBDetector_v1.0.jar \
    --input-type VCF \
    --input-variants ./input-variants.vcf \
    --input-bam ./input.bam \
    --sample-name SAMPLE1 \
    --output-variants ./output.vcf \
    --only-passed true
```

The rest of the arguments are optional:

- −−**only-passed** Defaults to false, if set true, variants that have non-PASSED FILTER attributes in the input vcf will be ignored. (It speeds up the execution time.)
- −−**minBaseQuality** Defaults to 0, the minimally considered base quality. If the quality of the considered base is less than this value, the read will be ignored.
- −−**minMappingQuality** Defaults to 0, the minimally considered mapping quality. If the mapping quality of read is less than this value, then the read is ignored.
- −−**standardization-parameters** The Bayesian prediction model requires the standardization and log-transformation of the predictive variables ($\mathbf{x}_i$):

$$\mathbf{x}'_i = \frac{\mathbf{x}_i - \mu_i}{\sigma_i}, \qquad \rightarrow \qquad \mathbf{z}_i = \log(1 + \mathbf{x}'_i)$$

where $\mu : i$ and $\sigma_i$ are the mean and standard deviation of attribute $i$. The attributes are: **TUMOR.ALT**: number of alternate allele supporting reads at a given genomic coordinate, **TUMOR.depth**: number of reads covering a given genomic coordinate, **TUMOR.AF**: tumor allelic frequency (AF = ALT/depth), **SOBscore**: the strand orientation bias score, evaluated by the tool. By default, the tool uses the means and standard deviations of the TCGA dataset, on which the Bayesian model was originally trained, however, when the user have multiple samples, that belong to the same cohort, it is worth to use the means and standard deviations of the whole cohort instead. This can be specified by the −−**standardization-parameters** argument. The parameters need to be stored in a simple tab-delimited text file, the first line of which should contain the means, the second the standard deviations of the four attributes. Example `altStandardization.txt`:

```
TUMOR.ALT	TUMOR.depth	TUMOR.AF	SOBScore
3.367	14.71	0.544	0.513
0.811	4.052	0.152	0.227
```

##### The output

The output in this case is also a vcf file: the original vcf with the appended strand orientation bias info. The following line shows an example of this extra info through a single variant:

```
numF1R2Alt=3;numF2R1Alt=1;numF1R2Ref=13;numF2R1Ref=15;numF1R2Other=0;numF2R1Other=0;SOB=0.5000;
    pArtifact=7.504e-01;artiStatus=snv
```

- **numF1R2Alt**: Number of alternate allele supporting reads in the F1R2 orientation
- **numF2R1Alt**: Number of alternate allele supporting reads in the F2R1 orientation
- **numF1R2Ref**: Number of reference allele supporing reads in the F1R2 orientation
- **numF2R1Ref**: Number of reference allele supporting reads in the F2R1 orientation
- **numF1R2Other**: Number of F1R2 reads that do not support the reference nor the alternate alleles (example, REF:A, ALT:C, and the read supports a T)
- **numF2R1Other**: Number of F2R1 reads that do not support the reference nor the alternate alleles
- **SOB**: The strand orientation bias score calculated from the first two attributes of this list.
- **pArtifact**: The posterior probability that the variant is a sequencing/storing/preparation/etc... artifact
- **artiStatus**: The predicted status of the variant, it is either **snv** or **artifact**

##### 1.3.2 Using a tabulated file as input

Many times the variants we are working with are collected into tab-delimited files (or R/pandas data frames). SOBDetector can work with this format too. All we have to do is to specify our intentions by setting the input-type to `Table` instead of `Vcf`:

```
java -jar SOBDetector_v1.0.jar \
    --input-type Table \
    --input-variants ./input.table \
    --input-bam ./input.bam \
    --output-variants ./output.table \
```

CHROM POS REF ALT

Only the first four columns of such tab delimited files are mandatory. The necessary order of these attributes (using the standard vcf notation) has to be the following:

The file might contain other columns as well, those will not be assessed. The output file of the analysis will have the same format as the input, containing the same columns, but additional attributes will be appended to them with the strand bias information.

An example of the contents of such a tabulated file can be seen below:

| CHROM | POS | REF | ALT | NLOD | TLOD |
|---|---|---|---|---|---|
| chr5 | 236441 | T | G | 21.57 | 5.56 |
| chr5 | 256357 | G | A | 1.58 | 24.59 |
| chr5 | 256458 | A | G | 8.73 | 6.51 |
| chr5 | 413313 | ATT | A | 2.86 | 4.87 |
| chr5 | 745329 | T | A | 10.53 | 4.83 |

Naturally, the chromosome notation has to be the same as in the BAM file. The output after running SOBDetector on it:

| CHROM | POS | REF | ALT | NLOD | TLOD | numF1R2Alt | numF2R1Alt | numF1R2Ref | numF2R1Ref | numF1R2Other | numF2R1Other | SOB |
|---|---|---|---|---|---|---|---|---|---|---|---|---|
| chr5 | 236441 | T | G | 21.57 | 5.56 | 6 | 2 | 32 | 29 | 0 | 0 | 0.500 |
| chr5 | 256357 | G | A | 1.58 | 24.59 | 15 | 15 | 61 | 59 | 1 | 2 | 0.000 |
| chr5 | 256458 | A | G | 8.73 | 6.51 | 5 | 3 | 23 | 20 | 0 | 0 | 0.250 |
| chr5 | 413313 | ATT | A | 2.86 | 4.87 | . | . | . | . | . | . | . |
| chr5 | 745329 | T | A | 10.53 | 4.83 | 4 | 2 | 11 | 25 | 0 | 0 | 0.333 |

The same optional arguments can be used in the `Table` mode as in the `Vcf` mode, however the −−**only-passed** argument has no effect on the running time, since it is not checked.

#### 1.4 Additional Supplementary figures

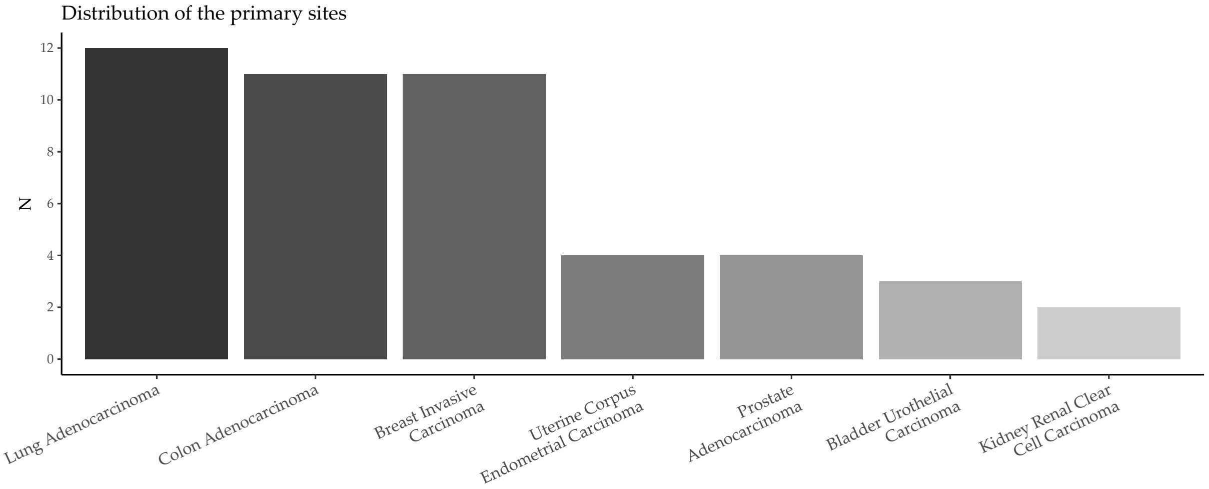

**Suppl.Fig. 3:** A summary about the number of patients and the primary tumor sites that were considered in the analysis.

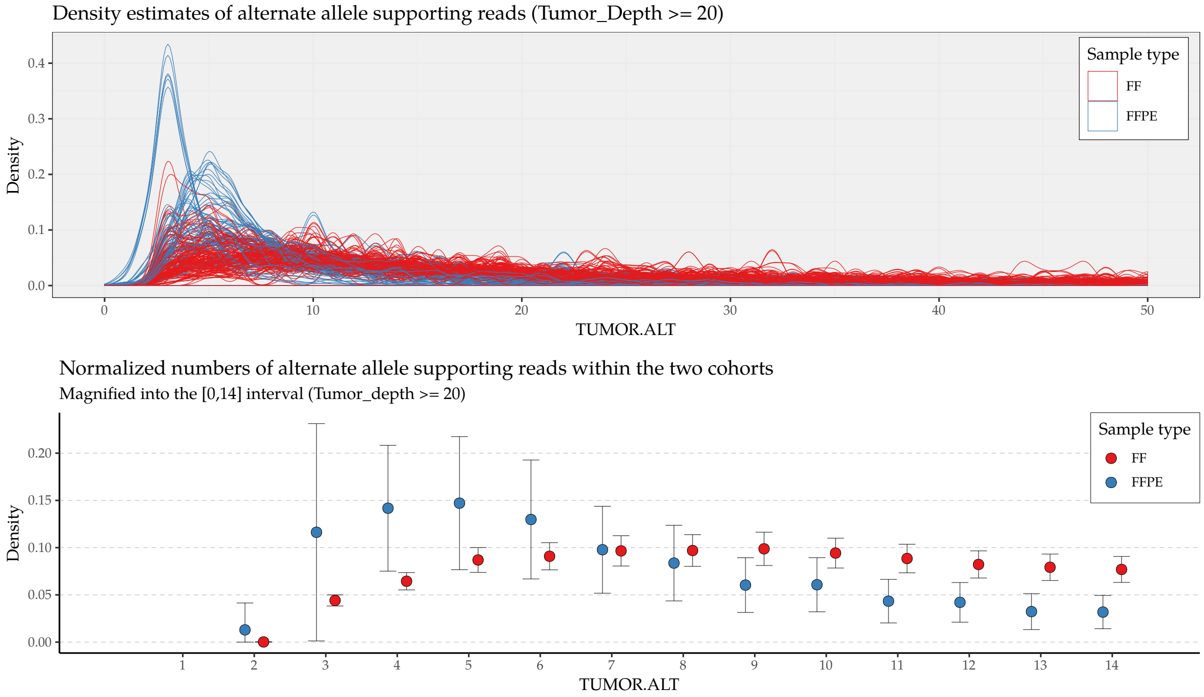

**Suppl.Fig. 4:** Top panel: The distributions of the number of alternative allele supporting reads in each of the somatic TCGA (MuTect2) vcfs, estimated by a kernel smoother strategy (epanechnikov kernel with width=1). The coloring represents the fixation process (FF or FFPE).

Bottom panel: The mean normalized distributions of the two sample types, magnified into the [0,14] TUMOR.ALT range. The error bars represent the standard deviations.

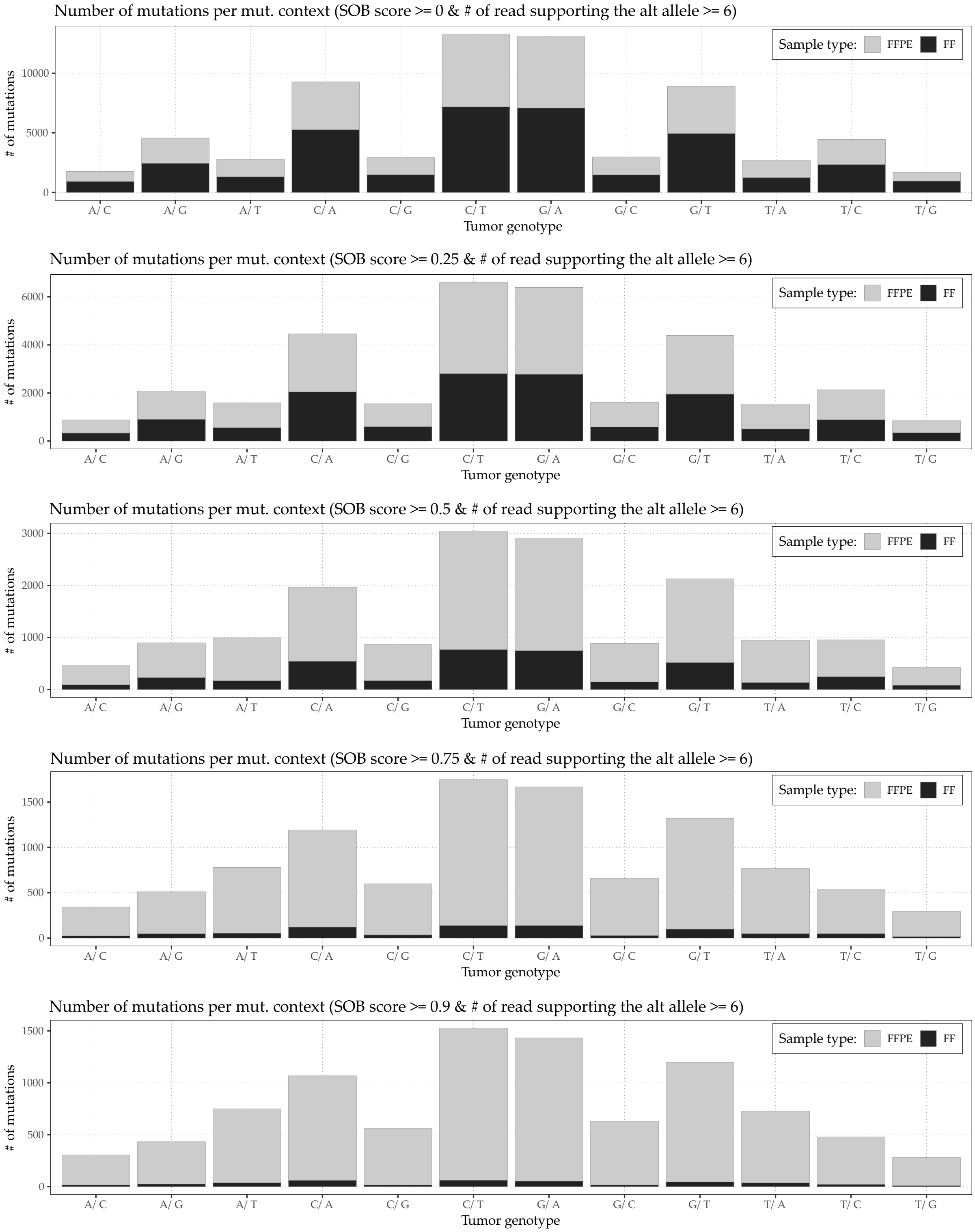

**Suppl.Fig. 5:** The figure was created using the dataset that contained all the variants from the downloaded vcfs, that were supported by at least 6 reads. When we further filter these mutations by setting an additional threshold on their SOBScores, it becomes clear, that above SOBScore > 0.8, the remaining mutations are almost exclusively present only in the FFPE samples.

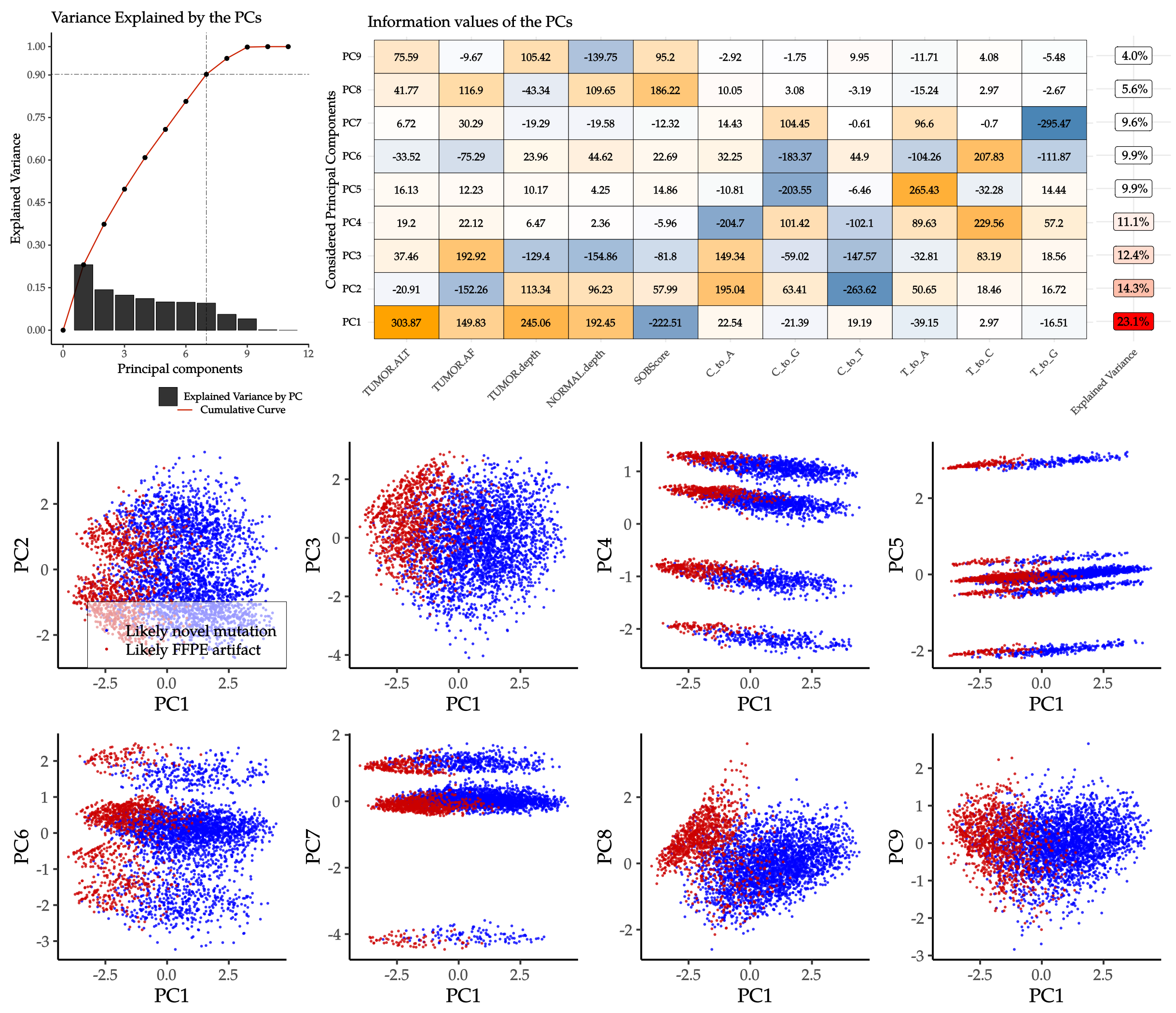

**Suppl.Fig. 6:** On top: Information content (i.e. the eigenvectors multiplied by their corresponding eigenvalues), and explained variance of the principal components.
On bottom:Projection of a subset of the trainable dataset, containing 5000 randomly selected samples to the planes spanned by the first 9 principle components of the whole set. The horizontal axis always contains the first PC, while the verticle axes iterate over PC2 to PC9.

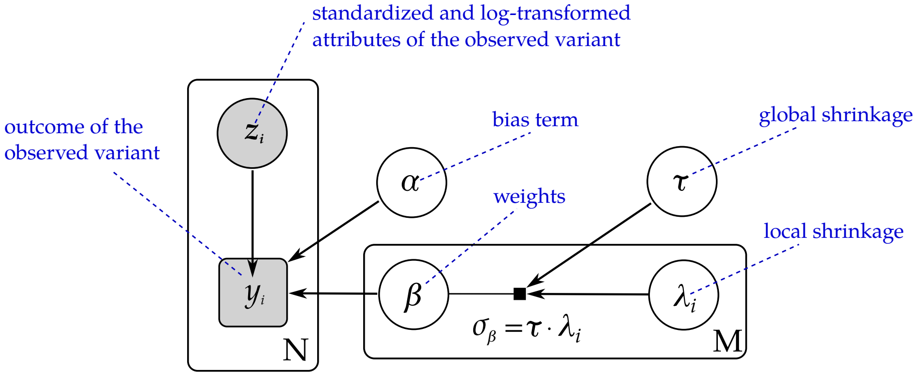

**Suppl.Fig. 7:** Graphical representation of the horseshoe-regularized Bayesian logistic regression model. Grey shaded nodes are the observed variables, white nodes are all unobserved model parameters. The two rectangles around the nodes with *N* and *M* indicate iterations.

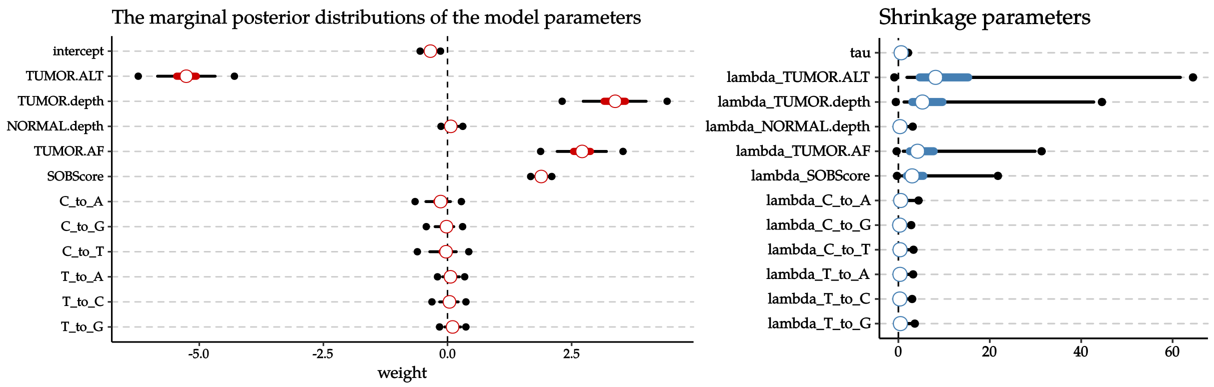

**Suppl.Fig. 8:** Representation of the marginal posteriors of the model weights (including the intercept) when the horseshoe prior was used. The middle red circles correspond to the means of the distributions, the red and black segments on both sides of the mean represent the 50% and 95% inter-quartile ranges respectively. The black points are the extrema (min and max). The right panel of the figure illustrates the marginal posteriors of the global (*tau*) and local shrinkage (*lambdas*) parameters.

| | mean | se_mean | sd | 2.5% | 25% | 50% | 75% | 97.5% | n_eff | Rhat |
|---|---|---|---|---|---|---|---|---|---|---|
| **intercept** | -0.34 | 0.00 | 0.06 | -0.47 | -0.39 | -0.34 | -0.30 | -0.22 | 2822.25 | 1.00 |
| **TUMOR.ALT** | -5.26 | 0.00 | 0.30 | -5.84 | -5.46 | -5.26 | -5.06 | -4.68 | 3735.67 | 1.00 |
| **TUMOR.depth** | 3.37 | 0.01 | 0.32 | 2.73 | 3.15 | 3.38 | 3.59 | 4.00 | 3653.82 | 1.00 |
| **NORMAL.depth** | 0.07 | 0.00 | 0.08 | -0.06 | 0.01 | 0.07 | 0.12 | 0.24 | 2064.65 | 1.00 |
| **TUMOR.AF** | 2.71 | 0.00 | 0.25 | 2.21 | 2.54 | 2.71 | 2.88 | 3.20 | 3152.57 | 1.00 |
| **SOBScore** | 1.89 | 0.00 | 0.07 | 1.76 | 1.84 | 1.89 | 1.93 | 2.02 | 6628.14 | 1.00 |
| **C_to_A** | -0.15 | 0.00 | 0.13 | -0.44 | -0.22 | -0.14 | -0.06 | 0.06 | 939.36 | 1.01 |
| **C_to_G** | -0.03 | 0.00 | 0.09 | -0.25 | -0.07 | -0.02 | 0.02 | 0.13 | 554.34 | 1.01 |
| **C_to_T** | -0.05 | 0.00 | 0.13 | -0.36 | -0.11 | -0.03 | 0.02 | 0.18 | 843.67 | 1.01 |
| **T_to_A** | 0.06 | 0.00 | 0.09 | -0.10 | 0.01 | 0.06 | 0.12 | 0.24 | 1395.37 | 1.01 |
| **T_to_C** | 0.04 | 0.00 | 0.09 | -0.16 | 0.00 | 0.04 | 0.10 | 0.22 | 768.16 | 1.01 |
| **T_to_G** | 0.10 | 0.00 | 0.08 | -0.06 | 0.05 | 0.10 | 0.16 | 0.27 | 1024.19 | 1.01 |
| **lambda_TUMOR.ALT** | 13.88 | 0.41 | 23.25 | 1.89 | 4.66 | 8.14 | 15.28 | 61.69 | 3203.05 | 1.00 |
| **lambda_TUMOR.depth** | 9.03 | 0.35 | 14.00 | 1.24 | 3.04 | 5.27 | 9.65 | 42.74 | 1579.83 | 1.00 |
| **lambda_NORMAL.depth** | 0.67 | 0.02 | 1.64 | 0.01 | 0.15 | 0.35 | 0.73 | 3.00 | 7086.48 | 1.00 |
| **lambda_TUMOR.AF** | 7.41 | 0.25 | 18.15 | 1.06 | 2.51 | 4.19 | 7.73 | 29.90 | 5253.32 | 1.00 |
| **lambda_SOBScore** | 5.06 | 0.13 | 9.33 | 0.77 | 1.84 | 3.01 | 5.46 | 20.71 | 4903.42 | 1.00 |
| **lambda_C_to_A** | 0.99 | 0.08 | 3.91 | 0.04 | 0.24 | 0.50 | 1.00 | 4.20 | 2643.16 | 1.00 |
| **lambda_C_to_G** | 0.58 | 0.02 | 1.20 | 0.01 | 0.11 | 0.28 | 0.65 | 2.71 | 6138.21 | 1.00 |
| **lambda_C_to_T** | 0.67 | 0.02 | 1.25 | 0.02 | 0.15 | 0.36 | 0.74 | 3.17 | 5515.44 | 1.00 |
| **lambda_T_to_A** | 0.67 | 0.02 | 1.31 | 0.02 | 0.15 | 0.35 | 0.75 | 3.09 | 4482.81 | 1.00 |
| **lambda_T_to_C** | 0.61 | 0.02 | 1.21 | 0.02 | 0.13 | 0.31 | 0.70 | 2.90 | 6446.97 | 1.00 |
| **lambda_T_to_G** | 0.77 | 0.02 | 1.42 | 0.03 | 0.20 | 0.44 | 0.87 | 3.42 | 5494.26 | 1.00 |
| **tau** | 0.71 | 0.01 | 0.46 | 0.17 | 0.40 | 0.60 | 0.91 | 1.89 | 2895.65 | 1.00 |
| **lp__** | -1150.59 | 0.06 | 3.59 | -1158.68 | -1152.79 | -1150.23 | -1148.07 | -1144.56 | 3712.01 | 1.00 |

**Suppl.Tab. 2:** Summary of the marginal posterior probabilities of the first model, estimated by the markov-chain monte carlo walkers.

| | mean | se_mean | sd | 2.5% | 25% | 50% | 75% | 97.5% | n_eff | Rhat |
|---|---|---|---|---|---|---|---|---|---|---|
| **intercept** | -0.25 | 0 | 0.02 | -0.29 | -0.26 | -0.25 | -0.23 | -0.2 | 9311.33 | 1 |
| **TUMOR.ALT** | -3.34 | 0 | 0.13 | -3.59 | -3.42 | -3.34 | -3.25 | -3.09 | 3912.39 | 1 |
| **TUMOR.depth** | 1.17 | 0 | 0.09 | 0.99 | 1.11 | 1.17 | 1.23 | 1.35 | 3821.62 | 1 |
| **TUMOR.AF** | 0.77 | 0 | 0.08 | 0.61 | 0.71 | 0.77 | 0.82 | 0.93 | 3805.95 | 1 |
| **SOBScore** | 1.8 | 0 | 0.03 | 1.75 | 1.78 | 1.8 | 1.81 | 1.85 | 10324.46 | 1 |
| **lambda_TUMOR.ALT** | 3.24 | 0.14 | 11.09 | 0.45 | 1.19 | 1.96 | 3.46 | 12.7 | 6242.91 | 1 |
| **lambdaq_TUMOR.depth** | 1.5 | 0.04 | 2.6 | 0.2 | 0.56 | 0.95 | 1.63 | 5.93 | 4098.67 | 1 |
| **lambda_TUMOR.AF** | 1.27 | 0.09 | 4.57 | 0.14 | 0.42 | 0.75 | 1.32 | 4.84 | 2825.25 | 1 |
| **lambda_SOBScore** | 1.95 | 0.04 | 2.96 | 0.28 | 0.76 | 1.27 | 2.17 | 7.6 | 4753.93 | 1 |
| **tau** | 2.29 | 0.03 | 1.82 | 0.53 | 1.16 | 1.81 | 2.79 | 7.07 | 4479.64 | 1 |
| **lp__** | -8043.61 | 0.04 | 2.37 | -8049.09 | -8044.98 | -8043.27 | -8041.86 | -8039.97 | 3660.91 | 1 |

**Suppl.Tab. 3:** Summary of the marginal posterior probabilities of the final model, estimated by the markov-chain monte carlo walkers.

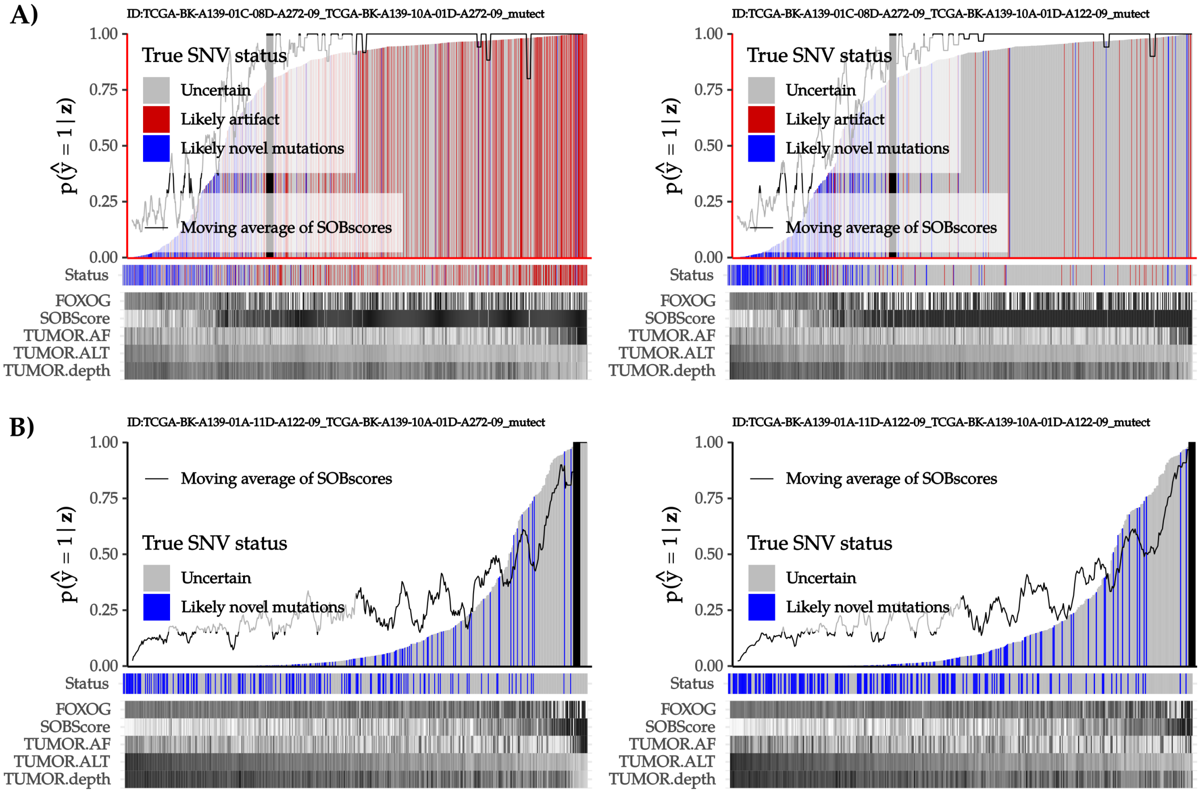

**Suppl.Fig. 9:** Examples for the dynamic detection of the prediction threshold. **A:** Two typical breast cancer FFPE samples with assumed artifacts, genuine mutations (red and blue bars respectively), and uncertain variants (grey bars). Genuine mutations and likely artifacts clearly separate when the variants are sorted according to their prediction scores. The moving average of the SOBscores shows an increasing trend, and a threshold can be defined where it becomes stationary at $\langle SOBscore \rangle \simeq 1$. **B:** Two breast cancer FF samples with assumed genuine mutations and uncertain variants.

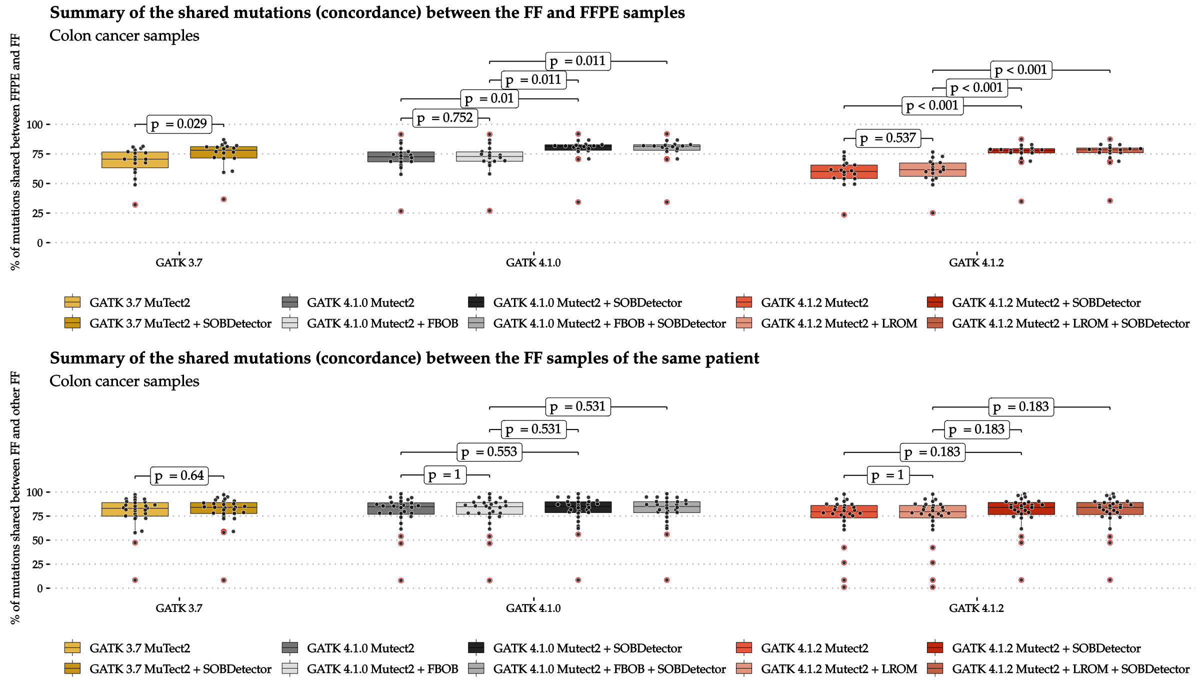

**Suppl.Fig. 10:** The shared ratio (described in the main text) between the matched FFPE and FF samples within the colon adenocarcinoma cohort. Below: The shared ratios between matched fresh-frozen sample pairs

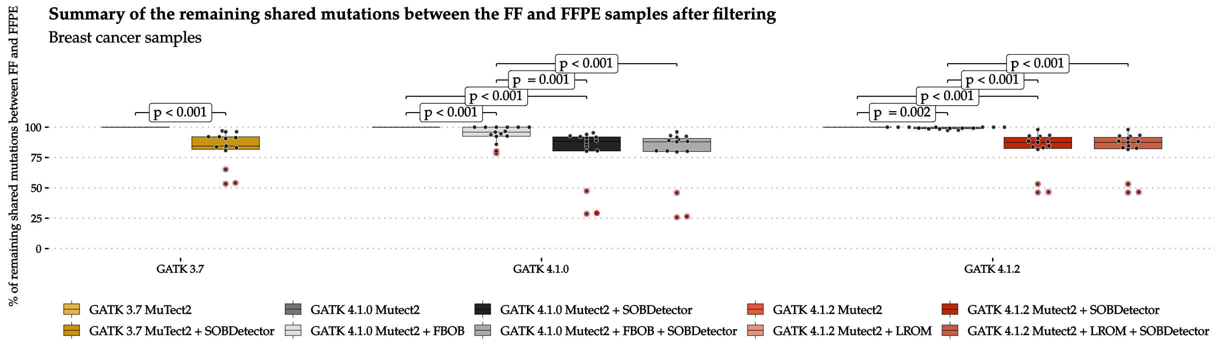

**Suppl.Fig. 11:** The remaining shared ratios (described in the main text) between the matched FFPE and FF samples within the breast invasive carcinoma cohort.

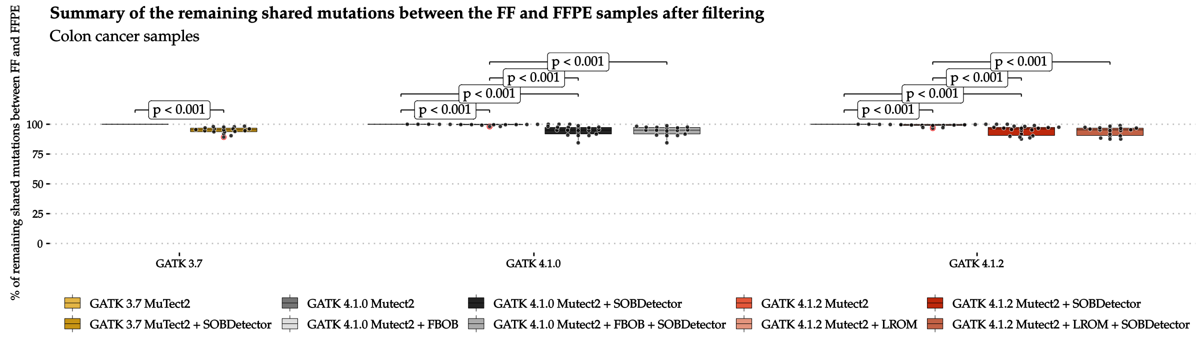

**Suppl.Fig. 12:** The remaining shared ratios (described in the main text) between the matched FFPE and FF samples within the colon adenocarcinoma cohort.

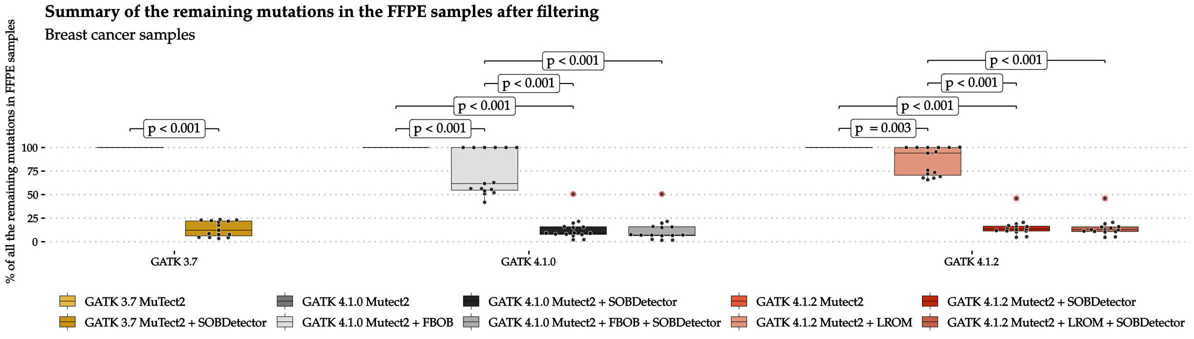

**Suppl.Fig. 13:** The remaining ratios of the original mutations between the matched FFPE and FF samples within the breast invasive carcinoma cohort.

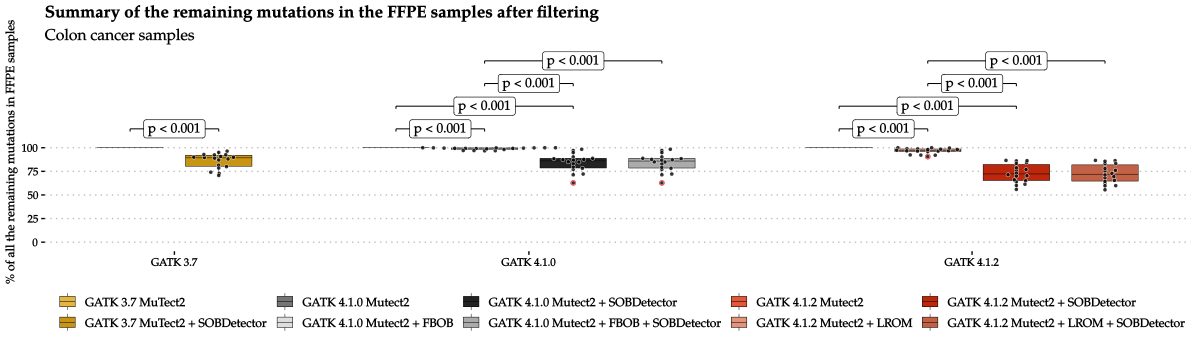

**Suppl.Fig. 14:** The remaining ratios the original mutations between the matched FFPE and FF samples within the colon adenocarcinoma cohort.

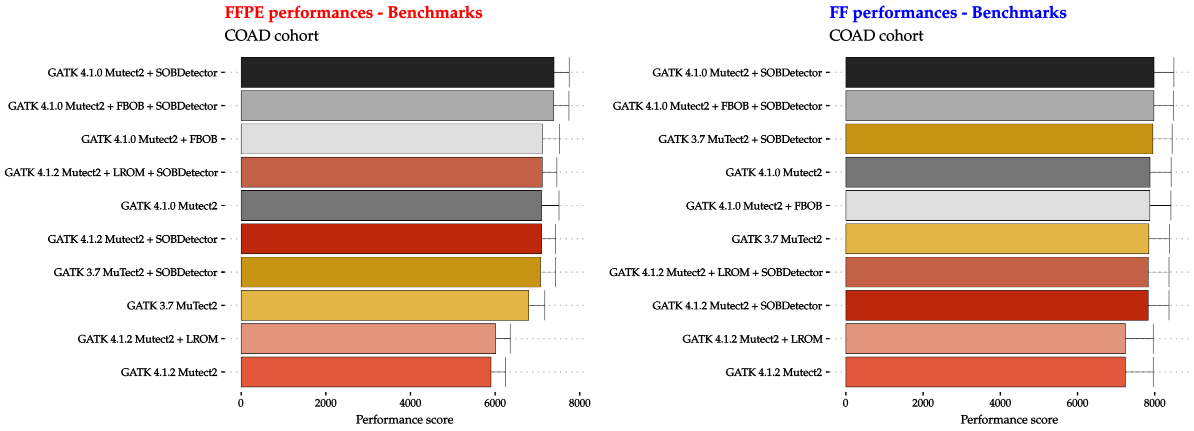

**Suppl.Fig. 15:** Performance scores oof the pipelines on the colon adenocarcinoma samples.

### 2 Case Studies

In order to evaluate SOBDetector on non-TCGA cancer samples, we have collected FFPE-FF matched tumor pairs from two previous studies:

- 4 samples from the Oh et al. study [1]
- 11 samples from the Van Allen et al. study [2]

In the following, the analysis of these samples will be illustrated.

#### 2.1 Oh et al. Study

Binary alignment files of the Oh et al. cohort were deposited at SRA, under accession number PRJNA301548. This dataset contains 13 whole exome sequences from 4 cancer patients:

- **Patient 1:** fibrosarcoma (1 FF normal, 1 FF tumor, 1 FFPE tumor)
- **Patient 2:** lung adenocarcinoma (1 FF normal, 1 FF tumor, 1 FFPE tumor)
- **Patient 3:** lung adenocarcinoma (1 FF normal, 1 FF tumor, 1 FFPE tumor)
- **Patient 4:** dermatofibrosarcoma protuberans (1 FF normal, 1 FF tumor, 2 FFPE tumor (one from 2007, one from 2009)

The exomes were realigned to grch37. A subsequent post-processing was performed, that included the removal of optical/PCR duplicates (`sambamba` [3]) and recalibration of base quality scores (`GATK BaseRecalibrator`). Since a significant difference in performance between the `GATK v4.1.2` and `GATK v4.1.1 Mutect2` pipelines could not be determined, somatic variants in this cohort were called using the `GATK v4.1.1` version only.

##### 2.1.1 Running Mutect2

In order of accuracy, the following codes will serve to show how the Mutect2 pipeline was run on these samples.

###### PileupSummary collection

These lines were run on all 13 whole exome bams.

```
java -jar GenomeAnalysisTK.jar GetPileupSummaries  \
     -I  tumor_location/sampleID.bam \
     -V  common_biallelic_exonic_variants \
     -O  output_path/sampleID_summary.table
```

```
java -jar GenomeAnalysisTK.jar CalculateContamination \
     -I output_path/sampleID_summary.table \
     -O output_path/sampleID_contamination.table
```

###### Somatic Mutation Calling

Since this cohort consisted of only 4 normal samples, a panel of normals was not created.

```
java -Xmx16g -jar GenomeAnalysisTK.jar Mutect2 \
     -R  reference_genome_location/grch37.fa \
     -I  germline_bam_location \
     -I  tumor_bam_location \
     -tumor  tumorSampleID \
     -normal germlineSampleID \
     --germline-resource germline_resource_path \
     --af-of-alleles-not-in-resource 0.0000025 \
     --disable-read-filter MateOnSameContigOrNoMappedMateReadFilter \
     -O output_vcf_path/tumorSampleID.vcf \
     -bamout reanalyzedBamLocation/tumorSampleID.bam
```

```
java -Xmx16g -jar GenomeAnalysisTK.jar FilterMutectCalls \
    -V output_vcf_path/tumorSampleID.vcf \
    --contamination-table contamination_table_path/sampleID_contamination.table \
    -O output_vcf_path/tumorSampleID_filtered.vcf
```

##### Strand Orientation Bias filtering

Collecting the sequencing artifact metrics files:

```
java -Xmx8g -jar GenomeAnalysisTK.jar CollectSequencingArtifactMetrics \
    -I tumor_bam_location/tumorSampleID.bam \
    -O tumor_bam_location/tumorSampleID_artifact \
    --FILE_EXTENSION ".txt" \
    -R reference_genome_location/grch37.fa
```

Filtering using FilterByOrientationBias (both for oxidative and ffpe artifacts)

```
java -Xmx16g -jar GenomeAnalysisTK.jar FilterByOrientationBias \
    --artifact-modes G/T \
    --artifact-modes C/T \
    -V output_vcf_path/tumorSampleID_filtered.vcf \
    -P tumor_bam_location/tumorSampleID_artifact.pre_adapter_detail_metrics.txt \
    -O output_vcf_path/tumorSampleID_filtered_sobf.vcf
```

##### Using SOBDetector

SOBDetector was used using the non-strand orientation bias filtered vcfs
(`output_vcf_path/tumorSampleID_filtered.vcf`),
and the reanalyzed (i.e. at certain loci realigned) bam files
(`reanalyzedBamLocation/tumorSampleID.bam`).

```
java -jar SOBDetector_v1.0.jar \
    --input-type VCF \
    --input-variants ./input-variants.vcf \
    --input-bam ./input.bam \
    --sample-name SAMPLE1 \
    --output-variants ./output.vcf \
    --only-passed true
```

##### Determining the standardization parameters of the cohort

Variants from the SOBDetector filtered vcfs were converted into tab-delimited table files using `GATK VariantsToTable`, then these tables were merged into a single data frame using R. After filtering on "PASS"-ed variants, the following additional hard-filters were applied on them:

TUMOR.depth >= 20 & NORMAL.depth >= 10 & TUMOR.AF >= 0.05 & NORMAL.AF == 0.

Using the remaining mutations, the means and standard deviations of the four SOBDetector attributes were calculated, and saved into a text file: `OhStandardization.txt`, with the following contents:

```
TUMOR.ALT   TUMOR.depth  TUMOR.AF   SOBScore
6.266954  40.88039  0.1725151  0.7898631
6.42966  28.36066  0.1127767  0.3091478
```

The first line contains the attribute names (the order is important), the second the means of each attribute, and the third their standard deviations. In order to increase the predictive accuracy, SOBDetector then was rerun on the vcf/tumor samples just as before, but with the additional
`-standardization-parameters OhStandardization.txt`
parameter. The final vcfs then were processed again in R.

##### 2.1.2 Summary of the attributes

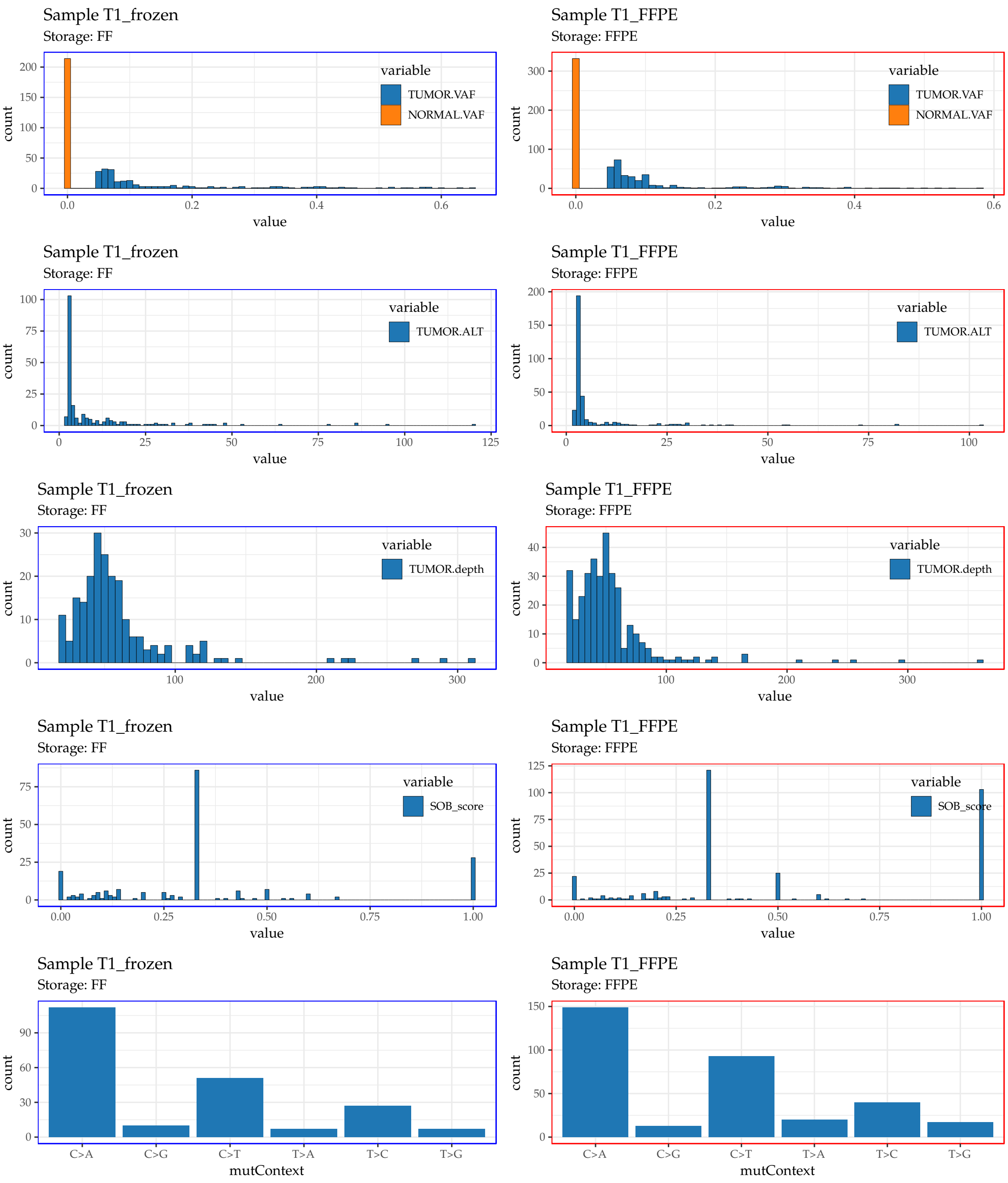

**Suppl.Fig. 16:** Variant attributes of Oh et al. Patient 1. Distributions belonging to frozen samples have blue frames around them, while those that belong the formalin fixed samples surrounded by red frames.

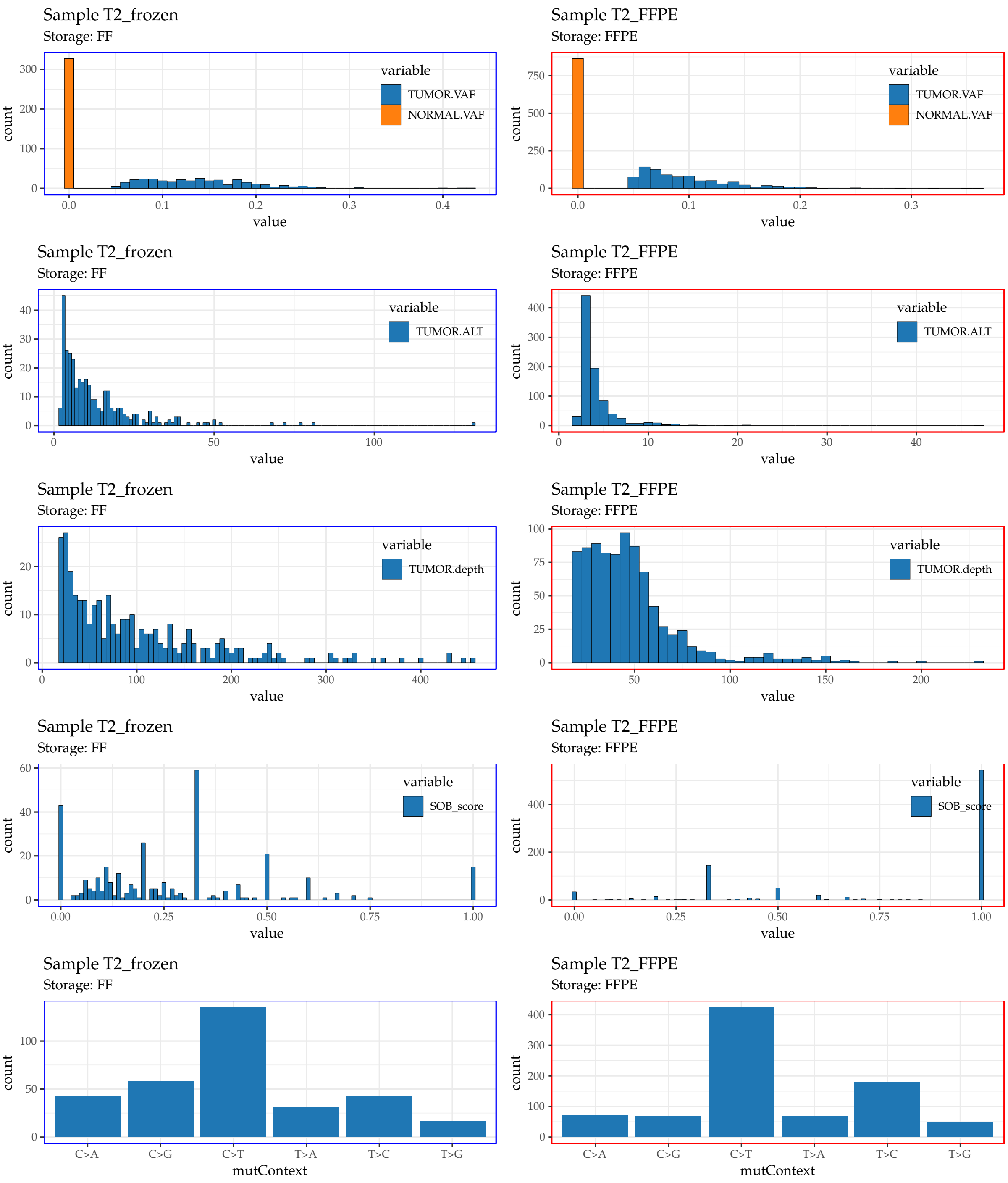

**Suppl.Fig. 17:** Variant attributes of Oh et al. Patient 2. Distributions belonging to frozen samples have blue frames around them, while those that belong the formalin fixed samples surrounded by red frames.

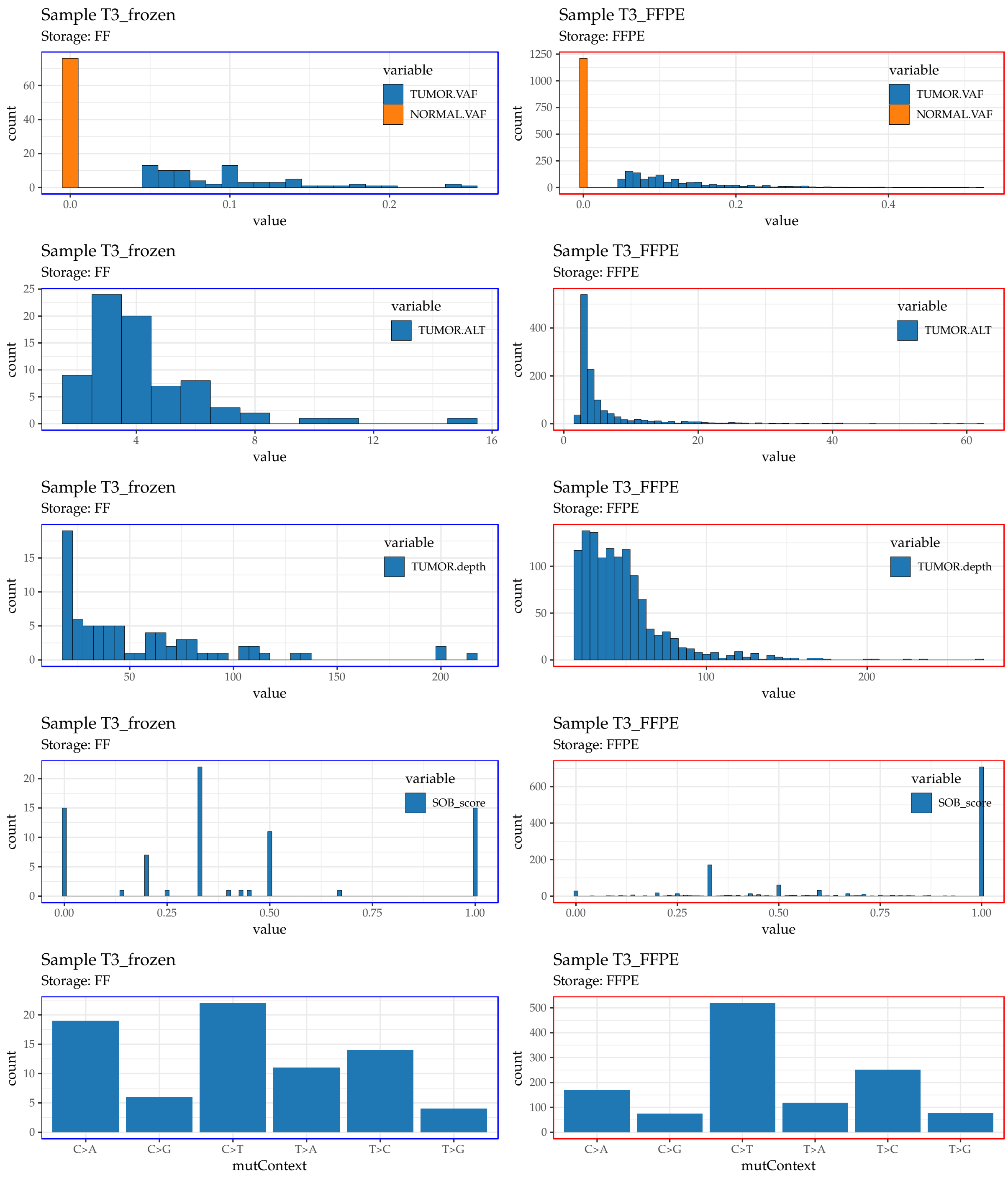

**Suppl.Fig. 18:** Variant attribute of Oh et al. Patient 3. Distributions belonging to frozen samples have blue frames around them, while those that belong the formalin fixed samples surrounded by red frames.

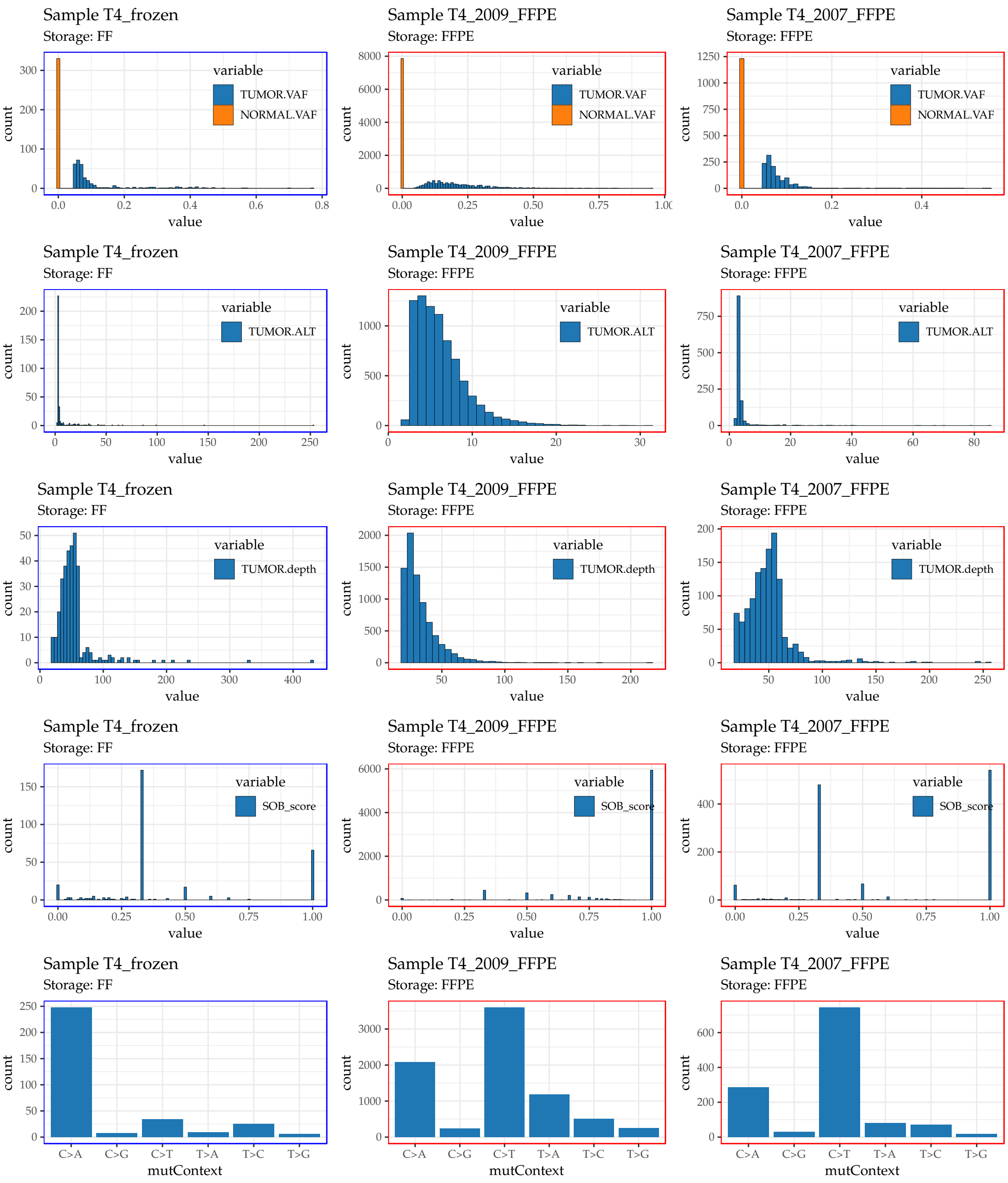

**Suppl.Fig. 19:** Variant attributes of Oh et al. Patient 4. Distributions belonging to frozen samples have blue frames around them, while those that belong the formalin fixed samples surrounded by red frames.

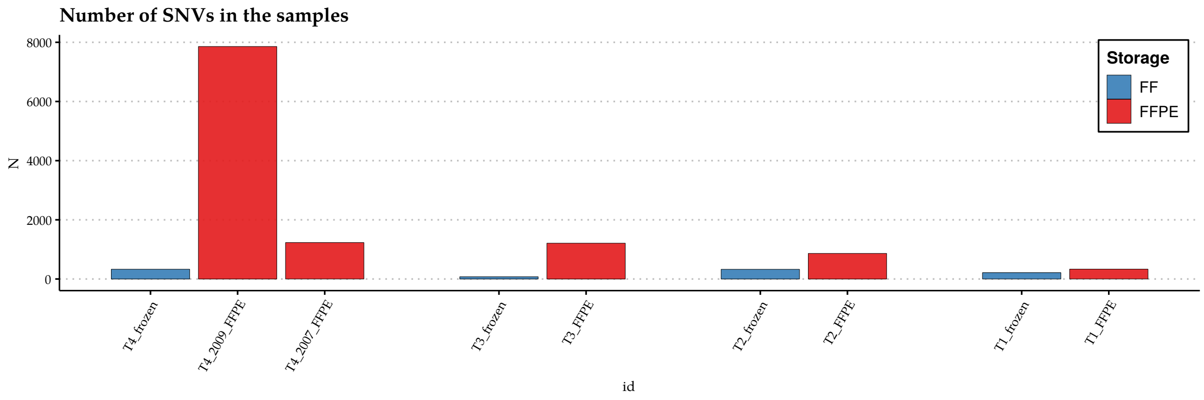

**Suppl.Fig. 20:** Number of mutations in the Oh et al samples, after filtering (both initially by Mutect2, and by applying the additional hard filters).

##### 2.1.3 Defining likely artifacts and likely genuine mutations

The definitions of the likely artifact and likely genuine mutation classes are less strict in this cohort, due to the smaller number of available samples per patient, than in the TCGA.

- **likely artifact**: variants that are only present in the FFPE tumor ($N_{\text{FFPE only}} = 11775$)
- **likely genuine mutations**: variants that are present in the FF and FFPE tumors ($N_{\text{shared}} = 410$)

**Remark:** the likely artifact group here purely means that a mutation is solely present in the FFPE sample! Many of these might be genuine mutations, which were not detected in the FF specimen due to intratumor heterogeneity. Others might be sequencing artifacts, without strand orientation bias. This is also visible on the distribution of the SOB scores of the FFPE samples:

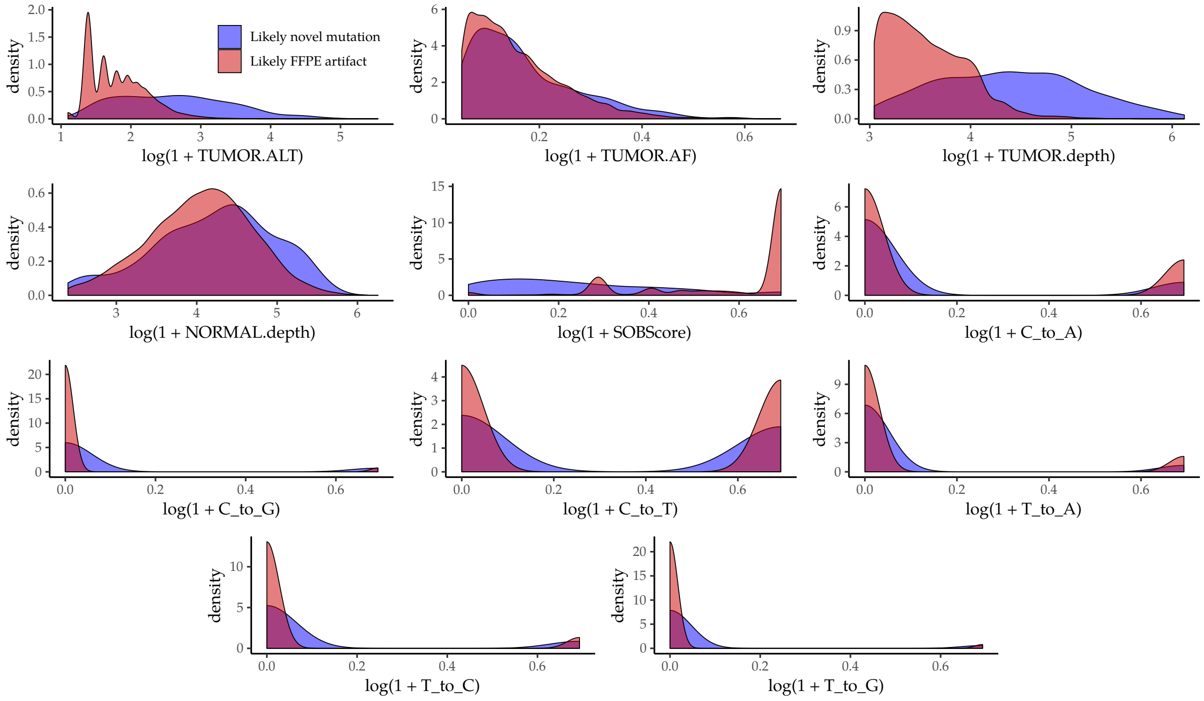

**Suppl.Fig. 21:** Log-transformed and standardized densities of the attributes extracted from the Oh et al. samples.

In order to evaluate the model, 400 likely genuine mutations (i.e. mutations shared by the corresponding FF and FFPE samples) and 600 likely artifacts were collected randomly into an evaluations set.

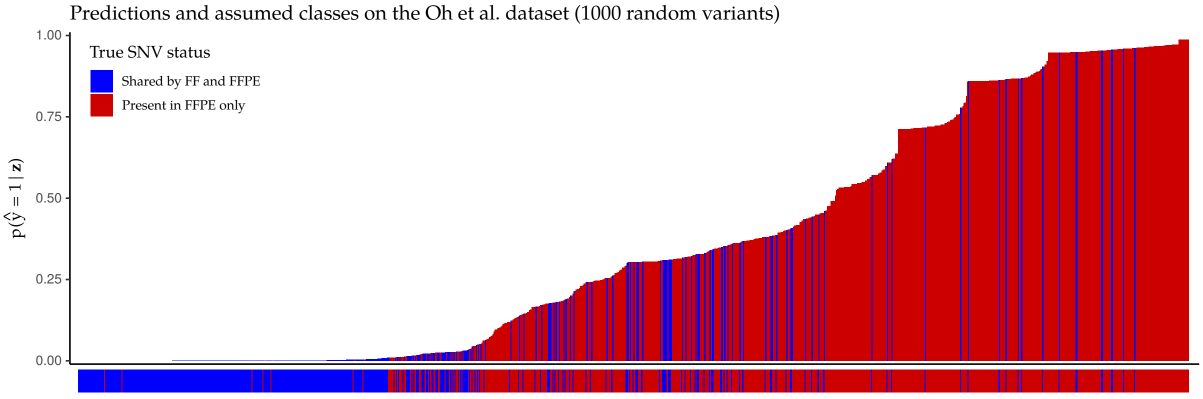

**Suppl.Fig. 22:** Predicted $P(\text{variant is artifact}|\text{attributes})$ SOBDetector probabilites of 1000 randomly samples variants from the Oh et al. dataset.

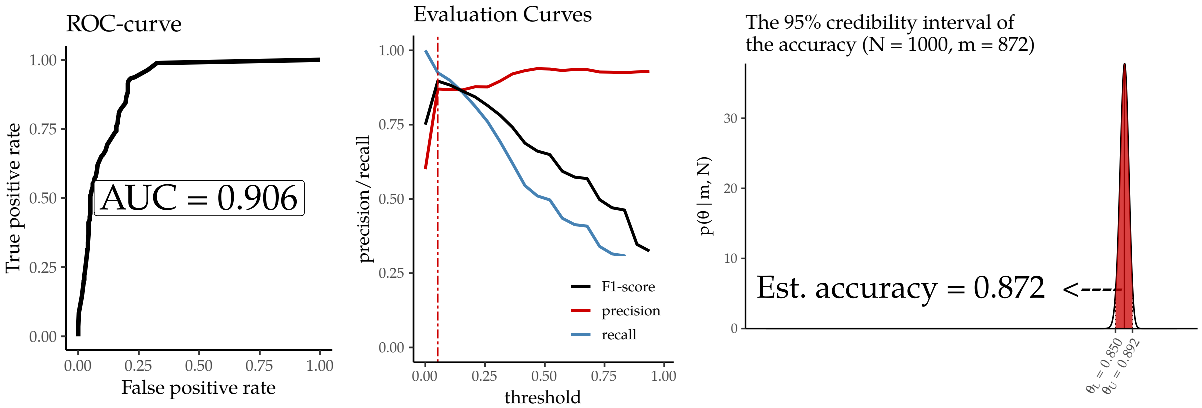

**Suppl.Fig. 23:** Evaluation curves and predicted accuracy - Oh et al. dataset.

Using the threshold where the harmonic mean of the recall and precision were at its maximum, the estimated accuracy is $0.872 \pm 0.02$. While this estimate is lower than the accuracy measured on the TCGA evaluation set, it is likely due to the many misclassified likely artifacts.

##### 2.1.4 Sample-level results

The following figures will show the results of a SOBDetector filtering on a sample-level. The figures contain sorted barplots of the SOBDetector prediction scores, with their corresponding SOB scores, tumor allele frequencies, number of variant supporting reads and coverage at their locus in the tumor. The FilterByOrientation (Mutect2) predictions are also indicated: white color corresponds to predicted genuine mutations, black to likely artifacts. The performance of the the artifact removal strategies is compared analogously to the main text: the yellow/blue/grey barplots show the shared and remaining_shared ratios between the FFPE (file1) and FF samples (file2), along with their performance scores.

#### Patient 1:

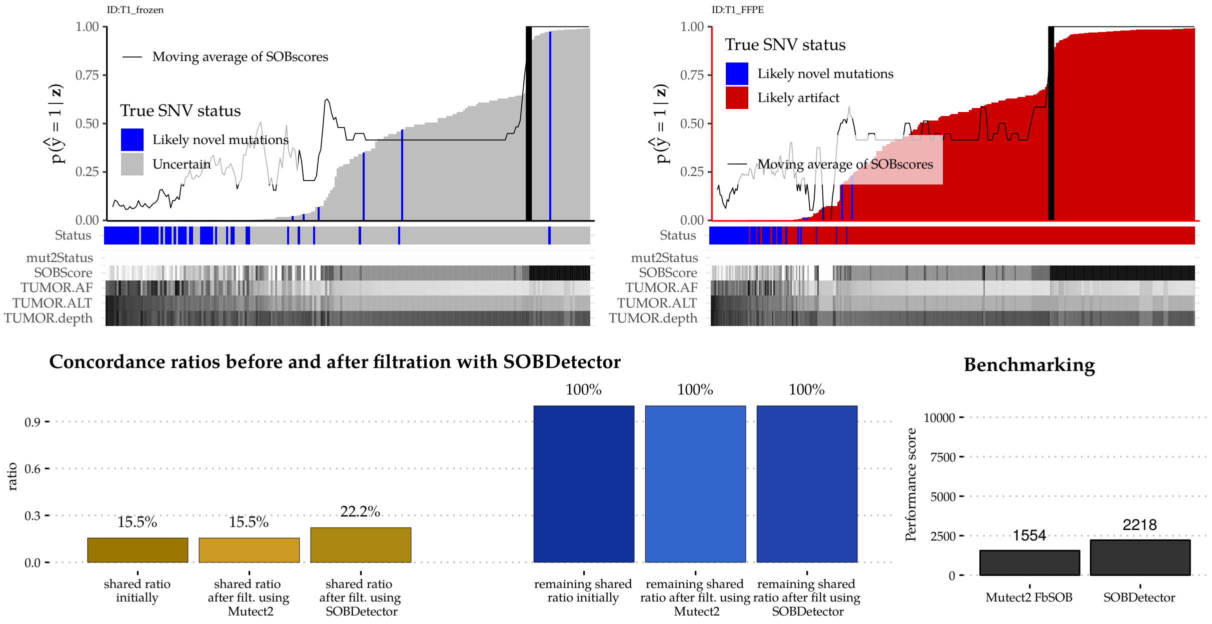

**Suppl.Fig. 24:** Sample-level results of Patient1.

#### Patient 2:

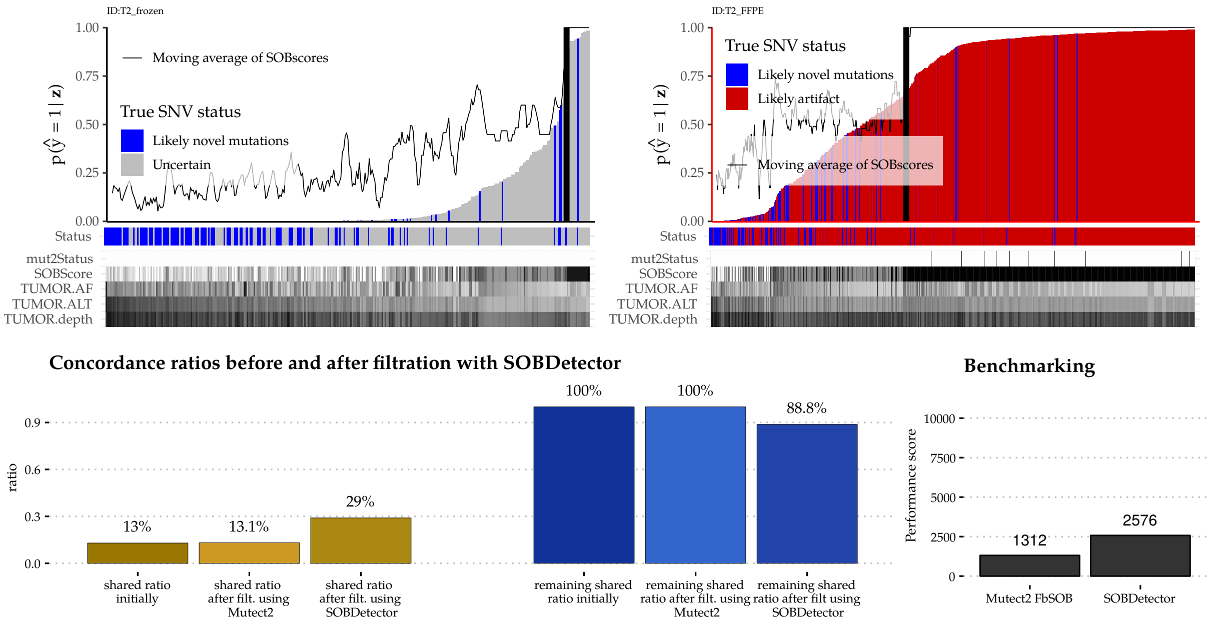

**Suppl.Fig. 25:** Sample-level results of Patient2.

#### Patient 3:

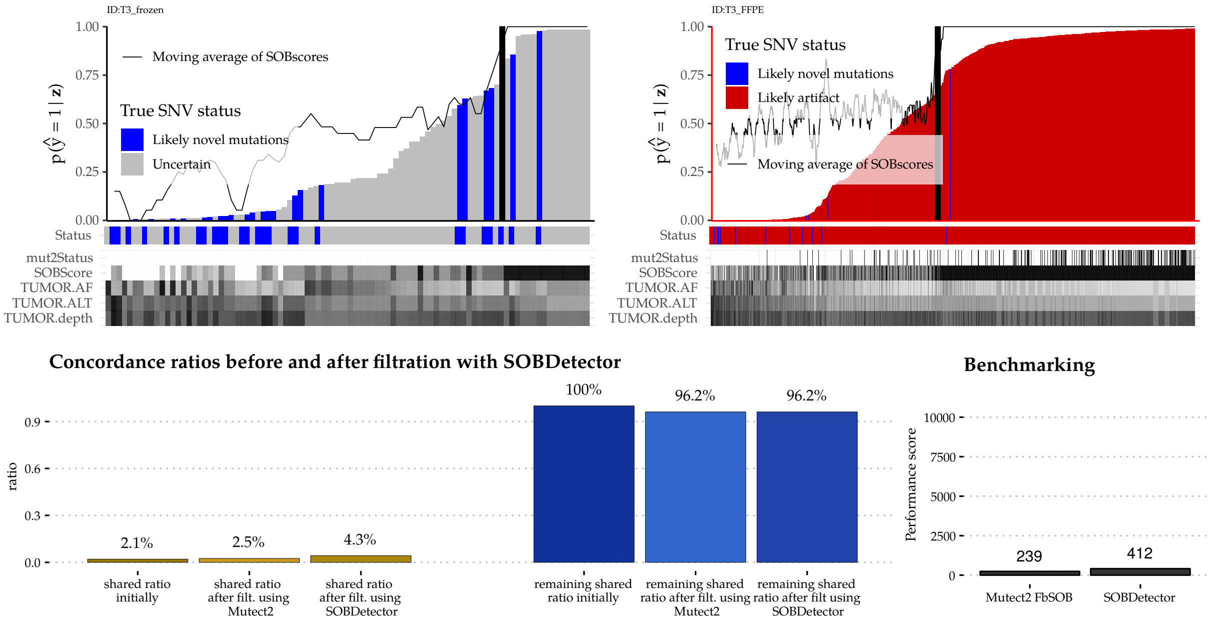

**Suppl.Fig. 26:** Sample-level results of Patient3.

#### Patient 4:

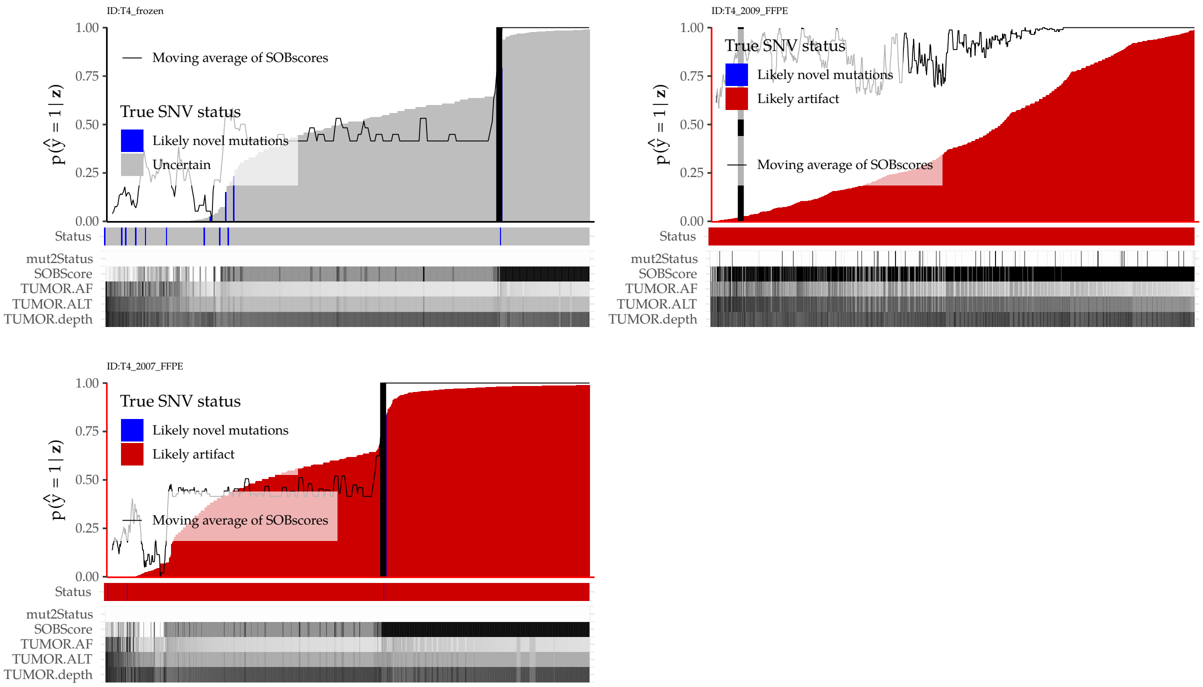

/continues on next page/

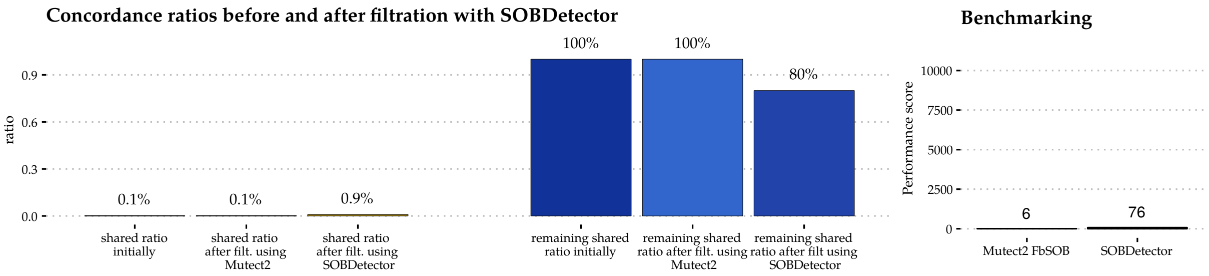

**Suppl.Fig. 27:** Sample-level results of Patient4.

##### 2.1.5 Conclusion

SOBDetector has managed to differentiate between the majority of the likely genuine mutations and likely artifact in the Oh et al. cohort. The FFPE damage however in these samples was high, only a small number of mutations were shared between the corresponding pairs, and the formalin fixed specimens had more filter-passing somatic SNV in general. The GATK strand orientation bias filter performed worse than SOBDetector in all 4 patient cases. In addition, the fresh frozen samples had mutation that exhibited strand orientation bias, most of which was not shared with the FFPE. This suggests, that a non-formalin-induced damage, such as oxidation has also affected this cohort.

#### 2.2 Van Allen et al. Study

Binary alignment files of the Van Allen cohort were deposited at dbGaP, under accession number phs000488.v2.p1. This dataset contains (hybrid captured) whole exome sequences from 208 lung adenocarcinoma patients, from whom "only" **11 had matched FFPE-FF tumor samples extracted from the same tumor**. The patient IDs used in this case-study follow the notation of the original work.

The 11 patients were identified from the supplementary materials of the article. Each patient has four samples, 1 FF normal, 1 FFPE normal, 1 FF tumor and 1 FFPE tumor. The dbGaP "Run" table, however, marked both FFPE samples as tumors, hence we have decided to run every sample against the FF normal, and call somatic mutations from them, using Mutect2 (`GATK 4.1.1`). By following this strategy, the FFPE normal cases could be easily identified, and all the filter passing mutations would be false positive mutations calls.

The exomes were realigned to grch37. A subsequent post-processing was performed, that included the removal of optical/PCR duplicates (`sambamba` [3]) and recalibration of base quality scores (`GATK BaseRecalibrator`). Since a significant difference in performance between the `GATK v4.1.2` and `GATK v4.1.1 Mutect2` pipelines could not be determined, somatic variants in this cohort were called using the `GATK v4.1.1` version only.

##### 2.2.1 Running Mutect2

Somatic variant discovery followed the same workflow as in the Oh et al. case study. The only difference is, that by using the 11 normal FF samples, a panel of normals was created, and used for accounting for systematic sequencing artifacts.

###### Determining the standardization parameters of the cohort

Variants from the SOBDetector filtered vcfs were converted into tab-delimited table files using `GATK VariantsToTable`, then these tables were merged into a single data frame using R. After filtering on "PASS"-ed variants, the following additional hard-filters were applied on them:

TUMOR.depth >= 20 & NORMAL.depth >= 10 & TUMOR.AF >= 0.05 & NORMAL.AF == 0.

Using the remaining mutations, the means and standard deviations of the four SOBDetector attributes were calculated, and saved into a text file: `AllenStandardization.txt`, with the following contents:

```
TUMOR.ALT   TUMOR.depth TUMOR.AF    SOBScore
17.93467  95.25824  0.1966972  0.2933799
21.90313  84.11611  0.1355127  0.2688528
```

The first line contains the attribute names (the order is important), the second the means of each attribute, and the third their standard deviations. In order to increase the predictive accuracy, SOBDetector then was rerun on the vcf/tumor samples just as before, but with the additional
`-standardization-parameters OhStandardization.txt`
parameter. The final vcfs then were processed again in R.

##### 2.2.2 Summary of the attributes

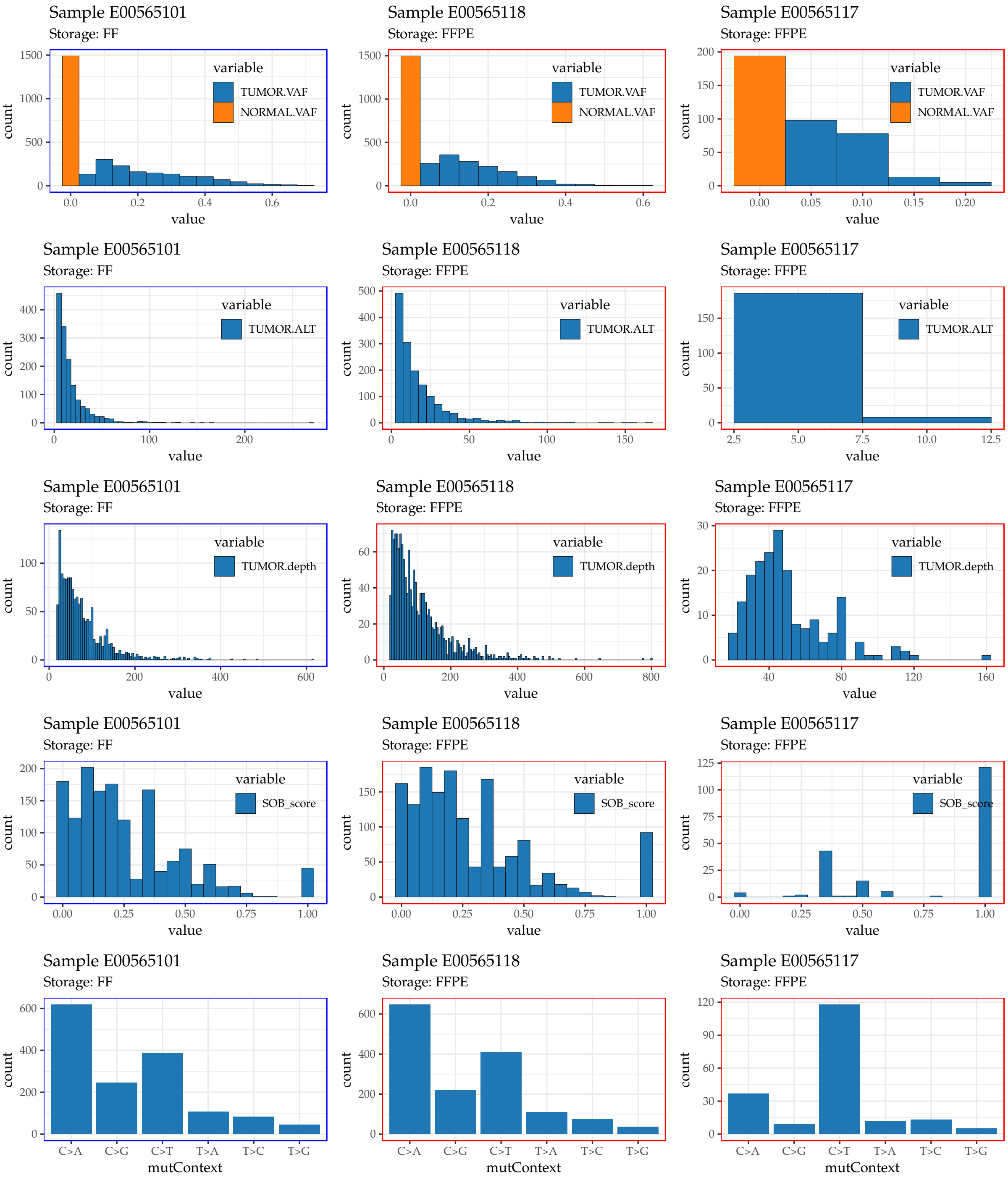

**Suppl.Fig. 28:** Variant attributes of Van Allen et al. Patient E00565. Distributions belonging to frozen samples have blue frames around them, while those that belong the formalin fixed samples surrounded by red frames.

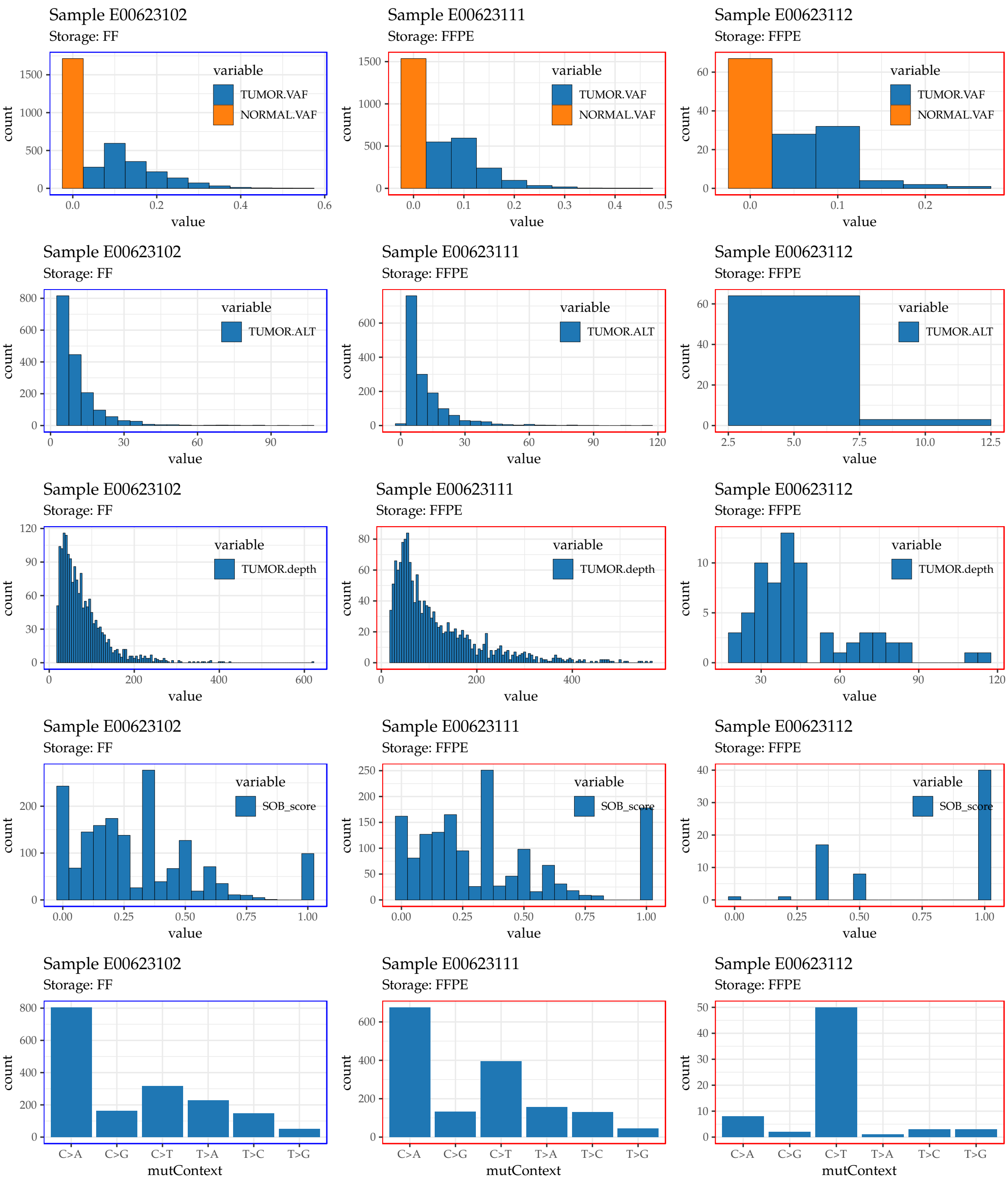

**Suppl.Fig. 29:** Variant attributes of Van Allen et al. Patient E00623. Distributions belonging to frozen samples have blue frames around them, while those that belong the formalin fixed samples surrounded by red frames.

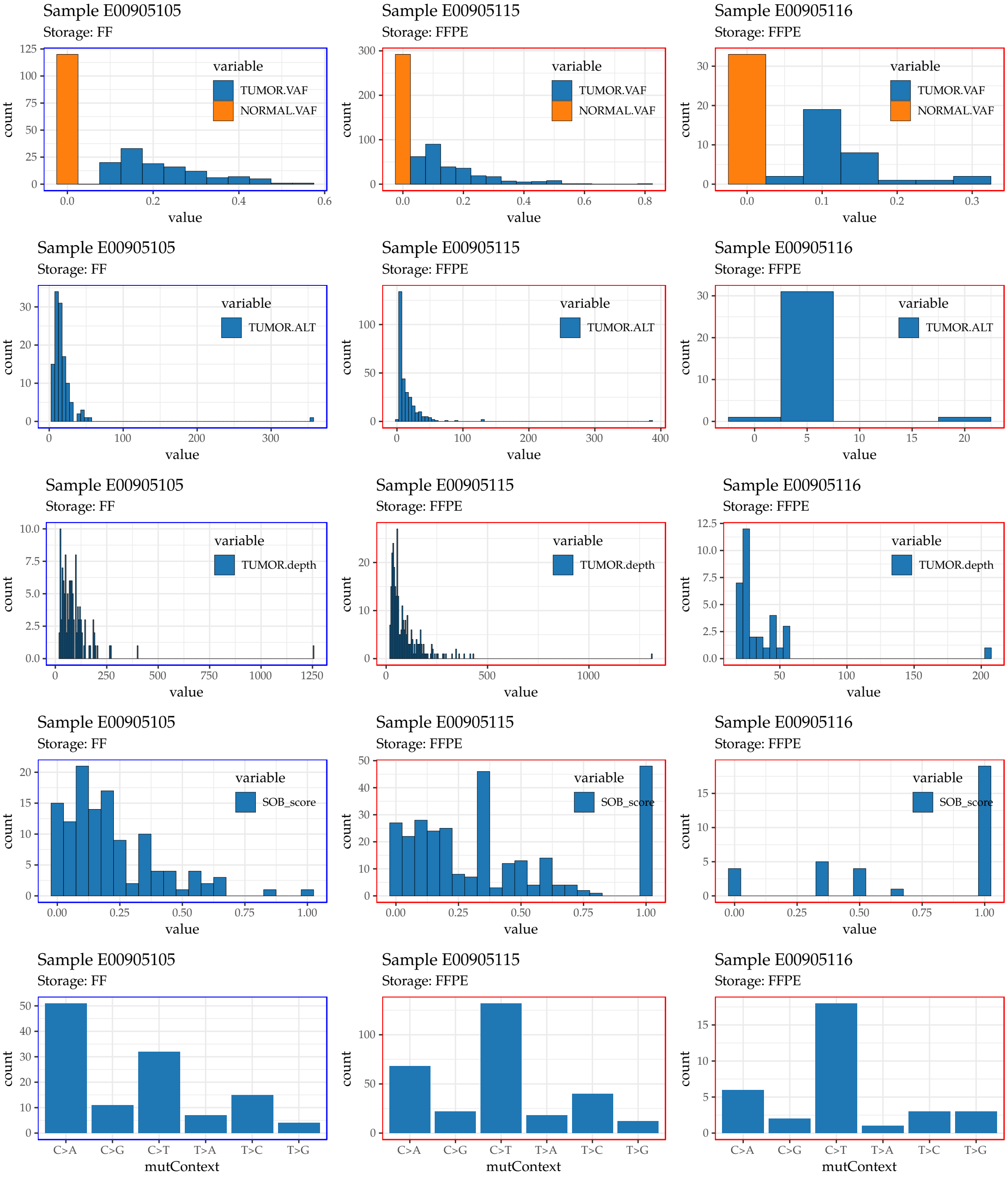

**Suppl.Fig. 30:** Variant attributes of Van Allen et al. Patient E00905. Distributions belonging to frozen samples have blue frames around them, while those that belong the formalin fixed samples surrounded by red frames.

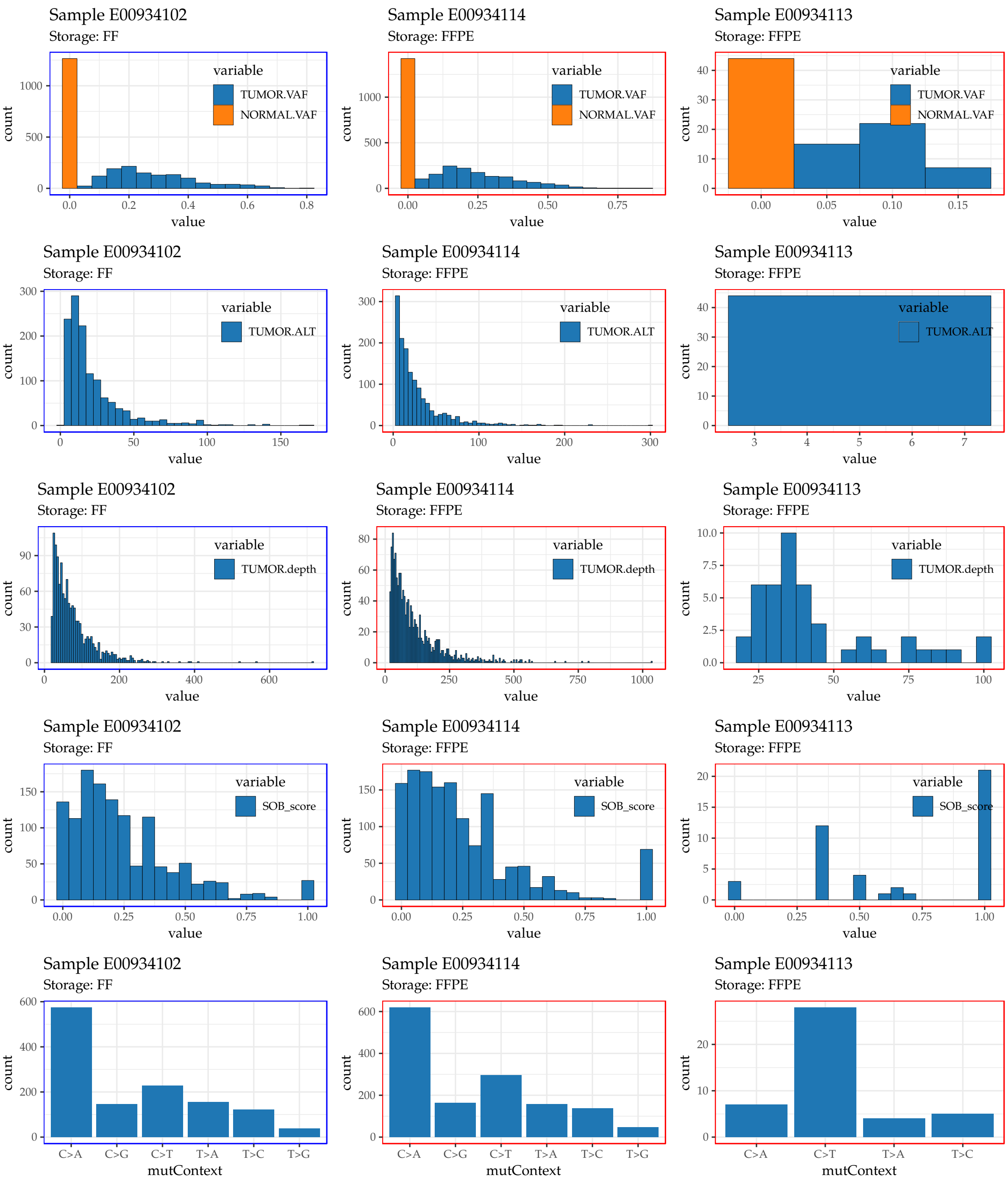

**Suppl.Fig. 31:** Variant attributes of Van Allen et al. Patient E00934. Distributions belonging to frozen samples have blue frames around them, while those that belong the formalin fixed samples surrounded by red frames.

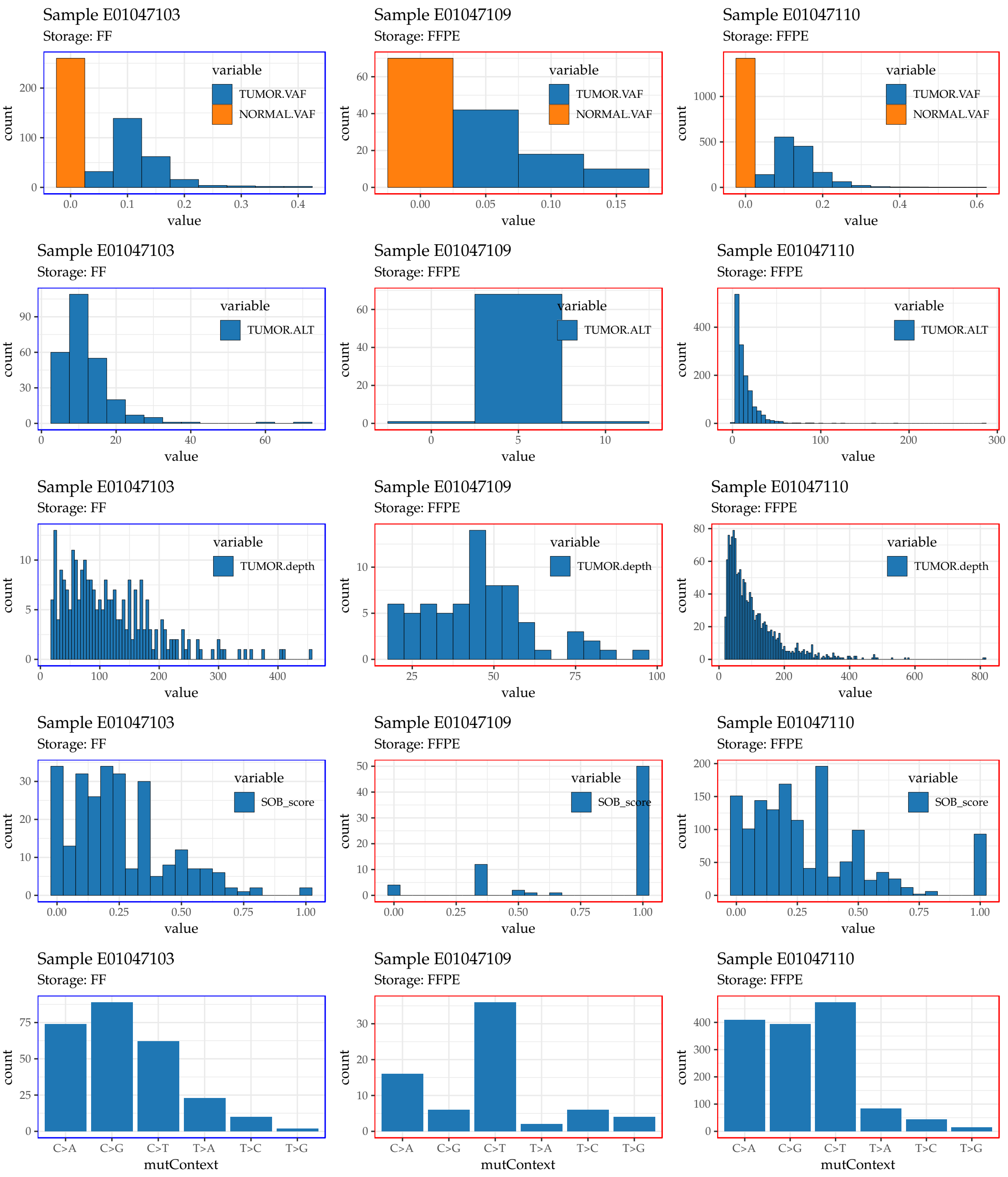

**Suppl.Fig. 32:** Variant attributes of Van Allen et al. Patient E01047. Distributions belonging to frozen samples have blue frames around them, while those that belong the formalin fixed samples surrounded by red frames.

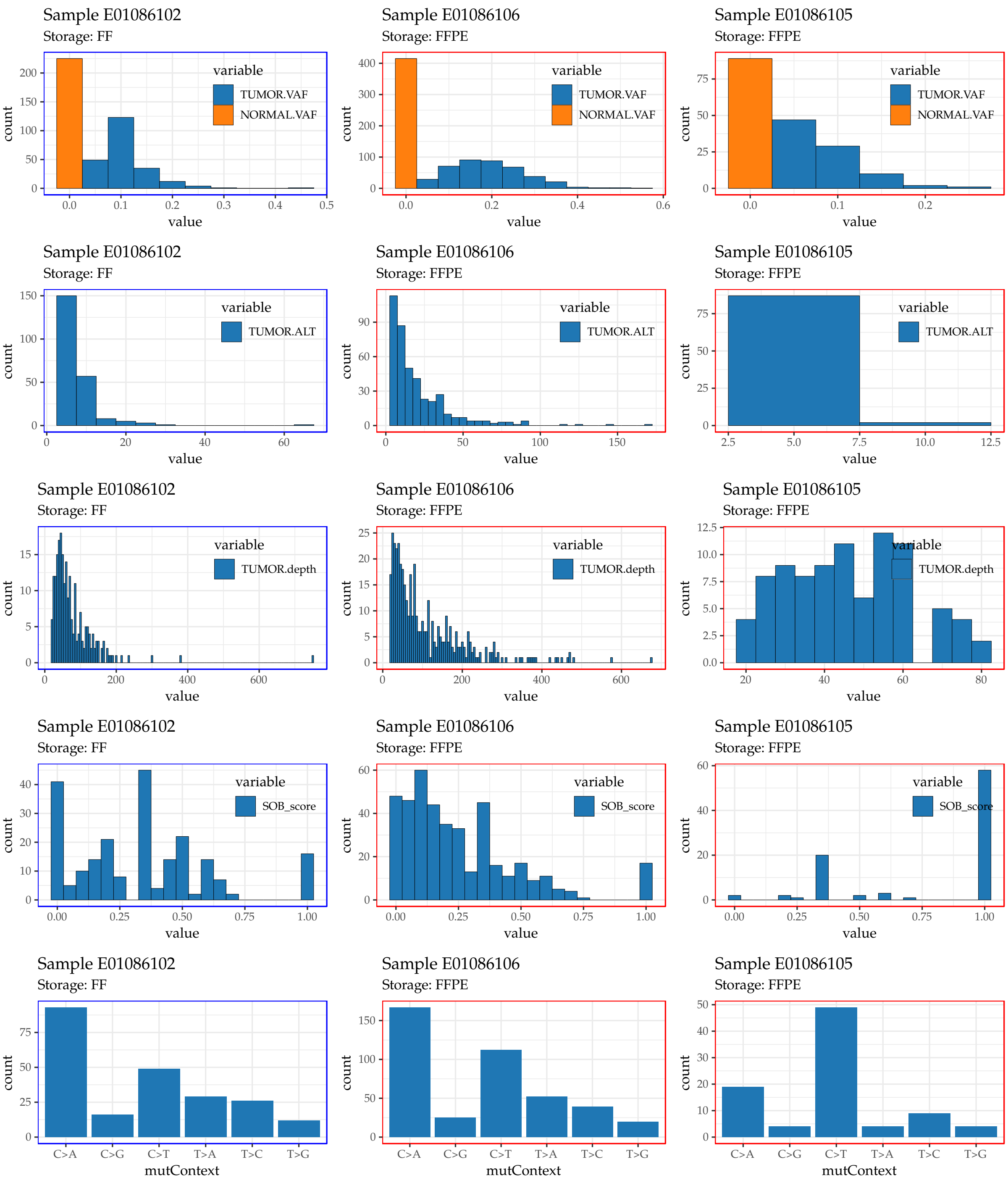

**Suppl.Fig. 33:** Variant attributes of Van Allen et al. Patient E01086. Distributions belonging to frozen samples have blue frames around them, while those that belong the formalin fixed samples surrounded by red frames.

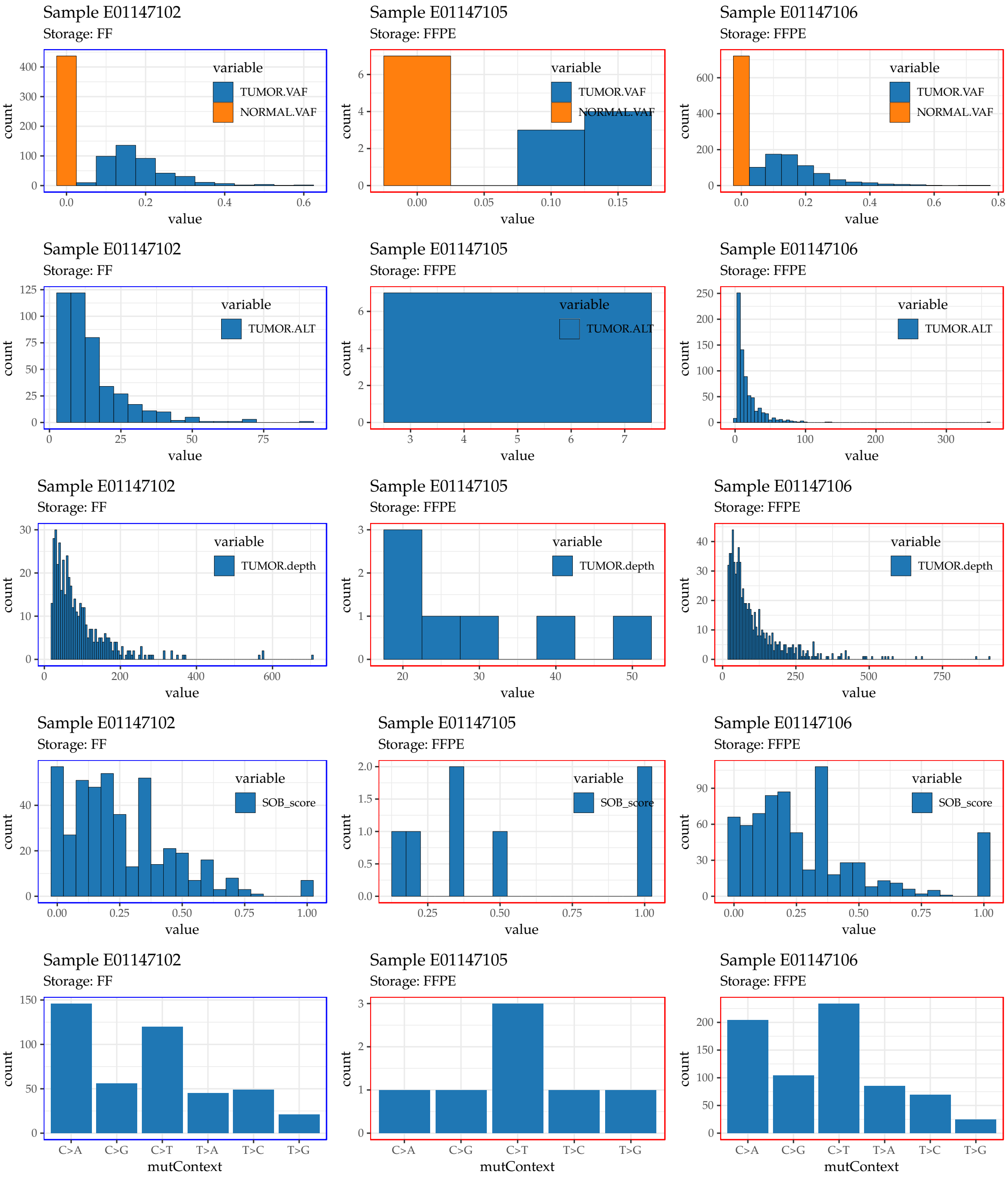

**Suppl.Fig. 34:** Variant attributes of Van Allen et al. Patient E01147. Distributions belonging to frozen samples have blue frames around them, while those that belong the formalin fixed samples surrounded by red frames.

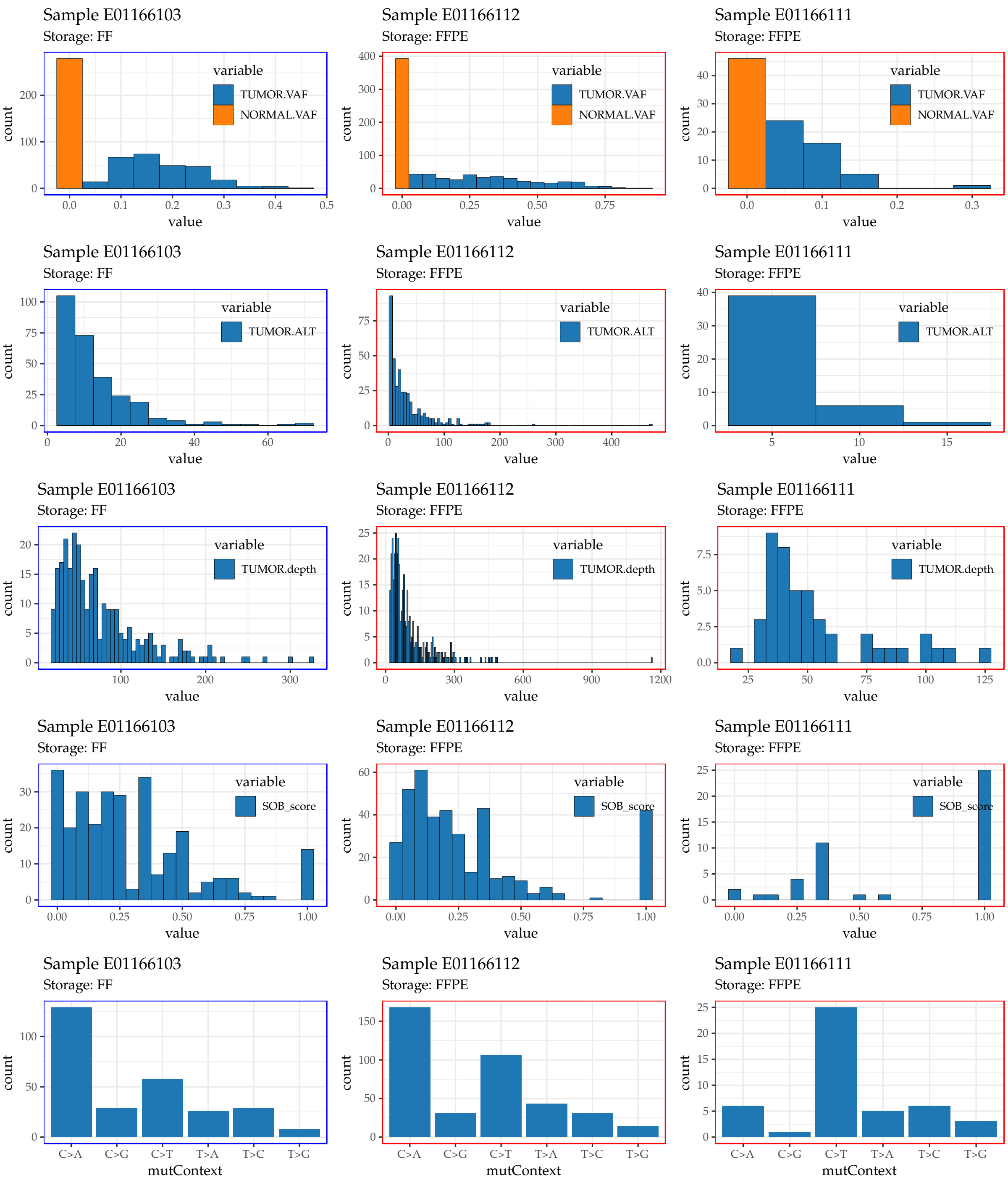

**Suppl.Fig. 35:** Variant attributes of Van Allen et al. Patient E01166. Distributions belonging to frozen samples have blue frames around them, while those that belong the formalin fixed samples surrounded by red frames.

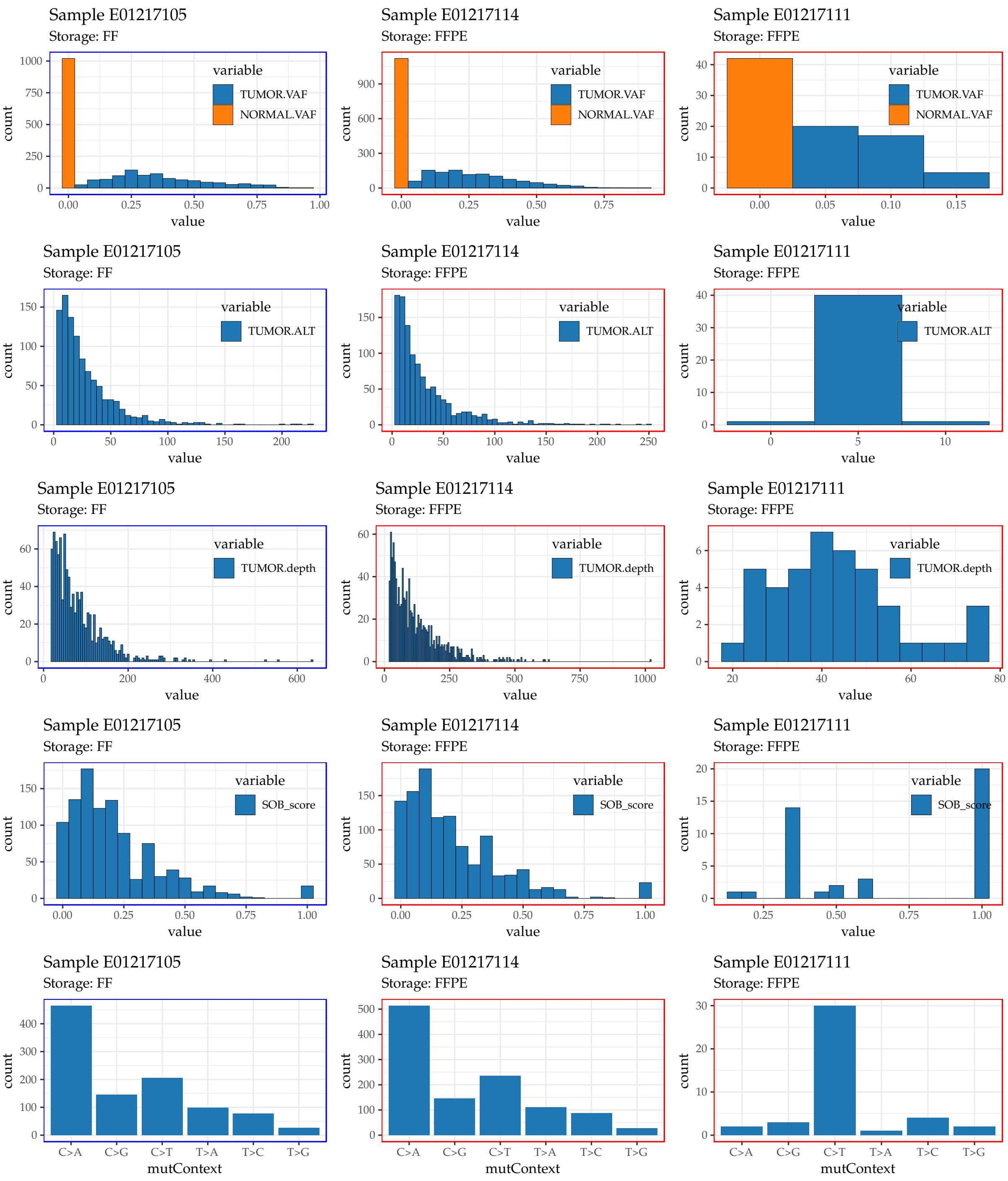

**Suppl.Fig. 36:** Variant attributes of Van Allen et al. Patient E01217. Distributions belonging to frozen samples have blue frames around them, while those that belong the formalin fixed samples surrounded by red frames.

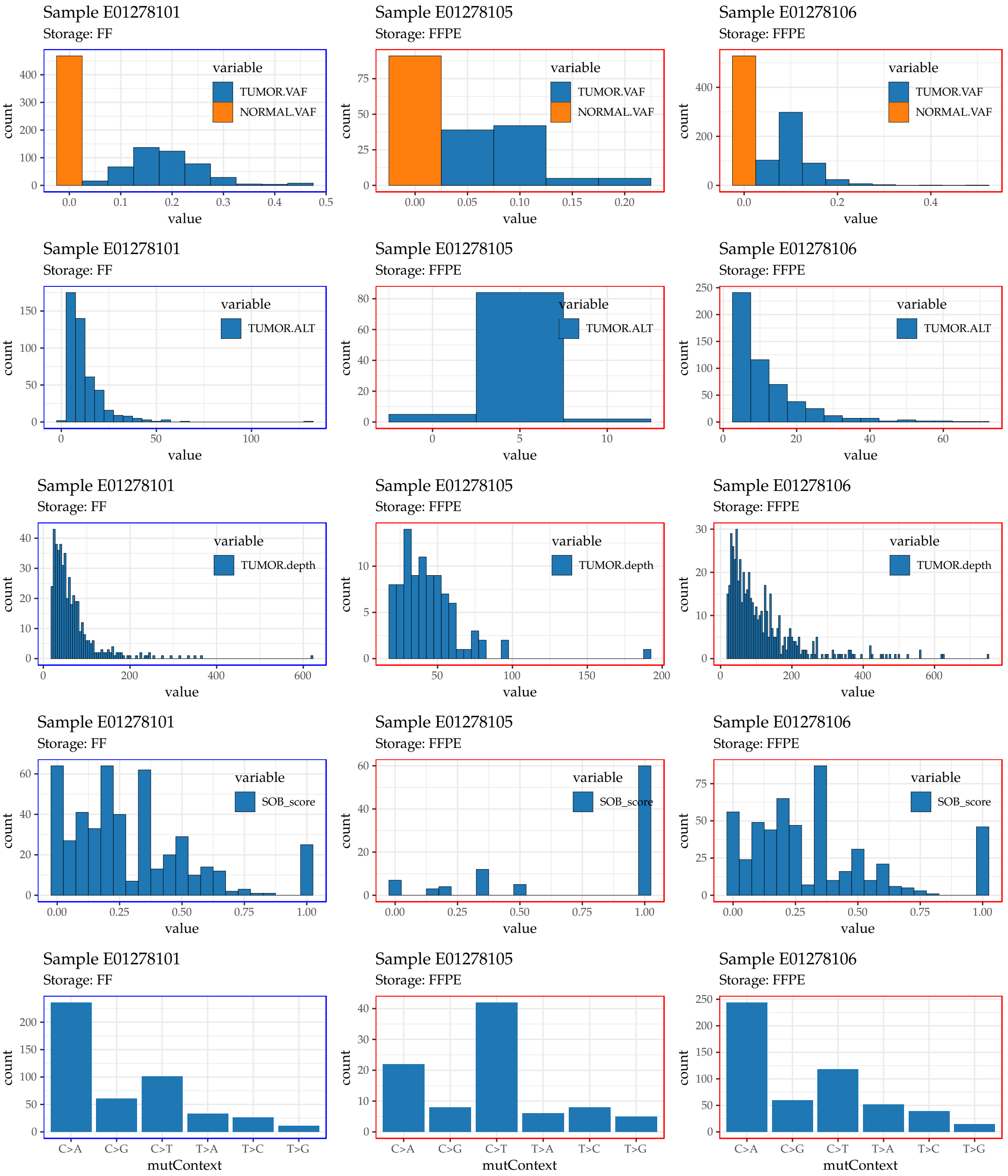

**Suppl.Fig. 37:** Variant attributes of Van Allen et al. Patient E01278. Distributions belonging to frozen samples have blue frames around them, while those that belong the formalin fixed samples surrounded by red frames.

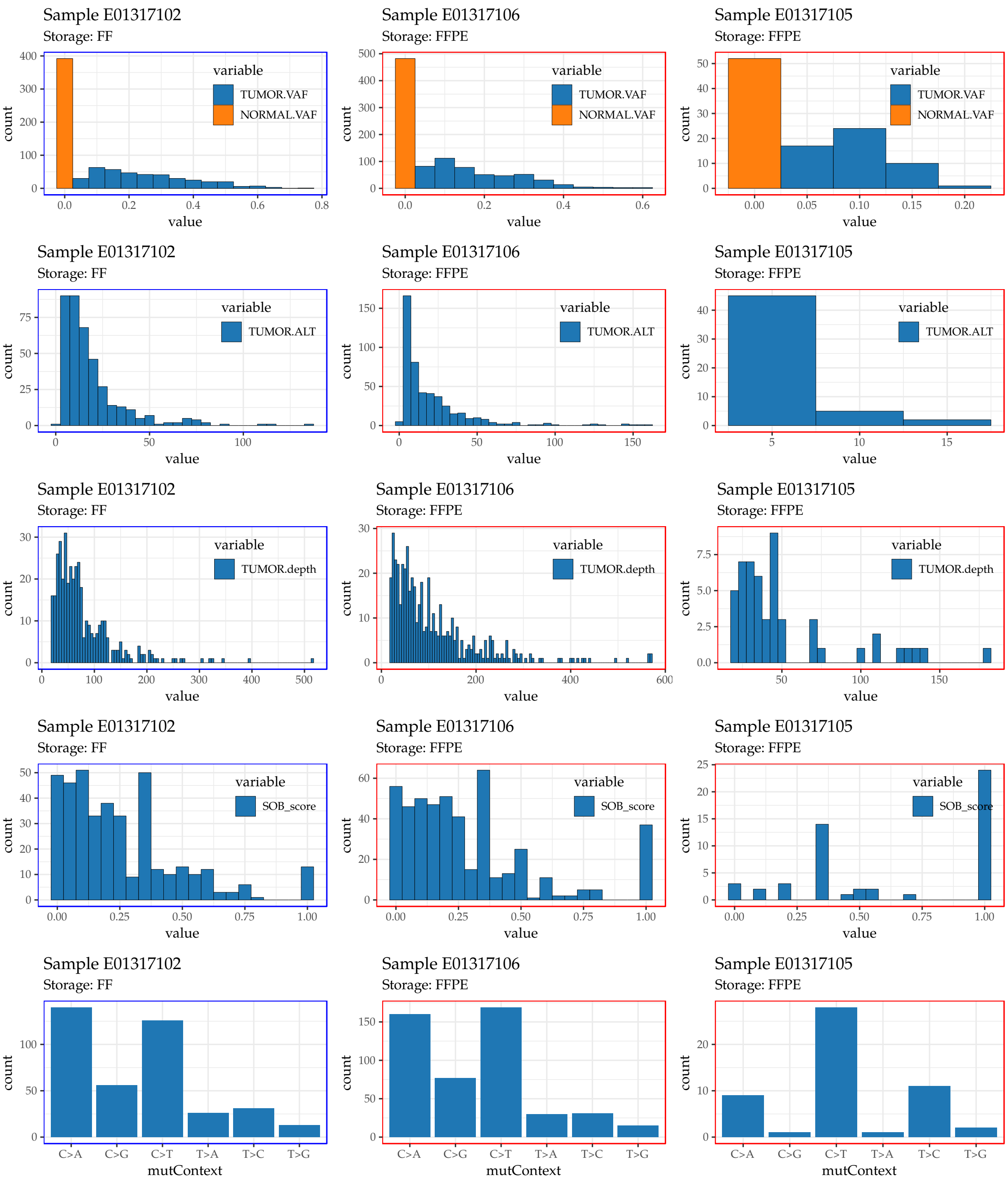

**Suppl.Fig. 38:** Variant attributes of Van Allen et al. Patient E01317. Distributions belonging to frozen samples have blue frames around them, while those that belong the formalin fixed samples surrounded by red frames.

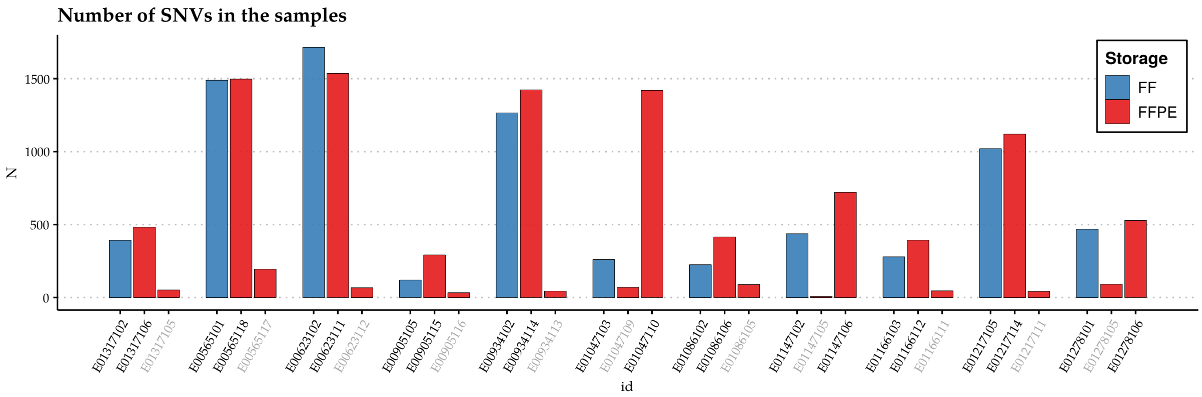

**Suppl.Fig. 39:** Number of mutations in the Van Allen et al. samples, after filtering (both initially by Mutect2, and by applying the additional hard filters). Based on the final amount of mutations, normal FFPE samples were identified and marked with grey sample names in the plot.

##### 2.2.3 Defining likely artifacts and likely genuine mutations

The definitions of the likely artifact and likely genuine mutation classes are even less strict in this cohort they they were in the Oh et al. case. **The FFPE samples had 1.5-2 times better coverage than their FF counterparts**, meaning that many clonal mutations with small allele frequencies could be captured in them, that did not appear in the FF samples. The likely artifact group therefore might contain more genuine mutations than real artifacts in this study.

- **likely artifact**: variants that are only present in the FFPE tumor ($N_{\text{FFPE only}} = 4702$)
- **likely genuine mutations**: variants that are present in the FF and FFPE tumors ($N_{\text{shared}} = 11720$)

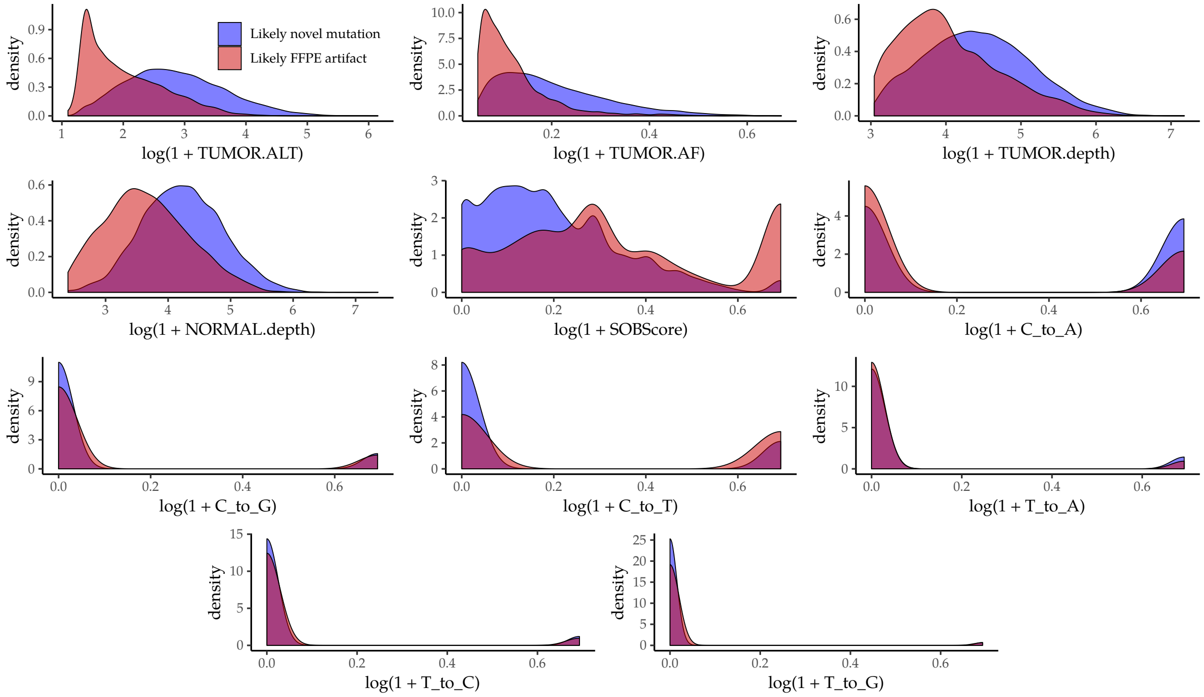

**Suppl.Fig. 40:** Log-transformed and standardized densities of the attributes extracted from the Van Allen et al. samples.

In order to evaluate the model, 500 likely genuine mutations (i.e. mutations shared by the corresponding FF and FFPE samples) and 500 likely artifacts were collected randomly into an evaluations set.

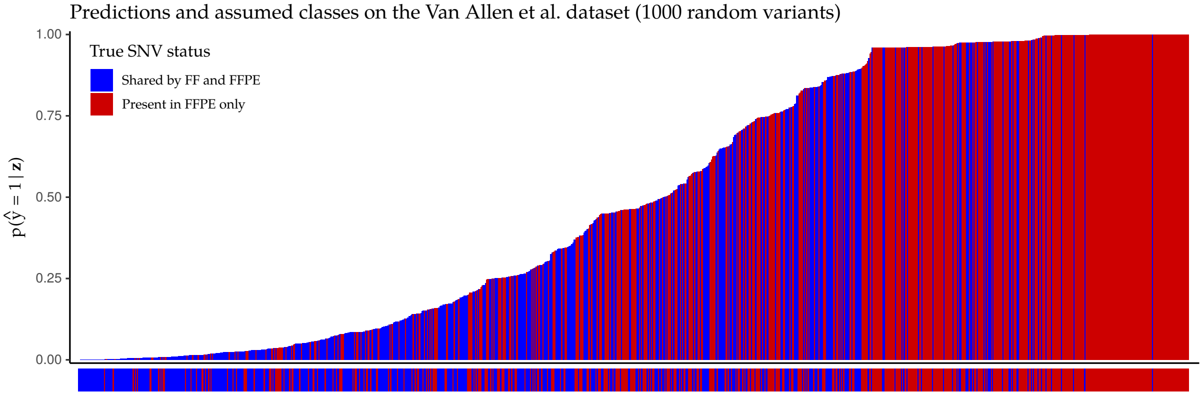

**Suppl.Fig. 41:** Predicted $P$(variant is artifact|attributes) SOBDetector probabilites of 1000 randomly samples variants from the Van Allen et al. dataset.

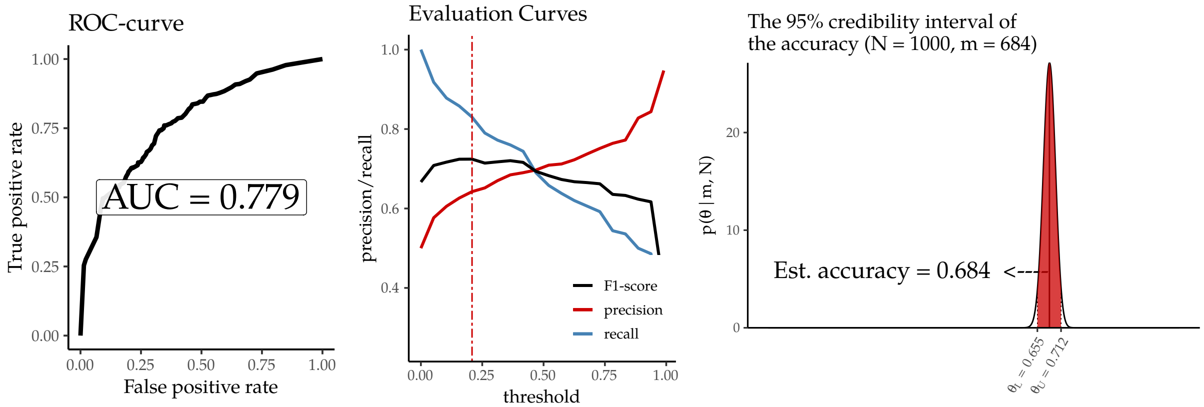

**Suppl.Fig. 42:** Evaluation curves and predicted accuracy - Van Allen et al. dataset.

Using the threshold where the harmonic mean of the recall and precision were at its maximum, the estimated accuracy is 0.684 ± 0.02. **While this estimate is significantly lower than the accuracy measured on the TCGA or on the Oh et al data sets, it is not surprising, as many of the FFPE-only variants are genuine (clonal, low-frequency) mutations.**

##### 2.2.4 Sample-level results

The following figures will show the results of a SOBDetector filtering on a sample-level. The figures contain sorted barplots of the SOBDetector prediction scores, with their corresponding SOB scores, tumor allele frequencies, number of variant supporting reads and coverage at their locus in the tumor. The FilterByOrientation (Mutect2) predictions are also indicated: white color corresponds to predicted genuine mutations, black to likely artifacts. Sadly FilterByOrientation flagged only 6 variants in the entire cohort (including the FFPE normals) as likely FFPE artifact. The two approaches could not be compared therefore.

#### Patient E00565:

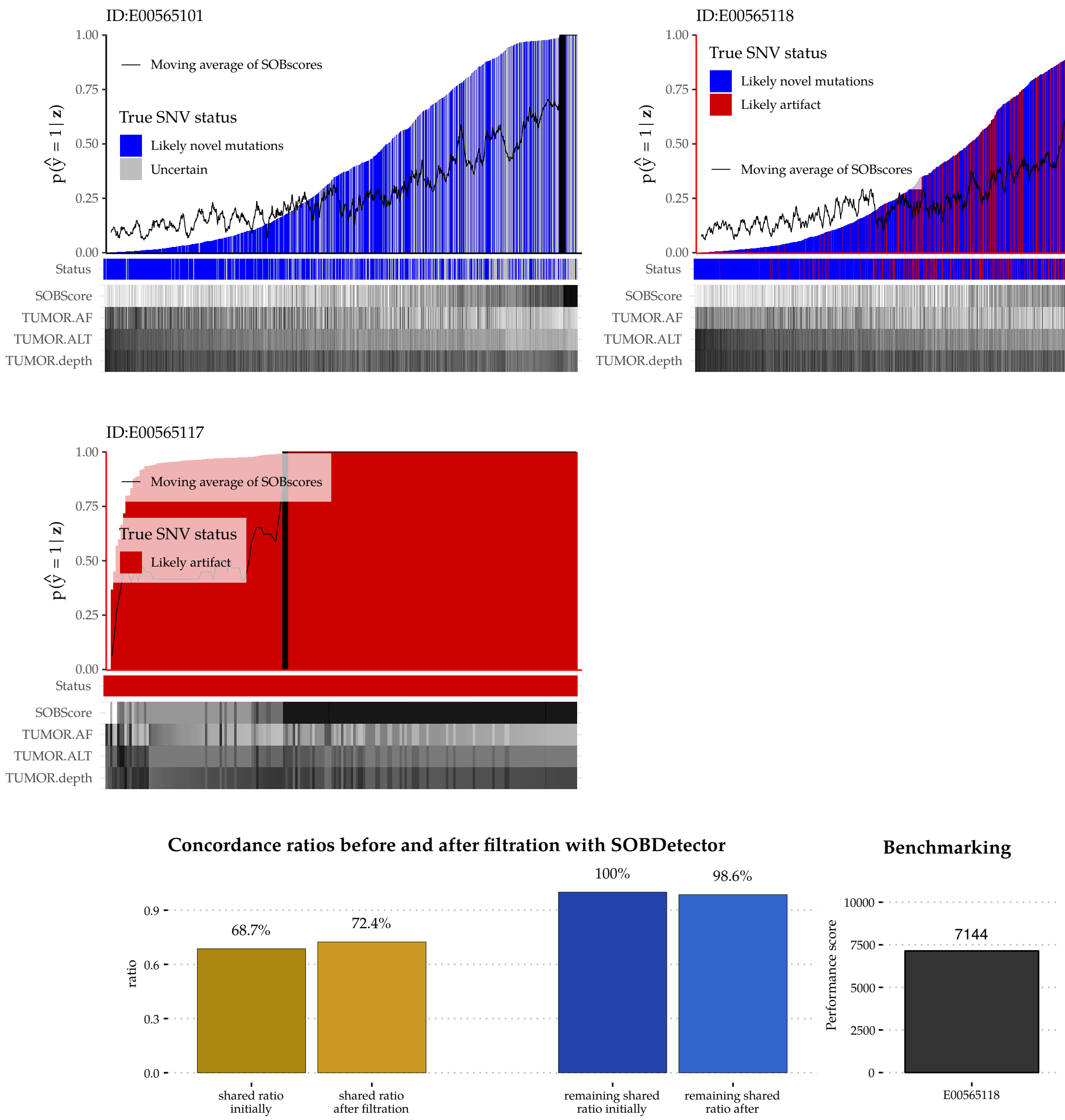

**Suppl.Fig. 43:** Sample-level results of Patient E00565.

#### Patient E00623:

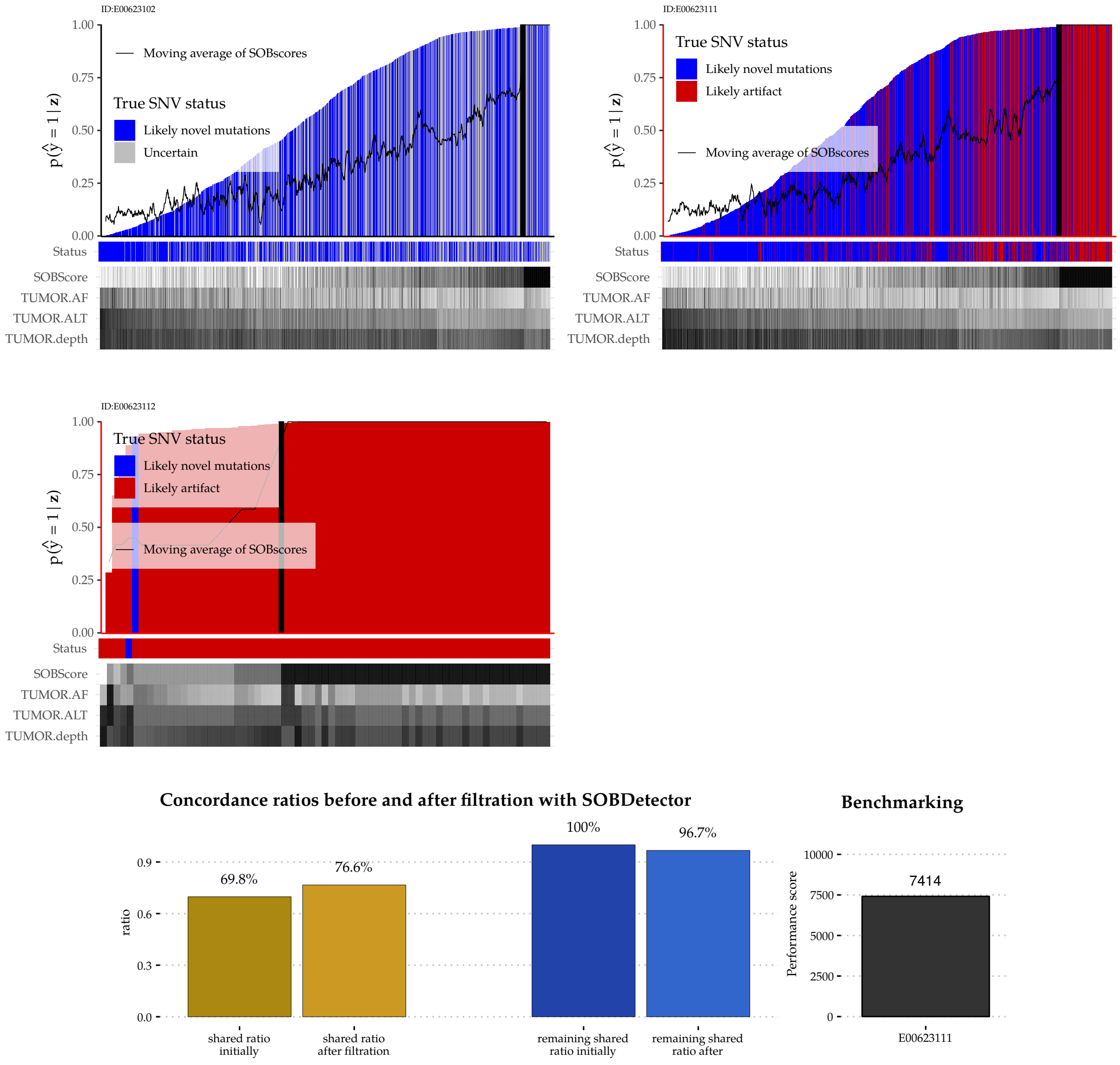

**Suppl.Fig. 44:** Sample-level results of Patient E00623.

#### Patient E00905:

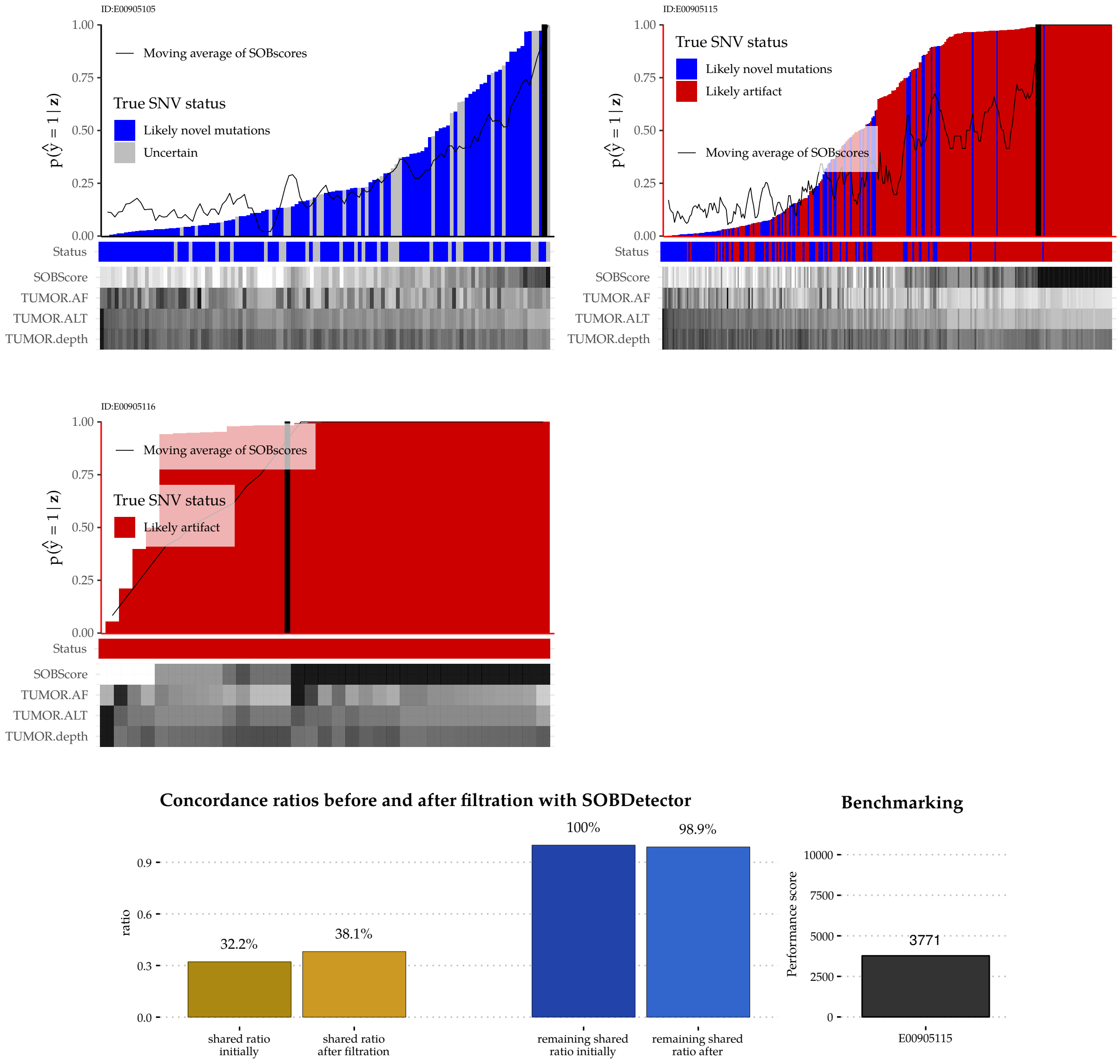

**Suppl.Fig. 45:** Sample-level results of Patient E00905.

#### PATIENT E00905:

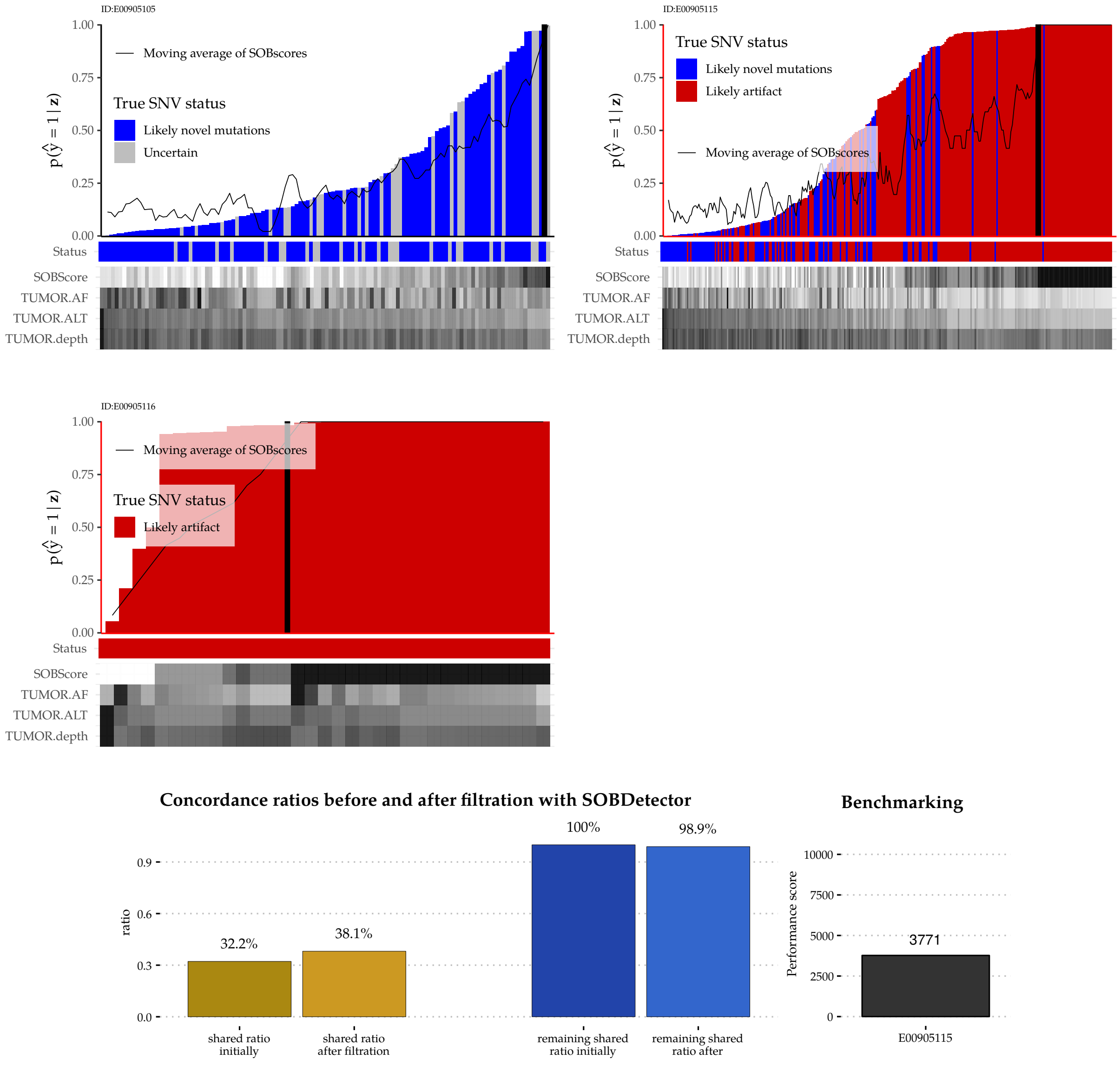

**Suppl.Fig. 46:** Sample-level results of Patient E00905.

#### Patient E00934:

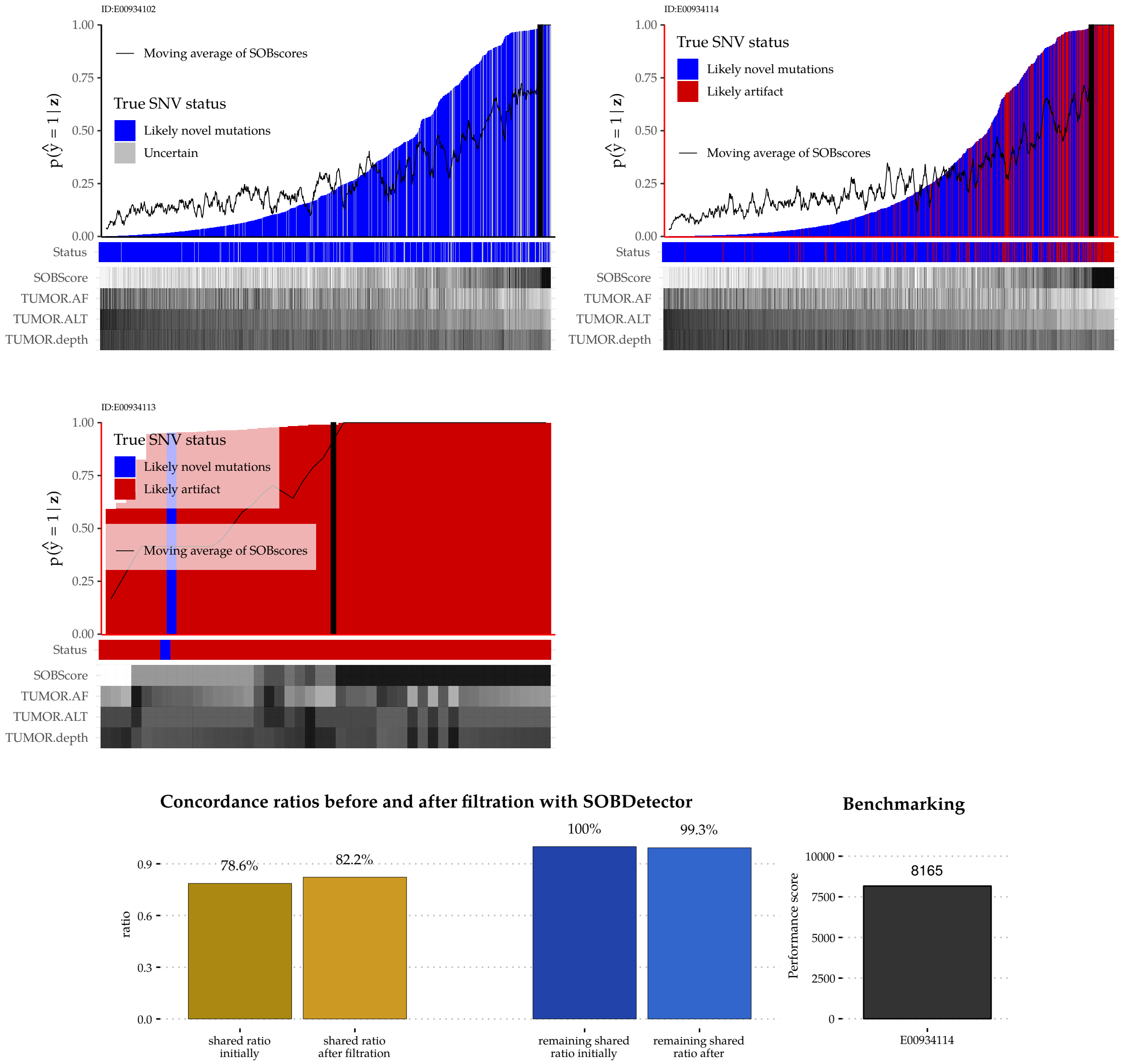

**Suppl.Fig. 47:** Sample-level results of Patient E00934.

PATIENT E01047:

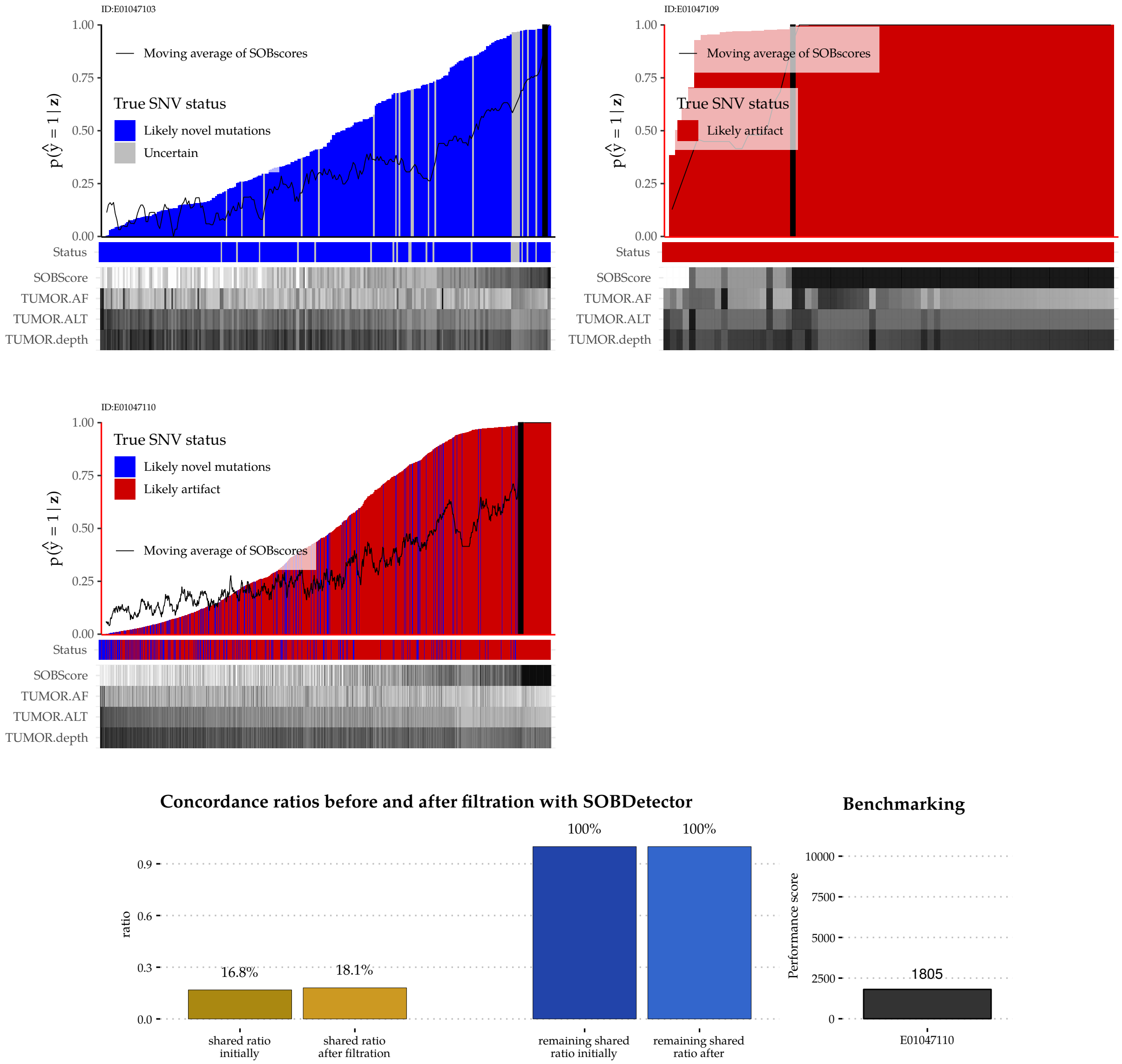

**Suppl.Fig. 48:** Sample-level results of Patient E01047.

##### Patient E01086:

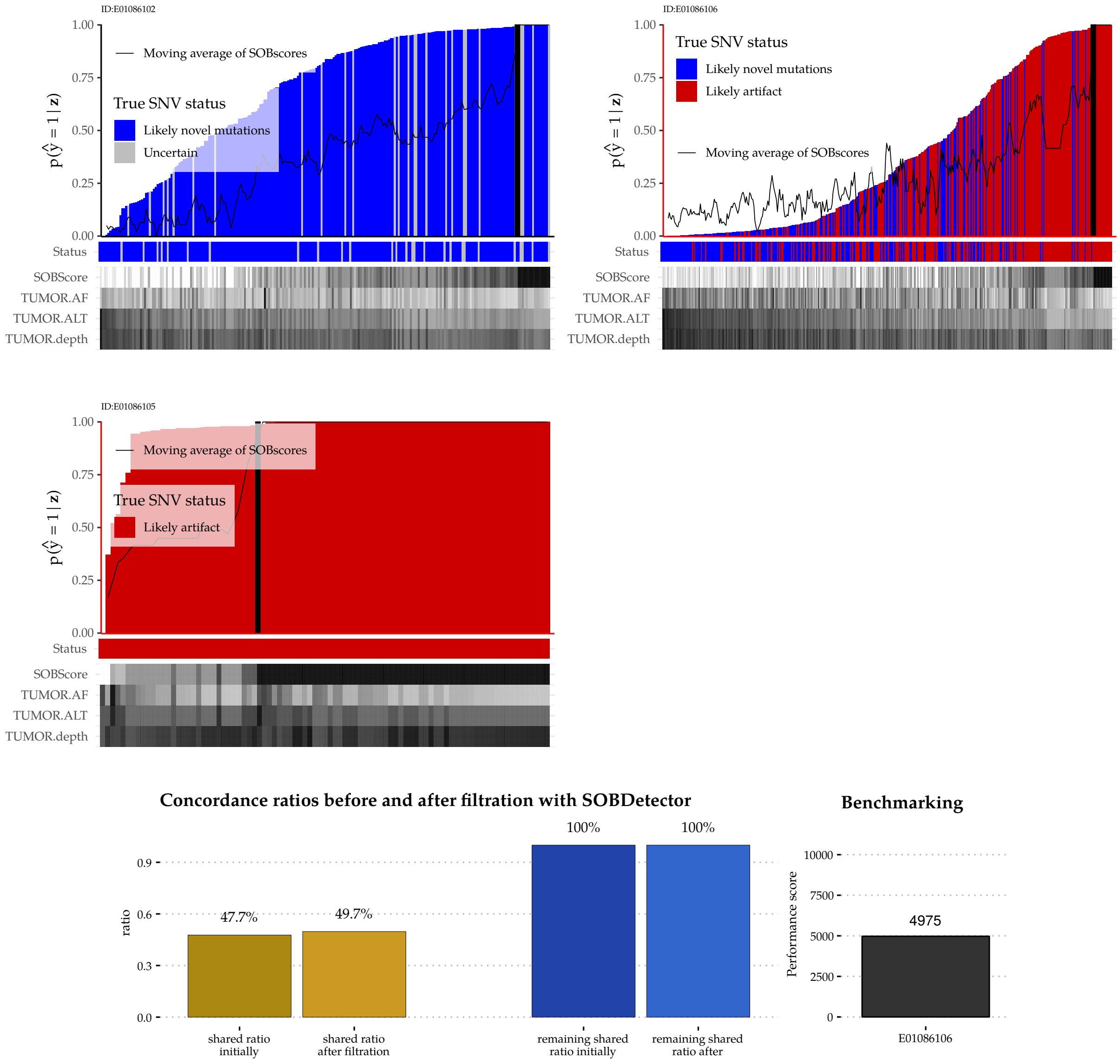

**Suppl.Fig. 49:** Sample-level results of Patient E01086.

#### Patient E01147:

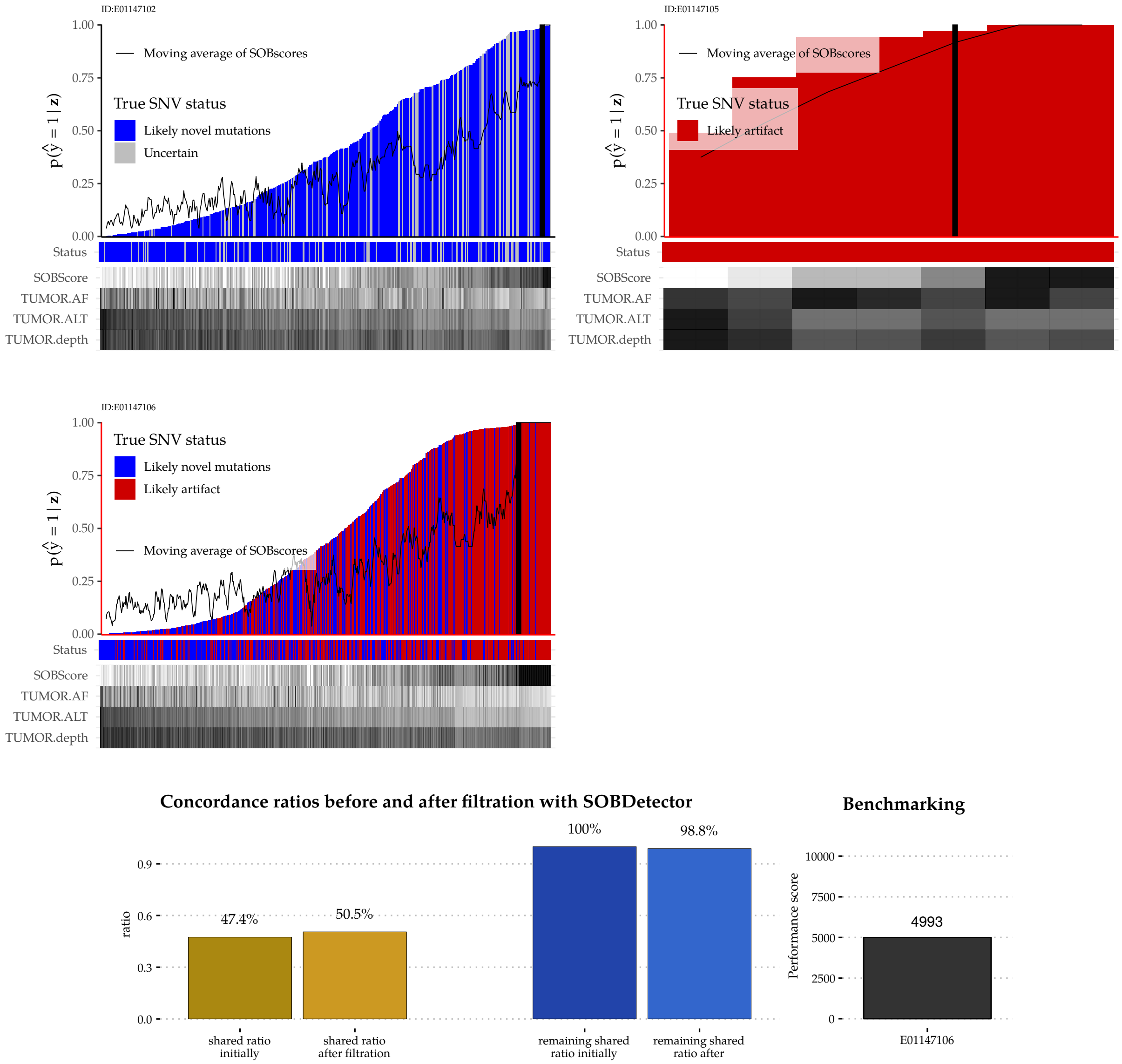

**Suppl.Fig. 50:** Sample-level results of Patient E01147.

#### Patient E01166:

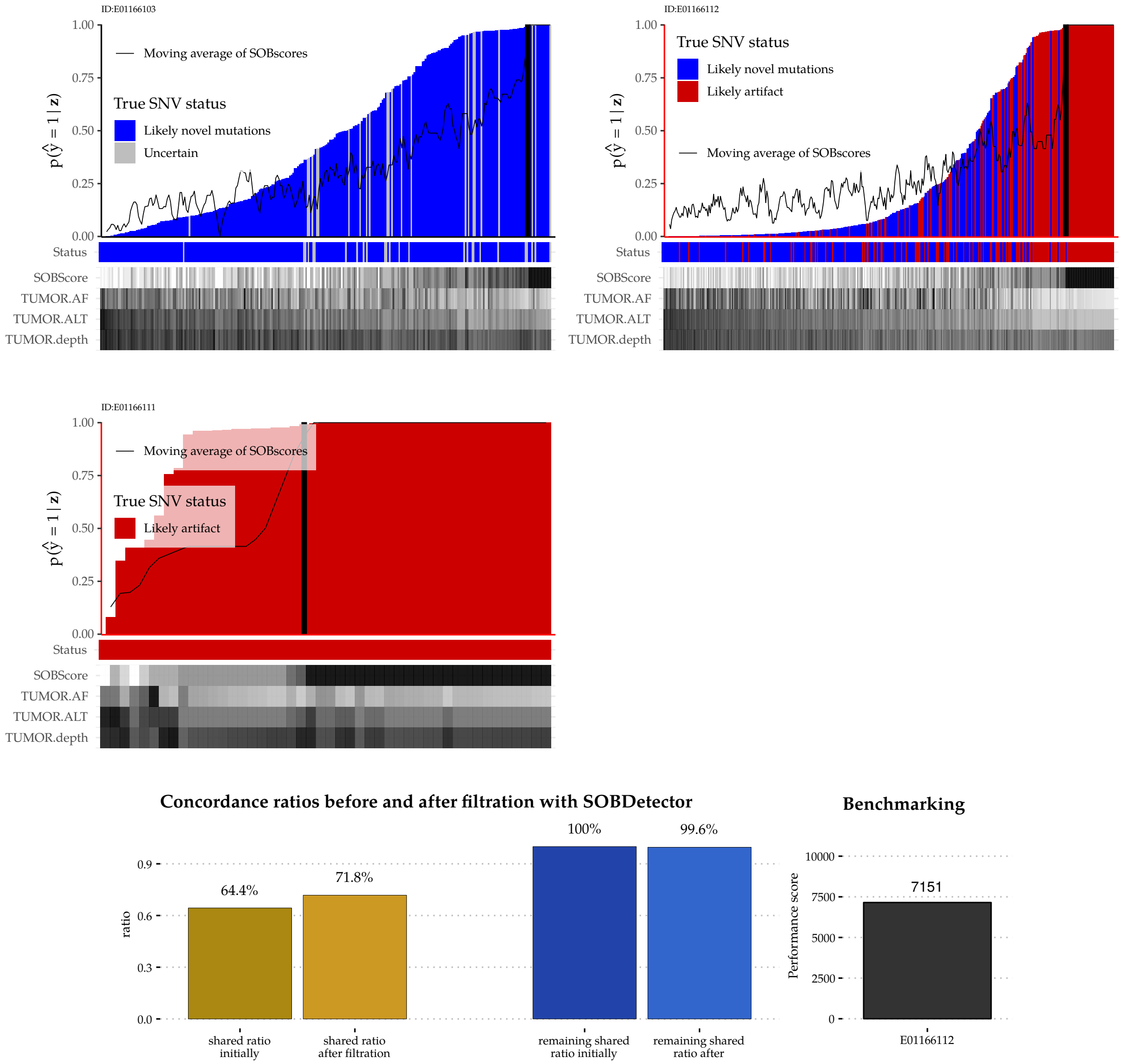

**Suppl.Fig. 51:** Sample-level results of Patient E01166.

#### Patient E01217

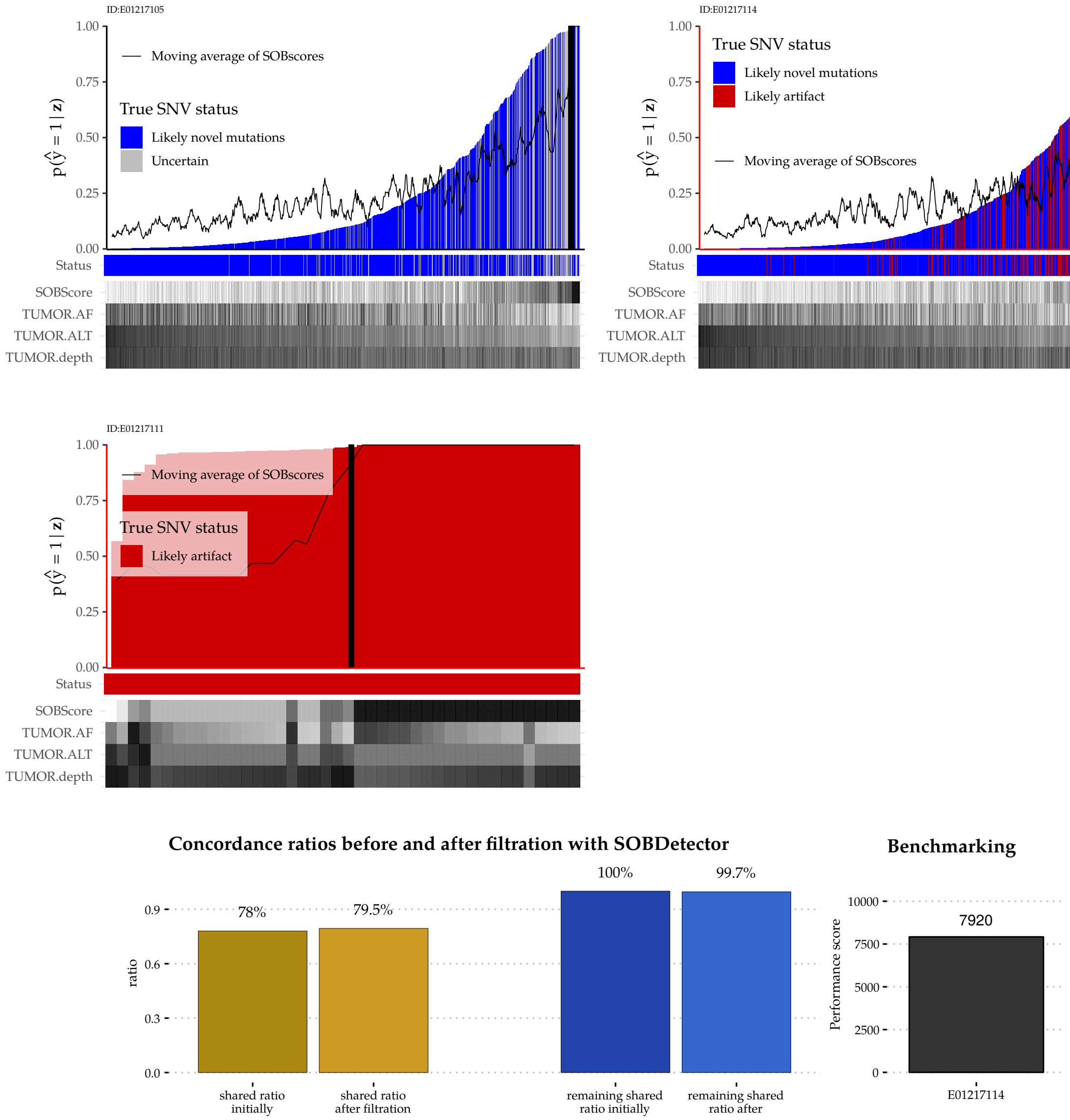

**Suppl.Fig. 52:** Sample-level results of Patient E01217.

#### Patient E01278

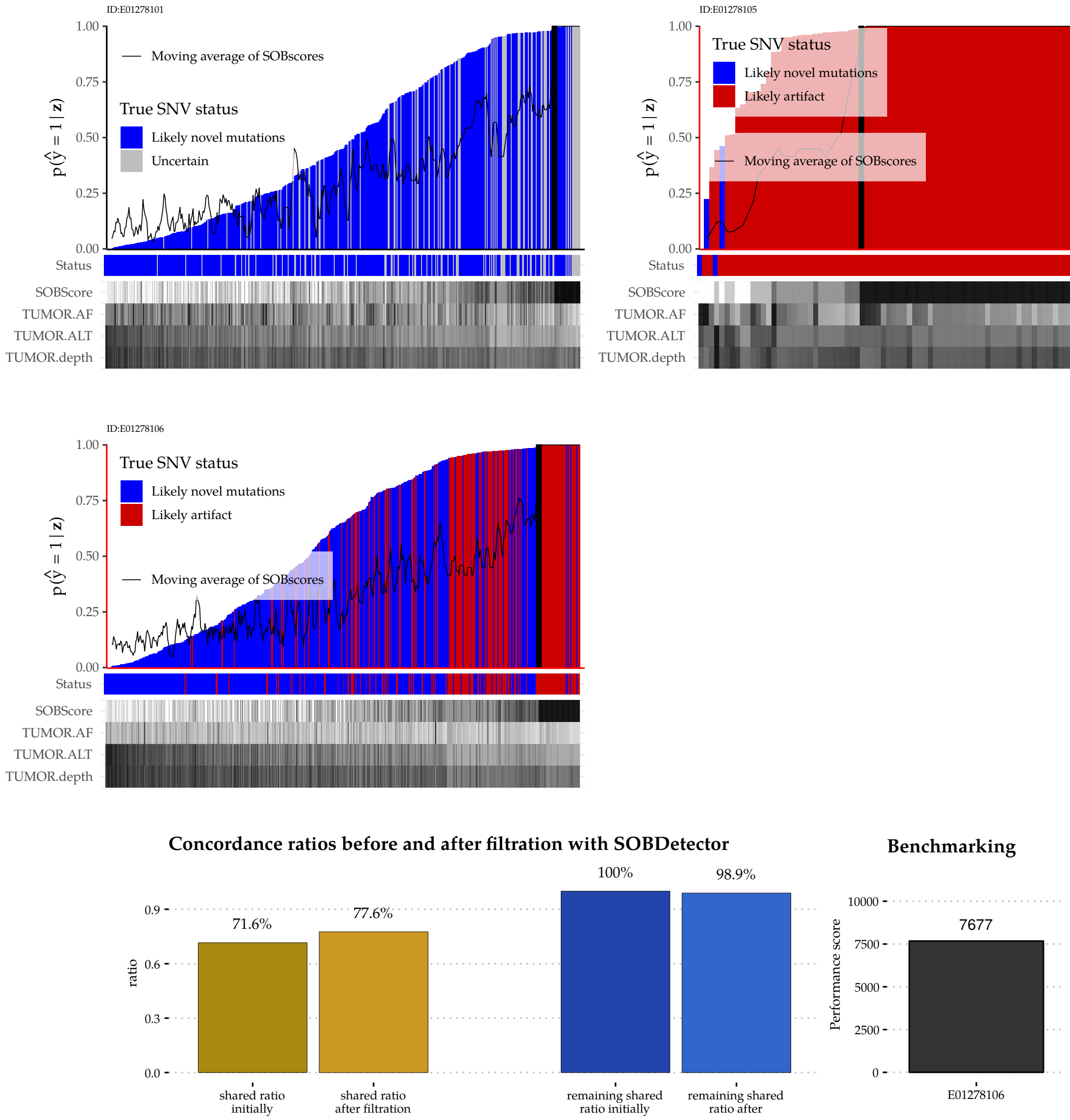

**Suppl.Fig. 53:** Sample-level results of Patient E01278.

##### Patient E01317

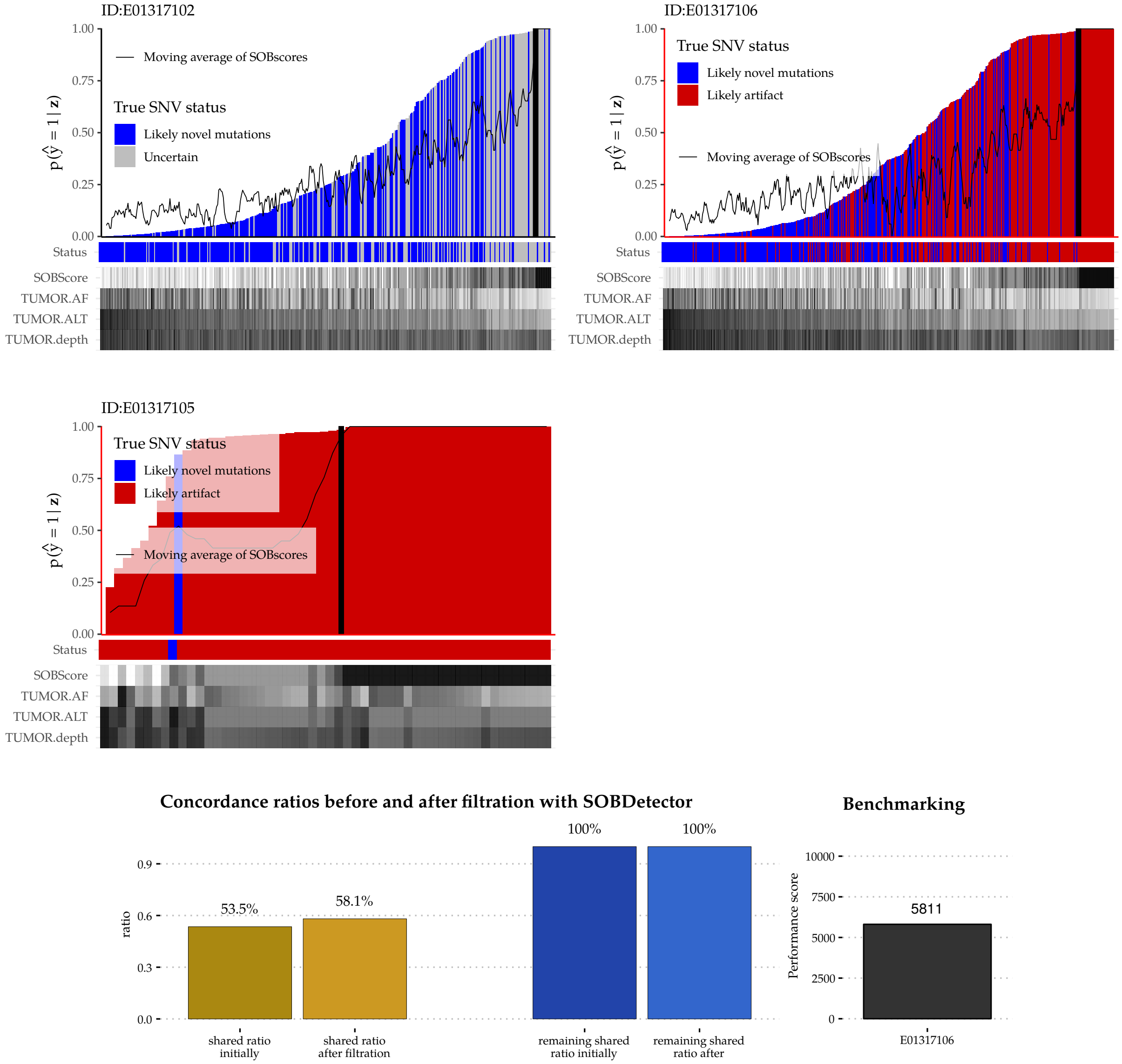

**Suppl.Fig. 54:** Sample-level results of Patient E01317.

##### 2.2.5 Conclusion

Due to higher coverage of the FFPE samples, many FFPE-only variants were low frequency, but genuine mutations. The validation curves therefore suggested a decreased performance from the part of SOBDetector, however the sample-wise figures clearly indicates that the tool managed to isolate a group of variants in every FFPE sample, that exhibited the orientation bias. As a result of this, the concordance has increased in every single case. Approximately half of the false positive mutation calls in the FFPE normals were marked by the tool. The other half of the mutations might be the results of other error sources, such as the sequencer itself. (The 11 normal FF samples probably were not adequate enough to remove all systematic errors.)
